# Common variation of cell abundances and gene expression in the human striatum

**DOI:** 10.64898/2026.03.20.713160

**Authors:** Steven C. Burger, James Nemesh, Olivia Yoo, Ezra Muratoglu, Charles Vanderburg, Jiayi Yuan, Khalid Shakir, Curtis J. Mello, Haoyuan Gao, Nirmala A. Rayan, Emily Finn, Julianna Milidantri, Abir Mohsin, Kathleen Kim, Sadie Drouin, Nikita Budnik, Melissa Goldman, Stanislav Bratchikov, Haley Fritch, Giulio Genovese, Marina Hogan, Olivia Catalini, Seva Kashin, Nicole Rockweiler, Alec Wysoker, Lauren Macaisa, Lucas Reese, Katelyn Flowers, Andrew W. Kraft, Stephen J. Fleming, Madelynn Coe, Riyaan Gunaratne, Liv Spina, Geneva Fischer, Catherine Crombie, Nolan Kamitaki, Evan Z. Macosko, Kiku Ichihara, Steven A. McCarroll

## Abstract

The cellular and molecular bases of inter-individual variation in the human brain are not yet understood, particularly beyond the neocortex. Here we describe cellular and gene-expression variation in the striatum and prefrontal cortex from single-nucleus RNA-seq analysis of 3.9 million nuclei from 178 adult brain donors. People with more astrocytes, microglia, or oligodendrocyte precursor cells (OPCs) in any one brain region tended to have this property in all gray matter regions sampled. OPCs proportions declined by 40% across ages 30–80 in cortex and striatum. Age-associated gene-expression changes differed substantially across cell types. Donor age was predicted from RNA-expression patterns with a median absolute error of 6.6 years across cell types. Common genetic variants were associated with expression of nearly 10,000 genes; these effects were mostly cell-type-specific and distinguished striatal from cortical neurons. These open-access data provide a foundation for future studies of human brain variation and vulnerabilities.

## Introduction

Human brains exhibit abundant biological variation — in size, in activity patterns, in behavioral proclivities, and in vulnerabilities to illness. Humans harbor much genetic variation and live in diverse environments that shape brain development and function in many ways.

The striatum, as the principal input nucleus of the basal ganglia, is a key brain region in which biological variation can influence brain function. Striatal activity contributes to behavioral flexibility, reinforcement learning, decision making, motivation, and motor control^1–7^. The striatum is also influenced by neuromodulatory systems that could reflect and further shape inter-individual differences in molecular state and cellular activity across striatal circuits. These networks are involved in many neurological and psychiatric disorders, including obsessive-compulsive disorder^8–10^ and pronounced neurodegeneration in Huntington’s disease^11,12^.

Much of our current cellular and molecular understanding of the striatum is derived from rodent and non-human primate models^13^, which have provided foundational insights into basal ganglia circuitry and cellular organization^14–19^. Considerable work has been done to identify and characterize the striatum’s cell types^20–24^, but we know less about how these cells vary from person to person.

Single-cell genomics generates detailed cellular and molecular profiles of any brain region; however, it has historically been challenging to determine which aspects of inter-individual variation are biologically real and which are technical. We sought to address these challenges using two aspects of experimental design. First, we analyzed several brain regions in each individual sampled, identifying cellular and molecular features that were consistent properties of an individual across their brain areas. Second, we performed all experiments in a well-controlled “village” format^25^, in which tissue samples from ∼20 donors were processed together and each nucleus later reassigned to its donor using transcribed genetic variants.

Here, and in concert with broader efforts by the BRAIN Initiative Cell Atlas Network (BICAN) consortium, we applied this approach to deeply analyze the main structures of the striatum – the caudate (CaH), putamen (Pu), and nucleus accumbens (NAC) – which share major cell types in common, but differ in functional connectivity^26,27^, gene expression^28,29^, and disease vulnerability^12,30–33^. We also analyzed the internal capsule (ic), a white matter tract between the caudate and putamen, and dorsolateral prefrontal cortex (DFC), an extensively characterized cortical area, to help recognize properties that extend beyond the striatum.

We created an open-access snRNA-seq resource comprising 3.9 million nuclei from 178 adult human brain donors. Many cellular features varied quantitatively among individuals and recurred across brain regions or associated with age and common genetic variation, supporting their biological basis. We provide a web portal (RRID:SCR_028065) for querying these data and informing the design of future tissue-based studies of human neurobiology and disease.

## Results

### A single-cell resource of inter-individual variation in the adult human striatum

We analyzed brain tissues that had been donated at the end of life by 178 persons. These included a core donor set (n = 147) and additional donors (n = 31) for whom one or more clinical or neuropathological findings became apparent in the course of the project (see **Methods**); for all results, we specify whether the core or extended (core + additional) set of donors were used (see also **Table S1,S2**). Donors were selected based on family consent to open data access, sample characterization by brain bank staff, assessments of tissue quality (see **Methods**), and availability of tissues from the regions of interest. The donor age distribution spanned from 27 to 90+ years old (median = 62; precise ages > 90 were unavailable to protect donor identity), and included 111 male and 67 female donors (**Table S2; Figure S1A-C**).

Variation between individuals is more subtle than the categorical differences that distinguish cell types and species from one another. To better recognize and measure such variation, we implemented a “nuclei village” experimental framework^25^. Systematic dissections of the striatum separated caudate, putamen, nucleus accumbens, and internal capsule, in alignment with the neuroanatomical structures defined in the Harmonized Ontology of Mammalian Brain Anatomy (HOMBA; RRID:SCR_027628) (see **Methods**, **Figure S1D,E**). Tissues from ∼20 donors (always from the same subregion) were combined into donor pools (“villages”). Each village was processed as a single sample through nuclei isolation, droplet encapsulation, cDNA barcoding, library preparation, and sequencing (**Figure 1A**). Consequently, any “batch” effects introduced during any of these experimental steps were shared by all of the donors in a village, making it easy to recognize them by using village as a covariate in analyses (see **Methods**). For each single-nucleus RNA profile, donor identity was recognized using combinations of thousands of transcribed sequence variants on RNA transcripts, using a computational approach we have previously described (Dropulation^34^; RRID:SCR_018142).

**Figure 1.**
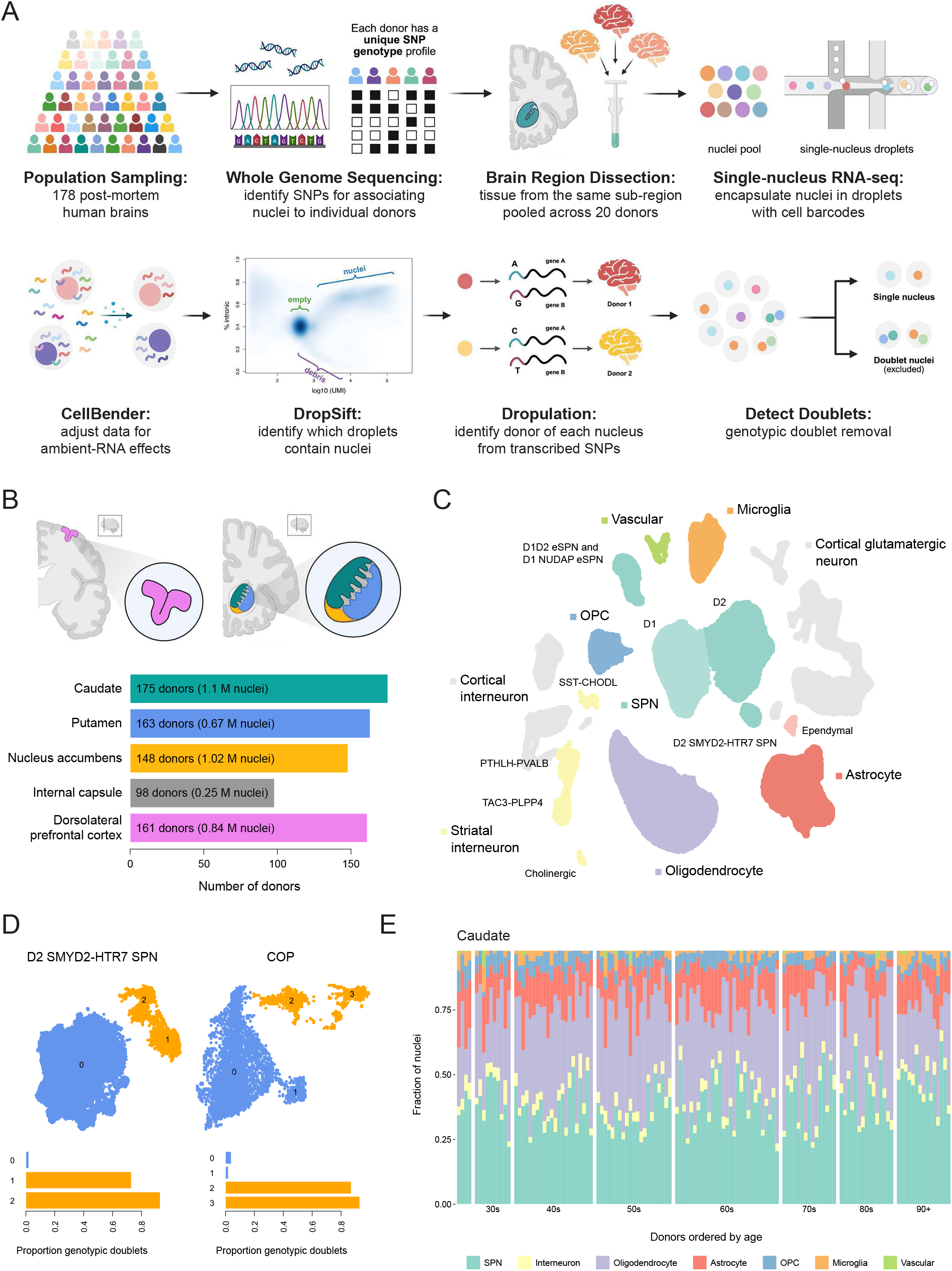
A single-cell resource for recognizing inter-individual variation in the adult human striatum. **A.** Schematic of data generation and processing. Top: multi-donor (“village”) snRNA-seq experimental workflow. Bottom: computational data processing workflow. Diagrams created with BioRender (RRID:SCR_018361). **B.** Brain regions sampled. Left: dorsolateral prefrontal cortex (pink). Right: caudate (green), putamen (blue), nucleus accumbens (yellow), and internal capsule (gray). Diagrams created with BioRender. **C.** UMAP of all 3.9M nuclei assigned to a single donor and cell type. **D.** Genotypic (two-donor) doublet calls identified by Dropulation help confidently identify clusters of cell types that resemble mixtures of other cell types. Left: UMAP of D2 SMYD2–HTR7 SPN; clusters 1 and 2 contain high proportions of genotypic doublets (and are excluded from downstream analyses) compared to cluster 0. Right: UMAP of committed oligodendrocyte precursor (COP); clusters 2 and 3 have high proportions of genotypic doublets (and are excluded from downstream analyses). **E.** Stacked bar plot shows representation of major cell types among the caudate samples from 131 donors. Donors are ordered from left to right by ascending age.

Using this approach, single-nucleus RNA-seq (snRNA-seq) data were generated from caudate head (n = 175), putamen (n = 163), nucleus accumbens (n = 148), internal capsule (n = 98), and dorsolateral prefrontal cortex (n = 161), with multiple regions sampled from the same donors (**Figure 1B, S1F**).

We first performed unsupervised clustering of single-nucleus RNA-expression profiles and manually annotated the resulting clusters according to the consensus basal ganglia taxonomy (**Figure S2**). Dropulation enables the identification of “genotypic” doublets containing pairs of nuclei from two donors, and we then used the expression profiles of these genotypic doublets to identify remaining profiles involving pairs of nuclei from the same donor (**Figure 1D**; see **Methods**). This analysis recognized, in each of the donors, all expected neuronal and non-neuronal cell classes across the sampled regions, including the diverse populations of striatal spiny projection neurons (SPNs, also known as medium spiny neurons or MSNs) (**Figures 1C,E** and **S2**).

After quality filtering and doublet removal, the final dataset comprised 3.9 million nuclei. We next examined inter-individual variation in each cell type’s abundance across brain regions and its associations with age and sex, followed by the effects of common genetic variation on gene expression in each striatal and cortical cell population.

### Glial cell abundances vary from person to person

The relative abundances of cell types varied substantially across brain donors (**Figure 2A**). This variability was also observed in a subset of donors using other experimental methods including bulk RNA-seq and *in situ* hybridization (see **Methods, Figure S3**). Sampling multiple brain regions from the same large panel of donors allowed us to test whether this variation reflects biologically meaningful, donor-level features that are shared across brain regions, or instead arises from localized or experimental factors. (To minimize effects of clinically apparent conditions or those evident by conventional neuropathological analysis, we used the core donor set for these analyses.)

**Figure 2.**
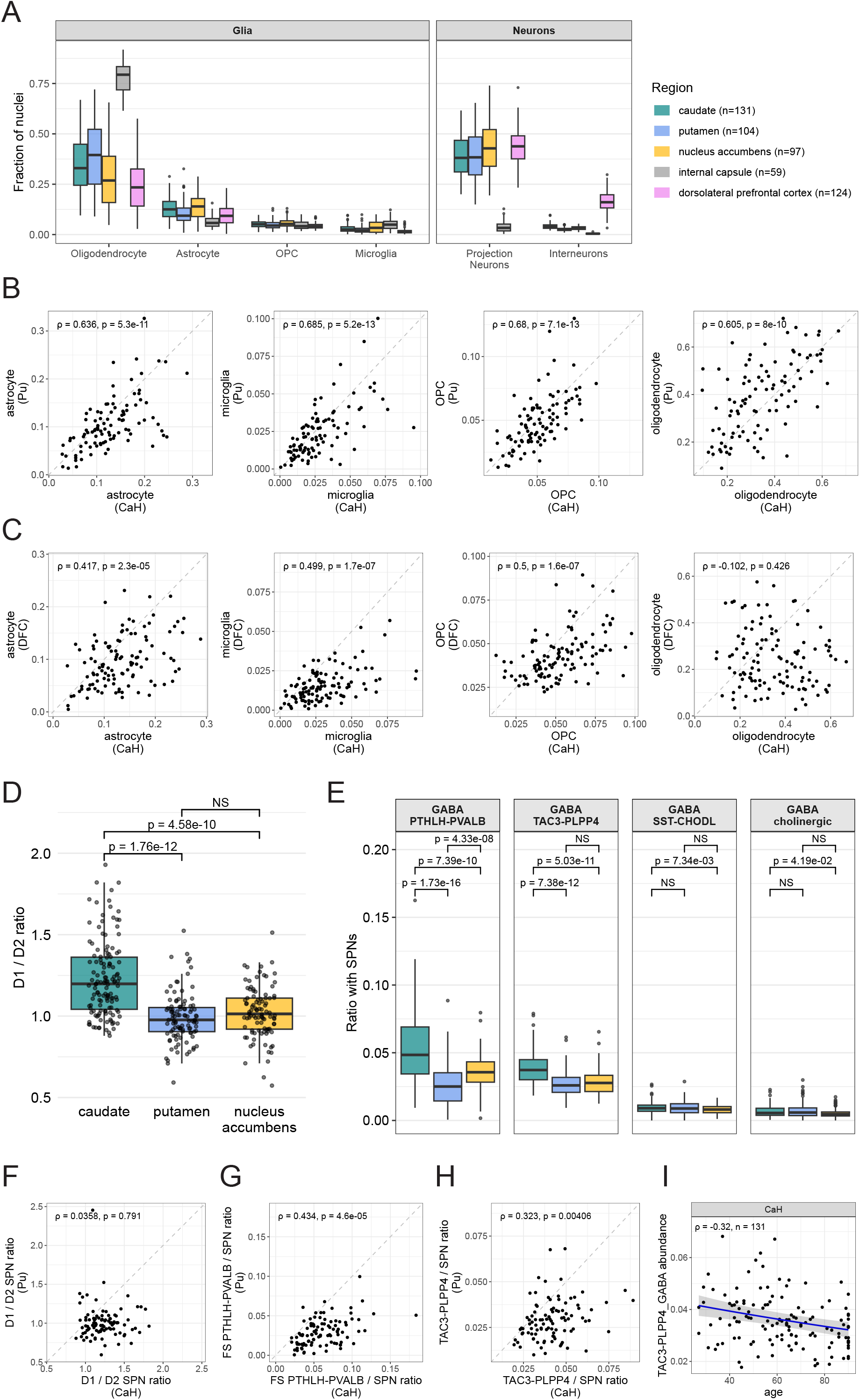
Variation in cell type abundances across individuals and between brain regions. (Extended analyses are in **Figures S4, 6, 7**). **A.** Representation of major cell types in the brain regions sampled. Boxes indicate the interquartile range across donors and whiskers extend to the most extreme points within 1.5x the interquartile range. **B.** Representation of glial cell type abundances (as a fraction of all nuclei sampled) in the caudate and putamen of 98 donors. Spearman correlation coefficient and corrected p-values are shown (Benjamini–Hochberg (BH) procedure, n = 498 tests; see Methods). **C.** Same as **B**, for 114 donors sampled in caudate and DFC. **D.** D1/D2 SPN ratio in CaH (n=131 donors), Pu (n=104 donors) and NAC (n=97 donors); nominal p-values are from the paired Wilcoxon signed-rank test. **E**. Quantitative representation of striatal interneuron subtypes relative to the number of SPNs in each striatal region. Statistical significance was assessed using the paired Wilcoxon signed-rank test and corrected for multiple comparisons (BH procedure, 12 tests). **F.** D1/D2 SPN ratios in CaH and Pu from 98 donors. Spearman rank correlation coefficient and corrected p-value shown (BH procedure, n = 498 tests). **G.** Representation of GABAergic PTHLH–PVALB interneuron abundances (relative to SPNs) in the caudate and putamen of 98 donors. Spearman correlation coefficient and corrected p-values are shown (BH procedure, n = 498 tests). **H.** Same as **G**, but for GABAergic TAC3–PLPP4 interneurons. **I.** TAC3–PLPP4 interneuron abundance declines with age (OR = 0.959, corresponding to a loss of approximately 4% per decade; BH-adjusted p-value = 0.003; n = 54 tests; **Table S5**). Each point represents a donor; TAC3 abundance (y-axis) is represented as a composition with SPNs (TAC3 / TAC3 + SPNs); the blue line indicates beta-binomial fit with 95% confidence intervals shown as gray ribbons. Spearman correlation coefficient and number of donors shown.

We found that abundances of the major glial cell types (astrocytes, microglia, OPCs, and oligodendrocytes) were strongly correlated from one striatal region to another: donors for whom the abundance of one cell class was high in the caudate tended to have this property in the putamen and nucleus accumbens as well (**Figure 2B, S4A** and **Table S4**), and to a lesser extent in the internal capsule (**Figure S4B**). These relationships arose from the entire set of donors, as opposed to a few outliers, and were highly significant even when assessed with non-parametric statistical tests (**Figure 2B, S4A**).

These relationships also extended beyond the striatum: the tendency of the individual donors to have relatively high or low abundances of astrocytes, microglia, or OPCs was also apparent in their DFC (**Figure 2C, S4C** and **Table S4**). These correlations were similarly strong when the abundance of each cell type was calculated relative to the number of neurons rather than the total number of nuclei (to prevent the glial cell types from affecting one another’s abundance estimates; **Figure S4D-F**). These correlations also persisted among all gray matter region comparisons after accounting for postmortem interval, RNA quality score, and library size – covariates potentially relating to tissue quality (see **Methods, Figure S5**). The only exception to this striatal-cortical correlation pattern involved oligodendrocytes (**Figure 2C, S4C**). These results suggest that at least some of the substantial inter-individual variation in glial abundance may reflect donor-level biological differences that extend across brain regions.

### Neuronal proportions differ among striatal regions

Sampling from a large number of donors in multiple brain regions made it possible to recognize quantitative features of striatal subregions that consistently differed from each other, in many different people (using the core donor set). In order to minimize technical noise arising from differential sampling of tissue micro-environments, we measured the abundances of neuronal types relative to one another.

SPNs are classically categorized based on their expression of either D1 or D2 dopamine receptors^35^. D1 and D2 SPNs are further distributed across two anatomical compartments in the striatum: the matrix and striosomes^36–38^ (**Figure S6A,B**). Striosome SPN proportion was similar in caudate and putamen, representing on average 17% of SPNs (**Figure S6C**). The observed ratio of D1 to D2 SPNs was 20% higher in caudate (mean 1.23) than in the putamen (mean 1.00) and nucleus accumbens (mean 1.02) (**Figure 2D**), and was consistent across both matrix and striosome SPNs (**Figure S6D**) and across donors (**Figure S6E**). This aligns with findings from mouse immunofluorescence studies^39,40^ and Slide-tags spatial transcriptomics analyses of the human striatum^41^.

SPNs are surrounded and modulated by a heterogeneous set of GABAergic interneurons with distinct structural and electrophysiological properties^42–44^. The two most abundant striatal interneuron types are the fast-spiking PTHLH–PVALB and the TAC3–PLPP4 types, the latter originally discovered in primate striatum by snRNA-seq^17^ and recently detected in some additional mammalian species^45^. PTHLH–PVALB interneurons were most numerous in caudate, followed by nucleus accumbens and then putamen, forming a medial-to-lateral gradient (**Figure 2E**). TAC3–PLPP4 interneurons were also most numerous in caudate but showed similar abundances in putamen and nucleus accumbens (**Figure 2E**).

We also looked for evidence of inter-individual variation in neuron abundances. Across individuals, the ratio of D1 to D2 SPNs was not significantly correlated across striatal regions (**Figure 2F, S6E**). However, the abundances of interneuron subtypes exhibited strong evidence of inter-individual variation, in patterns that appeared consistently across the striatum. PTHLH– PVALB and TAC3–PLPP4 interneurons showed strong correlations in abundance across all pairs of striatal regions (**Figure 2G,H, S7A-C**). Abundances of different types of interneurons were also correlated with each other: donors with higher abundances of PTHLH–PVALB interneurons also had higher abundances of TAC3–PLPP4 interneurons (**Figure S7D**).

Among interneurons (relative to SPNs), the abundance of TAC3–PLPP4 interneurons showed a modest age-associated decline, corresponding to approximately 18.6% between ages 30 and 80, and observable in all three striatal gray-matter areas sampled (OR = 0.959 per decade, corresponding to a loss of about 4% per decade; BH-adjusted p-value = 0.003; n = 54 tests; **Figures 2I**, and **S7E**, **Table S5**). Because TAC3–PLPP4 interneurons are sparse and large independent striatal datasets (with sufficiently deep sampling) are limited, we suggest that this association be considered preliminary until replicated. In addition, cortical SST interneurons in the DFC exhibited steep age-associated attrition (**Figure S7F**), an effect that can be seen (for this abundant interneuron population) in many snRNA-seq datasets ^25,46,47^. (Additional analytical details and (generally negative) analyses looking for potential sex effects are in **Table S5**).

### OPC proportions decline with age across brain regions

We next asked whether abundances of any glial cell types were shaped by age or sex, again using the core donor set. Broad cell classes were quantified relative to all nuclei, while analyses of neuronal subtypes considered them in relation to other neuronal types, to reduce sensitivity to glial variation and tissue microdissection (e.g. white-matter tracts). Among well-ascertained cell types (median > 1% of all nuclei across samples), and after correcting for multiple hypothesis testing, only OPC abundance was significantly associated with age (OR = 0.908, corresponding to a loss of about 9% per decade; BH-adjusted p-value = 5.25e-12; n = 54 tests; **Table S5**, **Figure 3A, S8A,B**). This decline was observed in all brain regions sampled. No glial cell type’s abundance showed a significant association with sex (**Table S5**).

**Figure 3.**
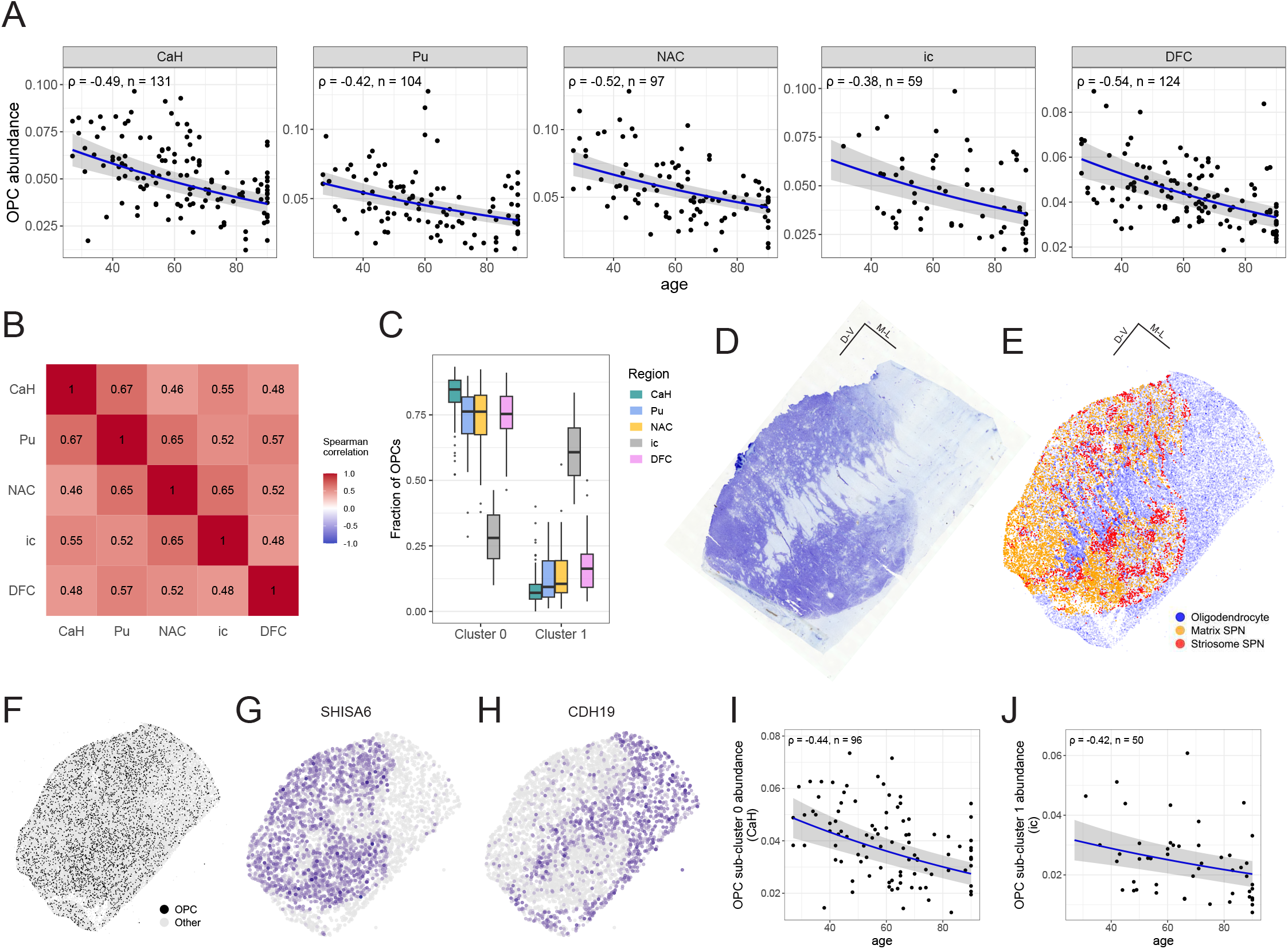
Decline in OPC populations in multiple brain regions with advancing age. (Extended analyses are in **Figure S9**). **A.** Loss of OPCs with advancing age in five brain regions; measured as a fraction of all nuclei per sample (OR = 0.908, corresponding to a loss of approximately 9% per decade; BH-adjusted p-value = 5.25e−12; n = 54 tests; **Table S5**). Each point is a donor. Spearman rank correlations and number of donors are shown for each region. Lines represent beta-binomial fit (95% confidence intervals shown with ribbons). **B.** Cross-region correlations of age-adjusted OPC fractions (observed minus age-expected fractions). **C.** Fraction of all OPCs in cluster 0 and cluster 1, by region. **D**. Nissl stain of the striatum from one donor. **E.** Spatial transcriptomics data of an adjacent section (same donor), generated with Slide-tags. Points are oligodendrocytes (blue), matrix SPNs (orange), or striosome SPNs (red) nuclei. **F.** Spatial distribution of OPCs (black). **G.** OPC expression of SHISA6 (cluster 0 marker) highlights gray matter compartments; color indicates log-normalized expression. **H.** OPC expression of CDH19 (cluster 1 marker) highlights white matter compartments. **I.** OPC cluster 0 abundance (fraction of all nuclei) declines with age in CaH (OR = 0.906, 95% CI = 0.882 – 0.931, BH-adjusted p-value = 1.36e-10; n = 54 tests; **Table S5**). **J.** OPC cluster 1 abundance (as a fraction of all nuclei) declines with age in ic (OR = 0.927, 95% CI = 0.891 – 0.964, BH-adjusted p-value = 8.91e-3; n = 54 tests; **Table S5**).

OPCs are proliferative oligodendrocyte progenitors with additional recognized functions in the adult CNS, including synaptic interactions with neurons^48^. We estimated that OPCs, as a proportion of all nuclei, decline by about 40% between age 30 and age 80 (see **Methods**), and found no evidence that this rate differed among any of the brain regions sampled (see **Methods**). We also found that the strong age-associated decline in OPC abundance persisted after correcting for a variety of potential technical effects such as postmortem interval and RNA quality (see **Methods**). (Loss of OPCs with age has been previously observed in the human neocortex ^46,49–51^.)

The abundance of OPCs could in principle also be affected by individually varying factors beyond chronological age. To assess this, we calculated an OPC fraction residual, the difference between a sample’s measured OPC fraction and the OPC fraction predicted from their age (and other model covariates, see **Methods**). These OPC fraction residuals were positively correlated between all pairs of brain regions (**Figure 3B, S8C**): donors with higher-than-expected OPC abundance for their age in one brain region tended to exhibit similarly elevated OPC abundance in other regions. The consistency across diverse brain regions of both the effect of age and the residual unexplained by age suggests additional donor-level influences beyond chronological age, potentially including genetic, health-related, and environmental factors.

To better characterize OPC loss and OPC diversity ^52,53^ across these brain regions, and recognize what OPC populations were affected, we sub-clustered the OPC gene expression profiles (from ∼155K nuclei), identifying seven transcriptionally distinct clusters (**Figure S9**, see **Methods**). The two most abundant clusters closely resembled OPC populations previously described as “immature” (cluster 0, expressing *SHISA6, SGCZ, GPC6*) and “mature” (cluster 1, expressing *CDH19*, *SEMA3E*, *GRIA4*)^54^. Among genes differentially expressed between these OPC populations, the most distinctive and interpretable signal for cluster 0 was the elevated expression of several genes that encode synaptic proteins (*GABRB2*, *KCNB2*, *SYN3*, *NCALD*, *SHISA6*) and specific transcription factors (*TCF7L1*, *PBX3*, *RFX4*). Cluster 1 was marked by higher levels of *CDH19*, which encodes a classic oligodendrocyte adhesion molecule. OPC clusters 0 and 1 were also distinguished by their expression levels of distinct sets of axon guidance genes (*CNTN5*, *EPHA5*, *ADGRV1* in cluster 0; *SEMA3E*, *PDZRN3*, *DACH2* in cluster 1) and extracellular matrix genes (*COL4A2*, *FBLN2*, *COL19A1* in cluster 0; *ADAMTSL1*, *COL20A1*, *CHRDL1* in cluster 1). These tissue-context-dependent expression profiles are consistent with the idea that OPCs have functions beyond serving as oligodendrocyte progenitors^55^.

Both of these clusters were observed in all brain regions; however, the internal capsule (the only white matter region analyzed) had a far lower proportion of cluster 0 OPCs and greater proportion of cluster 1 OPCs (3.6–7.1 fold; DFC and caudate, respectively) (**Figure 3C, S9C,D**), similar to the pattern of oligodendrocyte enrichment in the internal capsule relative to gray matter regions (**Figure 2A**). To ask whether these clusters might vary in their spatial distributions (e.g., potentially between gray and white matter regions) we examined Slide-tags spatial transcriptomics data of the striatum generated concurrently^41^ (**Figure 3D,E).** OPCs were evenly distributed throughout both the gray and white matter of the striatum (**Figure 3F**), consistent with reports in mice^56^. However, when visualized spatially, marker genes for cluster 0 highlighted gray matter compartments (**Figure 3G, S9E**), while marker genes for cluster 1 highlighted white matter compartments (**Figure 3H, S9F**). The spatial localization of these OPC populations further supports the idea that the fundamental difference between these subsets of OPCs may be tissue context (gray matter vs. white matter).

Across all brain regions sampled, the quantitative presence of both OPCs with a cluster 0 profile (gray matter-enriched; OR = 0.91, corresponding to a decline of about 9% per decade; BH-adjusted p-value = 1.4e-10; n = 54 tests) and OPCs with a cluster 1 profile (white matter-enriched; OR = 0.93, corresponding to a decline of about 7% per decade; BH-adjusted p-value = 8.91e-3; n = 54 tests) declined with age (**Figure 3I,J, S9G,H**), suggesting that this age-associated loss affects OPCs broadly and in diverse tissue microenvironments. Among OPC clusters, cluster 4, corresponding to mitotic OPCs (based on expression of cell-cycle genes such as *MKI67*, *TOP2A*, *CENPF*; see **Methods**) showed the steepest age-associated decline (OR = 0.78, corresponding to a decline of about 21% per decade; BH-adjusted p-value = 2.60e-7; n = 54 tests; **Table S5**). Committed oligodendrocyte precursors (COPs), a more differentiated, lineage-committed precursor state between OPCs and oligodendrocytes, also declined with age at a rate comparable to that observed for OPCs (OR = 0.86, corresponding to a decline of about 14% per decade; BH-adjusted p-value = 0.13e-3; **Table S5**). Together, these results suggest that the age-associated decline in OPC proportions may reflect reduced maintenance or replenishment of the OPC pool, rather than increased differentiation alone.

### Age-associated changes in cell-type-specific RNA expression

For each cell type, gene expression profiles also varied quantitatively among individuals. We used mixed-effects regression models to identify cell-type-specific expression differences associated with age and sex, adjusting for donor and village and for brain region, 10x chemistry, biobank, postmortem interval (PMI), genetic ancestry, and sample-level sequencing/QC covariates (**Table S3**; **Methods**).

The project also provided an opportunity to empirically evaluate two kinds of sample-exclusion criteria: “metadata-driven” exclusions based on reported health circumstances; and “snRNA-seq-data-driven” exclusions based on having highly atypical cell-type proportions or gene expression (see **Methods**). We found that metadata-driven exclusions substantially reduced statistical power without materially affecting estimated age-effect sizes, whereas snRNA-seq-data-driven exclusions often increased statistical power by reducing noise (**Figure S10**). We therefore used the extended donor set (n = 178), applying only snRNA-seq-data-driven exclusions, which removed approximately 5% of available samples (**Table S1,2**).

Sex-associated gene-expression differences were observed across all cell types, with the largest effects involving sex-chromosome genes; significant effects on autosomal genes were also detected but were generally smaller in magnitude (**Figure 4A, S11**).

**Figure 4.**
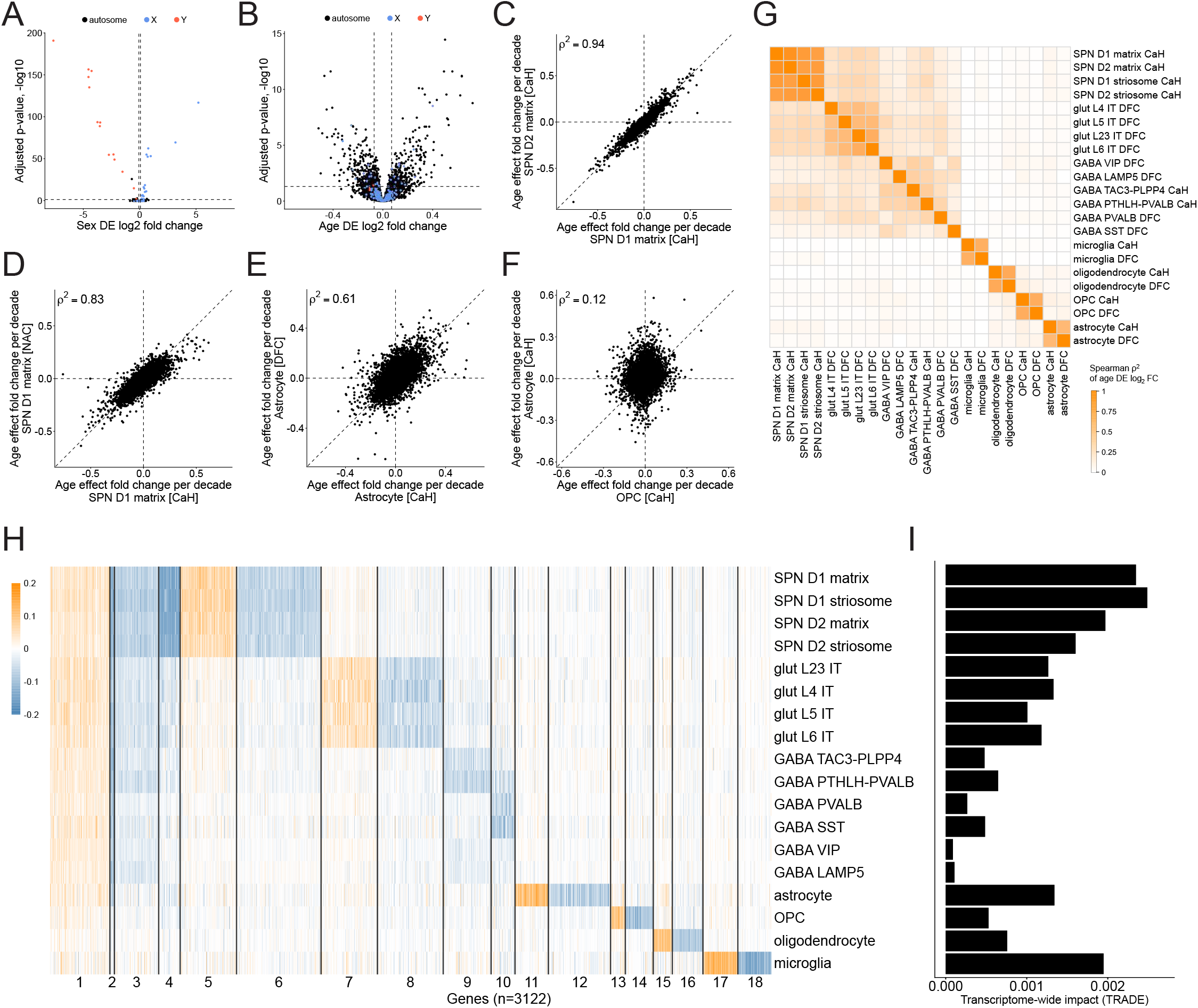
Age-associated changes in gene expression across brain cell types. (Extended analyses are in **Figure S13**.) **A–B.** Volcano plots (points are genes) of (**A**) sex differences in astrocyte gene expression and (**B**) age-associated gene-expression changes in astrocytes; in all panels age log_2_-fold-change is calculated per decade (see **Methods**). **C–F.** Comparison of age-associated gene-expression changes between (**C**) D1 SPNs and D2 SPNs (matrix subtype, sampled in the caudate), (**D**) D1 matrix SPNs sampled in the caudate (x-axis) or the nucleus accumbens (y-axis), (**E**) striatal astrocytes (x-axis) and cortical astrocytes (y-axis), and (**F**) OPCs (x-axis) and astrocytes (y-axis) in the caudate. **G.** Correlation heatmap of age-associated gene-expression changes for each pair of cell types; colors show signed Spearman’s ρ^2^. **H.** Clustering (k-means, see **Methods**) of genes (columns) by their patterns of age-associated expression changes across cell types. **I.** Magnitude of transcriptome-wide age-associated expression changes in each cell type, accounting for power differences (see **Methods**).

In each cell type, donor age was associated with the expression levels of hundreds to thousands of genes (**Figure 4B**). These effects replicated strongly in analyses of available data from the Ling et al. study (**Figure S12**).

Age-associated effects were highly correlated among SPN subtypes (**Figure 4C**), within and across striatal regions (**Figure 4D, S13B**). For glia of all types, age effects were also highly correlated across brain regions (e.g., between striatal and cortical astrocytes) (**Figure 4E, S13A,C**).

In contrast, age affected gene expression in distinct cell types in very different ways, with very different sets of changes in each of the glial cell types, and in striatal (relative to cortical) projection neurons (**Figure 4F-G, S13A**).

To better recognize shared and cell-type-specific patterns of brain aging, we clustered genes by their cell-type-specific age-associated expression changes (**Figure 4H, S13D**, see **Methods**). A small subset of the age-affected genes (cluster 1 in **Figure 4H**) tended to increase in expression with age across cell types. These included genes related to stress and glucocorticoid signaling (e.g., *FKBP5;* **Figure S14**), inflammatory pathways (*RSAD2, TNFSF13B, CD247*), DNA damage responses (*RAD51B, RPA1, BRIP1*), and metabolism (*SLC25A24, ALDH3A2, LSS*) — resembling age-associated gene-expression changes previously recognized in analyses of “bulk” brain tissues^57–59^.

Beyond this modest set of shared, pan-cell-type changes in gene expression, each cell type exhibited quite distinct age-associated changes (**Figure 4F-G, S13A**). For glial cell types, these changes were strongly concordant across striatal areas and the DFC (**Figure 4G, S13A**).

Among neurons, changes were strongly correlated only among certain closely related subtypes (such as D1 and D2 SPNs); SPNs, interneurons, and cortical pyramidal neurons showed only weak positive correlations with one another in their patterns of age-associated gene-expression change (**Figure 4G, S13A**). Age-associated changes in different glial cell types showed almost no correlation with one another or with the changes in neurons (**Figure 4G,H**).

In principle, age might affect the biology of some brain cell types more than others. This cannot be rigorously estimated from the number of differentially expressed genes (DEGs), since such estimates are strongly affected by the relative numbers of nuclei and UMIs of each cell type. To address this, we utilized two approaches. First, we used TRanscriptome-wide Analysis of Differential Expression (TRADE^60^); which estimates the underlying distribution of differential-expression effects. We also (to be certain) performed a version of the TRADE analysis with downsampling so that all cell types were represented by the same numbers of nuclei (see **Methods**) (**Figure 4I**). These analyses were strongly concordant with each other.

Projection neurons – including SPNs and cortical glutamatergic (pyramidal) neurons – exhibited a much larger impact of age on gene expression than interneurons did (**Figure 4H,I**). Among glial cells, microglia and astrocytes exhibited the largest age-associated changes, with much smaller changes in oligodendrocytes and OPCs (**Figure 4I**). Curiously, despite pronounced loss with age (**Figure 3A**), OPCs exhibited relatively modest age-associated effects on RNA expression (**Figure 4I**).

Several molecular programs declined with age in the neuronal classes sampled. Cellular respiration was enriched among the age-declining genes in all 14 neuronal populations, reaching FDR < 0.01 in 10; astrocytes showed a similar decrease, whereas the same terms showed strong enrichment among the age-increasing genes in microglia (cellular respiration: astrocytes, NES = −2.01, FDR = 9.8e-6; microglia, NES = 2.28, FDR = 4.5e-9). Translation- related gene-expression programs also declined with age across most types of projection neurons: cytoplasmic translation was negatively enriched in all four SPN subtypes and all four cortical populations of intratelencephalic-projecting (IT) neurons analyzed (FDR < 0.05). Among SPNs, age-downregulated programs also included cAMP metabolism in all four subtypes (FDR = 5.7e-4 – 0.0061), with *ADCY1*, *ADCY3*, *ADCY5*, *PDE4B*, *PDE7A*, and *PDE10A* among the genes with age-declining expression.

Glial populations showed distinct patterns. In microglia, increased expression of respiratory and mitochondrial programs was accompanied by reduced expression of TGF-β-response genes (FDR = 2.2e-4), including *TGFBR2, TGFB1, SMAD3, SKI,* and *SKIL*. Oligodendrocytes showed reduced expression of genes with roles in cell-projection morphogenesis (FDR = 4.0e-9) and increased expression of genes with roles in antigen processing and presentation (FDR = 9.4e-4), driven by *HLA-A*, *HLA-B*, *HLA-C*, *B2M*, *TAP1/2*, and *TAPBP*. OPCs showed reduced expression of cell-junction/adhesion and differentiation programs, including cell-junction assembly (FDR = 3.2e-8) and stem-cell differentiation (FDR = 2.3e-6); including *PDGFRA*, *HES1/5*, *SOX10*, and *NOTCH1*. Representative SPN genes from these and related enriched programs are shown in **Figure 5A**. Complete GSEA results for all neuronal and glial populations are provided in **Table S7**.

**Figure 5.**
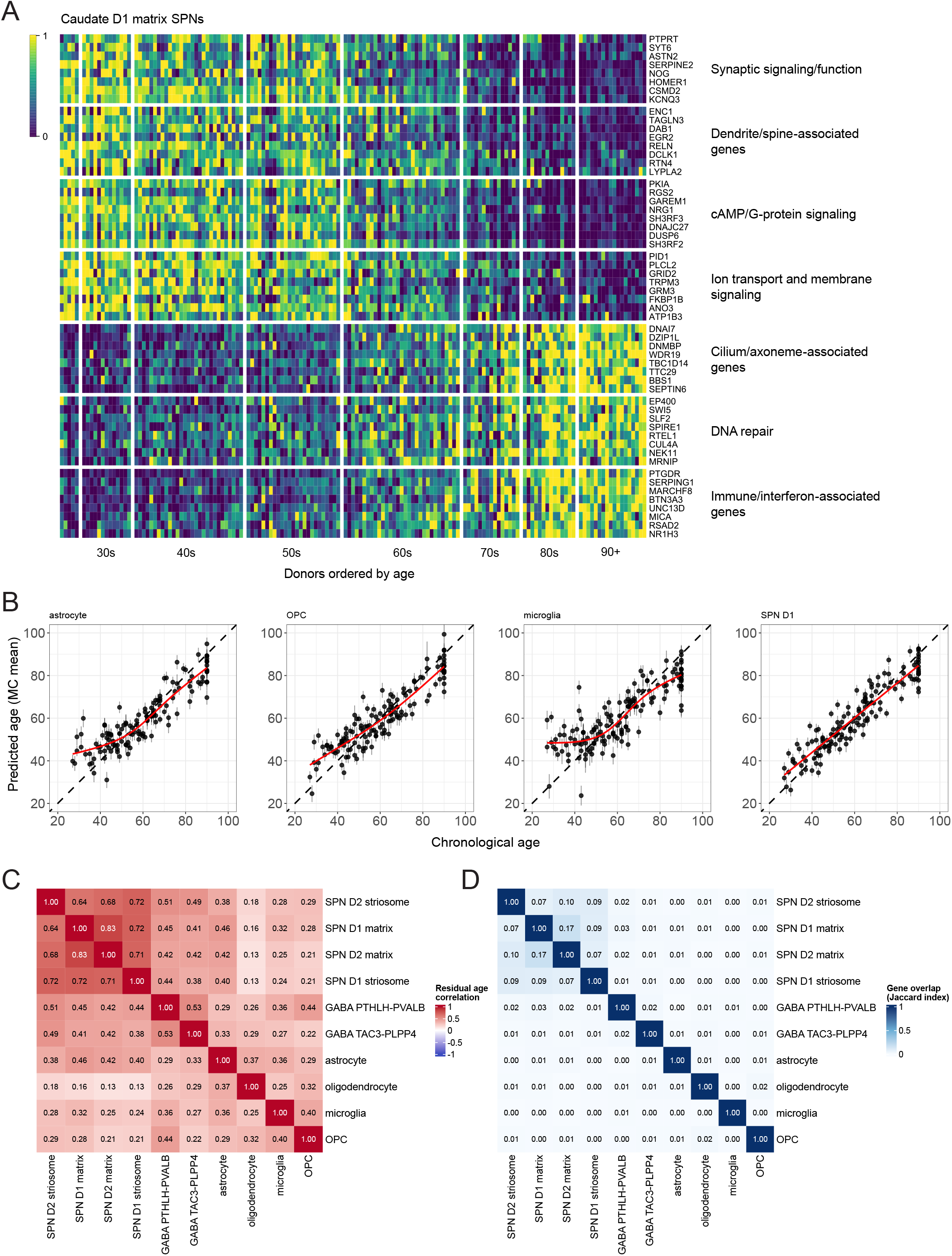
Chronological age can be predicted from gene expression in various cell types. (Extended analyses are in **Figures S15** and **S16**.) **A.** Age-associated changes in D1 matrix SPN gene expression, within the caudate. The 150 brain donors (columns) are ordered along the x-axis by ascending age (27 to 90+). Expression values (transcripts-per-million) for each gene (row) have been rescaled such that 0 (blue) represents the 10th percentile among all donors and 1 (yellow) represents the 90th percentile. **B.** RNA-expression-based age prediction in four representative caudate cell types. For each cell type, donor chronological age (x-axis) is plotted against RNA-expression-predicted age (y-axis), with predictions derived from an elastic net model. Each point represents one donor. The dashed line denotes the identity line (predicted = chronological age). Vertical error bars indicate plus or minus 1 SD around the predicted age. The red curve shows a generalized additive model (GAM) fit of predicted age as a function of chronological age. Age residuals (expression-predicted minus chronological age) are defined relative to this fitted relationship. **C.** Positive correlations of RNA-expression age- prediction residuals across cell types. The heatmap shows pairwise correlations, for each pair of caudate cell types, of GAM-corrected residuals between RNA-expression-predicted and chronological age. **D**. Minimal overlap of the genes used to predict age in most cell types. The heatmap shows Jaccard indices quantifying overlap among genes used in each cell-type- specific model.

### Inter-individual variation in RNA-expression-predicted age

An interesting question is whether age-associated RNA-expression patterns vary systematically among individuals, either broadly or in specific cell types. Inspired by the informativeness of methylation-based aging clocks in analyzing blood samples^61^, we asked whether cell-type-specific RNA-expression measurements could be informative about age-associated processes in the various brain cell types.

We constructed an age prediction model for each cell type using a penalized regression model (elastic net^62^, see **Methods**) (same extended donor set used for differential expression analyses). This model generated, for each donor and cell type, both a predicted age and an age residual (the difference between predicted and chronological age). We trained separate linear models for each cell type (by region) and first evaluated the ability of these models to estimate each donor’s age from RNA-expression data in the cell type alone (**Figure 5B, S15**). Across cell types, RNA-expression-predicted age was highly correlated with chronological age (**Figure 5B**). The median of the model-level mean absolute error was 6.6 years (range 4.7–9.6 years) (**Figure S15A**). We also trained a model constrained to select from those genes whose expression levels we had found to associate with age, yielding similar results (model-level mean absolute error: median 6.3 years; range 5.0–10.3 years; **Figure S16A-E**).

Within the striatum, model accuracy was greatest for SPNs (**Figure S15A**). Predictions at the youngest and oldest ages in the cohort were biased toward the cohort mean (**Figure 5B**); we corrected for this before analyzing variation in residual (predicted minus chronological) age across cell types and regions (**Figures S15D,E;** see **Methods**).

We next asked whether the differences (residuals) between RNA-expression-predicted and chronological age showed coordinated variation across cell types and brain regions. These residuals were positively correlated between all pairs of cell types in the analyses (**Figure 5C**), despite the fact that the genes used to predict age in different cell types were almost completely distinct (Jaccard index < 0.1 for most cell-type pairs; **Figure 5D**) and that, in general, these cell types had very different sets of age-associated expression changes (**Figure 4H**). These findings suggest a donor-level component of transcriptional variation shared across cell types and brain regions. These correlations were generally stronger when comparing the same cell type across brain regions than when comparing different cell types within the same brain region (**Figure S15H**). One possible explanation is that these cell-type-specific “aging clocks” capture, within an individual, coordinated biological processes associated with aging, along with some contribution of cell-type-specific variation.

### Effects of common genetic variation on gene expression

Humans exhibit abundant genetic variation. To map common genetic effects on gene expression in each of the cell types of the striatum and cortex, we jointly analyzed the snRNA-seq and whole-genome sequence data from the extended donor set (**Figure S10B**; cell type by region counts **Table S8;** see **Methods**). We identified 9,825 genes with significant evidence (at FDR < 0.01) of cis-regulation by a nearby SNP or haplotype (an “expression quantitative trait locus” or eQTL affecting an “eGene”) in one or more cell types (see three examples in **Figure 6A**). More deeply-sampled cell populations yielded more expression discoveries than sparser cell types did (**Table S9**). Though most of these eQTLs had not been reported previously – an earlier study of surgical cortical specimens from 75 adult donors^50^ had found about 200 eGenes per cell type – our eQTL results showed strong statistical support (in the form of sign tests) in the data from multiple earlier snRNA-seq analyses of the DFC (**Table S6**).

**Figure 6:**
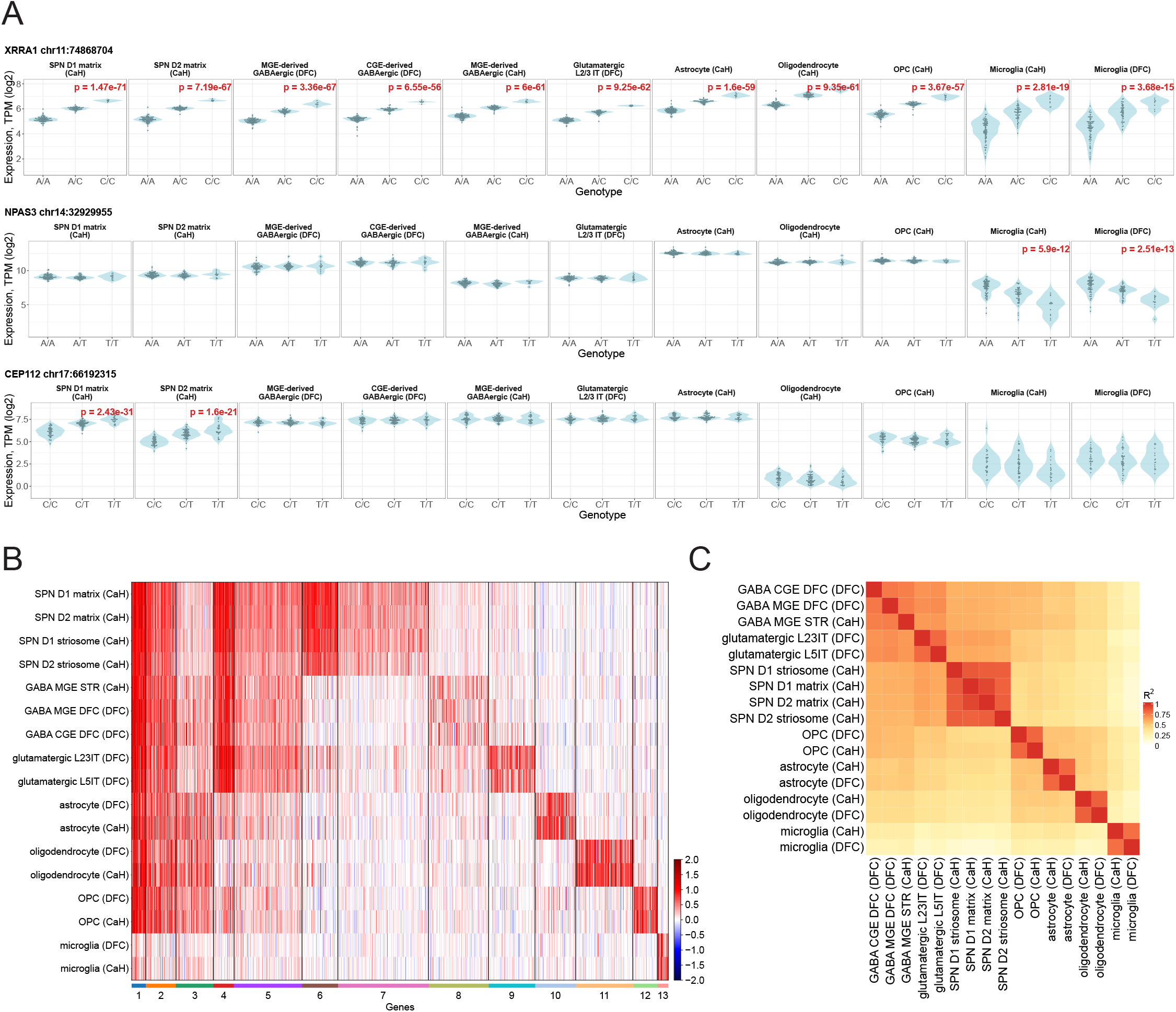
Effects of common genetic variation on gene expression. (Extended analyses including additional cell populations and brain regions are in **Figure S17**.) **A.** Log₂-transformed expression levels of three genes (of 9,825) whose expression levels associated with common genetic variants: *XRRA1* (rs10899052), *NPAS3* (rs61980314), and *CEP112* (rs7209037). Brain donors’ expression levels are shown for each genotype (as violin plots), for a variety of striatal and cortical cell types (red text indicates Benjamini–Hochberg adjusted p values). **B.** K-means clustering (K = 13) of the 9,825 eQTLs based on their patterns of effect sizes on gene expression across the cell types in the analysis. Effect sizes represent standardized change in gene expression (inverse-normalized log2-transformed values). For each effect, shades of red indicate the most common effect direction and shades of blue indicate any cell types with an opposite direction of association. **C.** Pairwise correlations (among cell types) of genome-wide sets of eQTL effects (corresponding to the rows in panel B). For each cell-type pair, signed Spearman’s ρ^2^ was calculated using eQTLs significant in at least one of the two cell types. Rows and columns are hierarchically clustered. The analyses in panels B and C are extended to additional brain regions in **Figure S17**.

To characterize patterns of cell-type specificity among these 9,825 eGenes, we used k-means clustering to group eQTLs by shared patterns of “effect sizes” – change in gene expression per allele inherited – across the full set of cortical and striatal cell types. Importantly, this analysis was indifferent to whether the eQTL relationship reached genome-wide significance in each of the cell types in the analysis (**Figure 6B**, see **Methods**), to minimize the effect of statistical power.

Some 15% of eGenes (n = 1,500; gene clusters 1–3 in **Figure 6B**) were shared across most cell types in the analysis (though often excluding microglia, which arise from a different developmental lineage^63^). Such effects varied in magnitude across the cell types but almost never varied in direction (**Figure 6A**, first row). Another 16% of the eQTLs (n = 1,619; clusters 4 and 5 in **Figure 6B**) were shared across many neuronal classes (including projection neurons and interneurons), often with more-modest effects in one or more non-neuronal cell types.

Some 68% of eQTL effects exhibited cell-type-specific patterns (clusters 6-13 in **Figure 6B**; see also **Figure S17** and **Table S10**). Examples included effects on *NPAS3* expression in microglia and *CEP112* expression in all of the SPN subtypes (**Figure 6A**). To validate these patterns, which had emerged from cell-type-specific analyses, we also used data for all of the cell types together in a single analysis in which we formally tested for genotype × cell-type interactions, focusing on eGenes with apparent cell-type-specific patterns (**Table S11;** see **Methods**;). Some 10.6% of eGenes were expressed exclusively in the implicated cell type; among the remaining testable eGenes, 80.9% showed the expected genotype × cell-type interaction (at p-value < 0.05) (**Table S11**). Downsampling to equalize statistical power across cell types did not materially change these estimates (see **Methods**, **Table S11**).

Like effects of aging, effects of common genetic variation in glial cells were shared/replicated across brain areas. Nearly all genetic effects that manifested in striatal astrocytes were also apparent in cortical astrocytes, and vice versa; the same was true for OPCs, oligodendrocytes, and microglia (**Figure 6B,C**).

SPN subtypes (D1 and D2 SPNs; and matrix and striosome SPNs) also tended to exhibit highly similar effects of common genetic variants (**Figure 6B,C**), as did SPNs from different striatal regions (**Figure S17**). Neurons with more-distinct molecular identities, however – such as SPNs, interneurons, and cortical pyramidal neurons – tended to manifest highly divergent sets of genetic effects (**Figure 6B,C**).

Two additional features of these regulatory effects concerned the genes they affected.

Among cell-type-specific eQTLs (in clusters 6–12), only a minority (24–41%) involved regulatory effects in the cell type that most strongly expressed the gene (this fraction was 62% for the microglia-specific cluster 13; **Table S10**). While predicted by emerging biological insights about the cell-type specificity of well-characterized enhancer activities, this result cautions against interpreting the cellular action of disease-implicated genetic variants based primarily on the cell type in which the nearby gene is most strongly expressed, a prevalent practice in “cell-type-enrichment” analyses of genome-wide association study (GWAS) results^64^.

Genes that are intolerant to loss-of-function mutations ^65^ tended to have smaller genetic effect sizes (change in gene expression per allele inherited) than less-constrained eGenes did (**Figure S18**; see **Methods**), consistent with recent findings for human blood eQTLs^69,70^ and suggesting that, as sample sizes expand, larger fractions of eQTLs involve evolutionarily constrained genes.

## Discussion

Decades of work by brain banks, and the generosity of brain donors and their families, is increasingly meeting new genomic analysis technologies that unlock the cellular and molecular information in these tissues. Human brain tissues offer unparalleled opportunities to investigate normal and pathogenic processes and the ways such processes vary from person to person. We hope these results will inform the design and interpretation of human brain studies, from selecting donors to appropriately calibrating statistical analyses.

One of the most remarkable observations from these data involved the degree to which cell type proportions vary from person to person (**Figure 2**). A fascinating question for much future work involves the biological and perhaps genetic causes of this variation. For OPCs, age-associated loss was profound but still explained only part of the inter-individual variation that manifested (in a correlated manner) across all of an individual’s brain regions (**Figure 3B**). SST+ interneurons in the cortex also showed attrition with age. Genetic factors seem likely to contribute, and as cohort sizes increase it will become possible to map such effects.

Age appeared to affect gene expression in some brain cell types much more than others (**Figures 4H,I** and **S13D,E**). SPNs showed the largest effects of aging on gene expression, followed by cortical glutamatergic (pyramidal) neurons; the various types of interneurons in these brain regions manifested much smaller changes. It is interesting to speculate whether this sensitivity arises in projection neurons’ need to support spatially distant metabolic and physiological processes, at scales that have greatly increased in human evolution.

One way in which these results challenged our expectations was that the cell types with the greatest vulnerability to age-associated loss – OPCs and certain types of interneurons – were not the cell types with the largest age-associated gene-expression changes (**Figure 4H**). In the case of OPCs (and, downstream, oligodendrocytes), it may be that renewal from a stem population attenuates the effects of aging, even as that stem population itself declines in capacity with age. In the case of interneurons, age-associated changes might lead to loss of the more severely affected interneurons – reducing their numbers, but also reducing the average amount of age-associated change among the remaining interneurons.

Comparing cell-type-specific effects across striatum and cortex, we found that sets of eQTLs for different types of neurons were quite distinct (**Figure 6**). This points to future discovery of a vast set of neuron-specific genetic effects — among which SPNs and cortical projection neurons represent only a small fraction of total neuronal diversity, and thus only early examples of, far more eQTL discoveries to come. Such expansions are expected to include a greater proportion of evolutionarily constrained genes.

Our hope is that these data and analyses, and much other such work in the future, will make natural human biological variation an ever-more-useful tool to neurobiology – a source not of unwelcome analytical complexity but of new questions and a deeper understanding of human biology and its dynamic range.

### Limitations of the study

Our analyses do not identify the biological mechanisms by which these sources of biological variation arise, nor (beyond the effects of common genetic variants on gene expression) make any particular claim about the relative contributions of genetics and environment. Such questions deserve extensive future work by a wide variety of scientific approaches. We hope our results inspire the conception of such studies and inform their design.

Humans have an enormous variety of subclinical biological conditions, as well as conditions that do not receive medical attention or a formal diagnosis. Our observations that certain kinds of biological variation appear frequently and in specific patterns make no claims about whether or not such biological variation is “healthy”, affects brain functions, or contributes to vulnerabilities – in fact, we believe this is an important topic for future research.

## Resource Availability

### Lead Contact

Requests for further information and resources should be directed to the lead contact, Steven A. McCarroll

### Materials Availability

This study did not generate new unique reagents.

### Data and Code Availability

In this study, we used data we generated for the BICAN (RRID:SCR_022794) which are accessed/deposited in the NeMO Archive (RRID:SCR_016152), under accession number: nemo:dat-pu20xdm.

Data may be accessed at: https://assets.nemoarchive.org/collection/nemo:dat-pu20xdm

All original code has been deposited at GitHub (RRID:SCR_002630) and is publicly available as of the date of publication: https://github.com/broadinstitute/bican-mccarroll-manuscript1 (RRID:SCR_028041).

## Acknowledgements

This publication was supported by and coordinated through the BRAIN Initiative Cell Atlas Network (BICAN) and funded by the National Institute of Mental Health (UM1MH130966). We thank the NIH NeuroBioBank network for providing tissue for this project. We also thank all of the brain donors and their families; this work would not be possible without their generous gifts to science. We thank Morgan Sheng’s lab for equipment access, and Bryan Song for image scanning advice. We also thank Sabina Berretta, Mark Daly, Elise Robinson, Marta Florio, and Avin Veerakumar for comments and suggestions on manuscript drafts.

## Author Contributions

S.A.M., K.I., and E.Z.M. conceived the study. C.V., C.J.M., H.G., N.A.R., K.K., J.M., A.M., S.D., N.B., and L.M. generated the data, with assistance from O.C., M.C., R.G., L.S., G.F., and C.C. S.C.B., J.N., O.Y., E.M., and J.Y. conceived and pursued the analyses, with assistance from K.S., M.G., S.B., A.W., L.R., N.A.R., H.F., G.G., S.J.F., N.R., and S.K., and mentorship from S.A.M. J.N., K.S., and A.W. created computational infrastructure and workflows that supported a variety of analyses. S.C.B., O.Y., J.N., E.M., and J.Y. interpreted data, with assistance from N.A.R., H.G., S.B., A.M., H.F., A.W.K., and N.K., and mentorship from S.A.M., K.I., and E.Z.M. S.K. created a browser for interacting with the data. E.F., M.H., and K.F. provided project management support. K.I., S.C.B., O.Y., E.M., J.N., J.Y., and S.A.M. wrote the manuscript, with input from all authors.

## Declaration of Generative AI and AI-assisted Technologies

During the preparation of this work, the authors used GitHub Copilot, GitHub Copilot CLI, Claude, ChatGPT, Le Chat, Perplexity, Phind.com, Groq, and Gemini for searching for the exact syntax of existing utility methods such as filtering and sorting, searching for packaged Docker images, software installation instructions, debugging software errors, generating workflow code from templates, and potential software performance optimizations. Grammarly was used for spell checking and grammar suggestions. Additionally, during the preparation of this work the authors used ChatGPT for minor editing/proofreading of manuscript text. After using these tools, the authors reviewed and edited the content as needed and take full responsibility for the content of the publication.

## STAR Methods

### Key resources table

Uploaded as a separate Word document

### Experimental model and study participant details

Postmortem human brain tissue was obtained through the Human Brain Cell Atlas Collection (HBCAC), a dedicated program within the NIH NeuroBioBank (NBB; RRID:SCR_003131) supporting the NIH BRAIN Initiative Cell Atlas Network (BICAN). Brain and Tissue Repositories contributing to the NBB HBCAC followed harmonized protocols for donor evaluation, brain collection, and tissue processing and preparation such as photodocumentation, slab thickness, and uniform sectioning. Postmortem human brain tissue was also obtained from the main NBB collection (“archival donors”). Archival donors appropriate for our study were selected by NBB, in accordance with HBCAC criteria. Postmortem tissue collection followed the provisions of the United States Uniform Anatomical Gift Act of 2006 described in the California Health and Safety Code section 7150 and other applicable state and federal laws and regulations. Informed consent for unrestricted research use and open data sharing was obtained by NBB from the legal next-of-kin. The Broad Institute Office of Research Subject Protection reviewed the use of postmortem brain tissue for research purposes and determined that the use of de-identified specimens from deceased individuals did not constitute human subjects research requiring IRB review (NHSR-8066) per federal regulation 45 CFR 46 and associated guidance.

Donor information is available in **Table S2**. Adult donors ages 27 to 90+ that were consented for open data sharing were considered for this study; donors without consent for open data sharing were not included. Exact donor age at time of death for donors over 90 years old is not available as a subject’s age, in combination with other health information, could potentially be a unique identifier for individuals aged 90 and older. These subjects are listed as aged 90+ years, following HIPAA Privacy Rules. Donors with positive serology results for HIV-1/2, Hepatitis B, or Hepatitis C were not considered for this study.

We sampled from donors flagged by NBB as potentially meeting the HBCAC criteria at the time of donor intake at the brain bank. All donors were evaluated for major or persistent neurological or psychiatric conditions, such as Alzheimer’s disease and schizophrenia. Toxicology testing was conducted at each contributing brain bank to determine substance presence at the time of death and to interpret substance exposure alongside clinical brain diagnoses. PMI information is available in **Table S2**. Donors were not excluded on the basis of PMI as this metric did not correlate with library size in our dataset (**Figure S19A,B**). All available donor metadata at the time of brain bank intake were comprehensively reviewed and used to guide donor selection and initial tissue sampling.

Donors that were later found to have major or persistent neurodegenerative or psychiatric conditions, or substance dependence (including alcohol, opioid, or cocaine) were not included in the core donor set (**Table S1**, **S2**). Additional exclusionary criteria for core donor set designation included clinical brain diagnoses of: recurrent major depressive disorder, and other psychiatric disorders (including post-traumatic stress disorder (PTSD), obsessive-compulsive disorder (OCD), and hoarding disorder). Exclusion criteria based on neuropathological evaluation included: Alzheimer’s disease, Cerebral infarction in donors under 65 years of age^67^, Primary age-related tauopathy (PART) in donors under 65 years^68,69^, and Limbic-predominant age-related TDP-43 encephalopathy (LATE) in donors under 80 years^70^. Age thresholds were informed by current literature characterizing the age-related onset of these pathologies.

Additional exclusionary metadata was based on causes of death, specifically: cause of death of combined overdose that includes substances of ≥2 drug classes and cause of death due to chronic substance use/abuse/dependence. Minor neurological or psychiatric conditions deemed to have minimal functional impact (e.g., migraine, specific phobias) were permitted. Donors with historical diagnoses in clinical remission were eligible, provided there was no confirmatory evidence of the disorder at the time of death per clinical records. Findings associated with normal aging or incidental focal pathology (e.g., localized ischemic events) were not considered exclusionary.

### Method Details

#### Tissue Procurement

To facilitate standardized tissue selection and annotation, brain slab images from HBCAC donors were uploaded to the custom-built Neuroanatomy-anchored Information Management Platform (NIMP; RRID:SCR_024684) developed by UTHealth Houston^71^. Brain slab image collections, prepared under HBCAC protocols, consisted of an average of about 30 coronally sectioned slabs, each approximately 5 mm thick. Slabs were photographed with identification numbers and scale bars, enabling tissue requesters to estimate dimensions accurately.

Neuroanatomists reviewed these slab collections and identified slabs that contained the relevant regions of interest (ROI). ROIs were drawn on the slab images to indicate the precise dissection sites and annotation pins were placed to link to neuroanatomical structures as defined in the Allen Institute HOMBA ontology. To maintain experimental uniformity across donors and brain regions, ROIs were matched as closely as possible in anatomical position, minimizing variability caused by normal morphological differences between individuals.

Archival donor tissue was requested through the NBB request process and subdissections were performed by NBB in accordance with NBB’s standard practice^72^. Tissue preparation varied by brain bank, with tissue arriving as small blocks, chips, or pulverized (tissue samples that were pulverized are designated as such in **Table S2**). DFC, caudate, putamen, and nucleus accumbens were sampled from the archival donors. Internal capsule tissue was not sampled for the archival donors.

#### RNA quality scoring

Genomic DNA and total RNA were extracted from frozen cerebellar tissue (or DFC if cerebellum was not available) from each individual donor and processed at Broad Clinical Laboratories.

RNA quality was assessed using the LabChip GX system (PerkinElmer). RNA integrity was evaluated by microfluidic capillary electrophoresis, generating RNA Quality Scores (RQS) on a scale of 1–10, with higher values indicating greater integrity. DV200 values (proportion of RNA fragments >200 nucleotides; reported on a 0–1 scale) were calculated as an additional measure of RNA quality predictive of RNA-sequencing performance. RQS and DV200 served as quantitative quality control measures as these values correlate with library size in our dataset (**Figure S19C**).

#### Tissue handling and dissections

The anatomical location of each ROI was confirmed by aligning slab images to the Ding atlas^66^ coordinates, with “pinned” reference points marking the anterior–posterior, dorsal–ventral, and medial–lateral positions. Care was taken to match ROIs across donors, ensuring uniform anatomical coverage despite individual morphological variation. DFC sampling targeted the cortical layers anterior to the premotor cortex and dorsal to the inferior frontal sulcus (**Figure S1D**). Striatal dissections were guided by internal landmarks to isolate the entirety of caudate nucleus, putamen, nucleus accumbens core and shell (treated as a single region for downstream analysis for consistency with archival donor sampling), and segments of the internal capsule without contamination from neighboring territories (**Figure S1E**). Dissection was conducted in a cryostat (−25 °C) using pre-chilled razor blades. Slabs were kept on dry ice until processing and immediately returned to −80 °C storage post-sampling. All cuts were made with care to avoid inclusion of unintended subregions and any adjacent white matter, particularly for small striatal nuclei. Subdissections were conducted by NBB staff for all donors designated as “archival”.

#### Nissl staining

Human striatum sections of 12 µm thickness from 14 donors were collected and stored at −80 °C. Sections were equilibrated to room temperature prior to staining, fixed in 70% ethanol for 2 minutes, followed by rehydration in distilled water for 30 seconds. Excess water was wiped off with kimwipe, and slides were stained with freshly filtered solution of 0.1% Cresyl Violet Acetate (Electron Microscopy Sciences 26089-01; spin filter: Corning Costar 0.22 µm Spin-X centrifuge filter) for 4 minutes. Excess dye was tapped off and the slides were rehydrated in water for 30 seconds, then sequentially fixed in 70%, 95%, and 100% ethanol for 30 sec, 30 sec, and 1 min, respectively. Lastly, slides were submerged in xylene for 5 minutes, mounted with Permount mounting medium (EMS, 17986-01). Nissl images were acquired using the Keyence BZ-X810 Microscope under brightfield channel, 4x magnification, and stitched with the Keyence Analyzer.

#### RNAscope

To quantify cell type proportions from intact human striatum sections, single molecule fluorescent in situ hybridization (smFISH) with RNAscope technology (ACDBio) was performed using the RNAscope Multiplex Fluorescent Kit v2 (ACDBio, 323100), following the manufacturer’s protocol. Specifically, 12 µm fresh frozen human striatum sections were collected and stored at −80 °C in air-tight containers. Tissue sections were fixed in 4% PFA for 1 hour at 4 °C on the day of RNAscope assay. Hs-SLC1A2-C2 and Hs-PDGFRA-C2 probes were used to detect astrocytes and oligodendrocyte precursor cells (OPCs) with the ClariTSA Fluorophore 570 (ACDBio, 323272) at 1:750 dilution. TrueBlack (biotium, 23007) was used for lipofuscin autofluorescence signal quenching at the end of the RNAscope assay, and slides were mounted with the ProLong Diamond antifade mountant (Invitrogen, P36961).

Fluorescence tile scan images were acquired using the ZEISS Axioscan 7 Slide Scanner at 20x magnification for QuPath analysis. Additional representative images were acquired using the ZEISS LSM 900 confocal microscope at 20x magnification.

#### Whole genome sequencing

Whole-genome sequencing (WGS) was performed on genomic DNA from each donor. PCR-free libraries were prepared using the KAPA HyperPrep Library Construction Kit (Roche) with custom Broad indices. Libraries were sequenced on the NovaSeq X platform (Illumina; RRID:SCR_024569) to generate 150 bp paired-end reads to a target mean coverage of 30x.

Reads were demultiplexed, aggregated, and aligned to the hg38 reference genome using Illumina DRAGEN processing. Data delivery included aggregated CRAM files, corresponding CRAI index files, and md5 checksum files.

#### Bulk RNA-seq library preparation

Up to 20 mg of frozen tissue per donor was disrupted and homogenized in 350 µL Buffer RLT Plus (Qiagen # 74134 RNeasy Plus Mini Kit) with an Omni THQ Tissue Homogenizer, and purified RNA was extracted following the RNeasy Plus Kit protocol. RIN and yield analysis was performed using an RNA ScreenTape (Agilent 5067-5576), and up to 1 µg (when yield allowed) of intact total RNA per sample was used as input into the NEBNext rRNA Depletion Kit v2 (Human, Mouse, Rat) protocol (NEB E7405). Ribo-depleted RNA was then constructed into sequencing libraries following the NEBNext Ultra II Directional RNA Library Prep Kit for Illumina (NEB E7760) protocol, with dual indices attached using the NEBNext Multiplex Oligos for Illumina (NEB E6440S) kit. All RNA samples were high quality and received an RNA fragmentation time of 8 minutes in the NEBNext protocol. All samples received 7 Index PCR cycles in the NEBNext protocol. Completed libraries were quantified then pooled together into groups of 16 and sequenced on one lane of an Illumina NovaSeqX 25B 300c flow cell per pool with the sequencing parameters R1: 151 bp, R2: 151 bp, i7: 8 bp, i5: 8 bp.

#### snRNA-seq library preparation and sequencing

To enable direct comparative analysis of nuclei from multiple brain donors while minimizing technical variance, we processed frozen brain tissue specimens from multiple donors as a single pooled sample. Each village comprised nuclei isolated from an average of 19 donors (range: 9-26 donors). For each donor, equivalent amounts of tissue (30 ± 5 mg for most regions, ∼15 mg for smaller anatomical regions) were collected. All pooled samples underwent nuclei isolation, droplet-based encapsulation, library preparation, and sequencing as a single batch. Between 2-8 encapsulation reactions were performed per village, depending on the desired sequencing depth. We used combinations of hundreds of transcribed single nucleotide polymorphisms (SNPs) in each cell’s sequenced reads to assign each nucleus to its donor-of-origin, using the computational approach described below.

Nuclei were isolated from frozen brain tissue using a modified density gradient protocol^73^. Briefly, tissue blocks were cryosectioned/microdissected at −25 °C and transferred directly to pre-chilled Dounce homogenizers containing Nuclei EZ lysis buffer (MilliporeSigma, #NUC101) supplemented with 1 U/µL RiboLock RNase Inhibitor (Thermo #EO0382). Mechanical homogenization was performed with 10-20 strokes of each pestle size, followed by incubation on ice for 10 min. The lysate was passed through a 20 µm vacuum cell strainer (SCNY00020), centrifuged at 500 × g for 5 min at 4 °C, and the pellet resuspended in G30 solution (30% iodixanol (STEMCELL #07820), 3.4% sucrose, 20 mM Tricine, 25 mM KCl, 5 mM MgCl₂, pH 7.8). The suspension was layered over a fresh G30 cushion in 1.5 mL volumes and centrifuged at 8000 × g for 10 min at 4 °C. Supernatants were discarded, and resulting nuclei pellets were resuspended in PBSAi buffer (composition as optimized in our protocol) and pooled into a single tube per village. Nuclei suspensions were brought to 1.5 mL total volume with PBSAi, centrifuged again (500 × g, 5 min, 4 °C), and the pellets resuspended in ∼100 µL of PBSAi. Nuclei were counted using a LUNA-FL Dual Fluorescence Cell Counter (Logos Biosystems, #L12005) with acridine orange/propidium iodide staining (Logos Biosystems #F23001) to assess concentration and purity.

Isolated nuclei were loaded onto a 10x Genomics Chromium instrument (RRID:SCR_024939) and processed using either Chromium NextGEM Single Cell 3’ Reagent Kits V3.1 (CG00204, Rev D) or Chromium GEM-X Single Cell 3′ Reagent Kits v4 (CG000731, Rev A) according to the manufacturer’s protocol, with minor modifications for nuclei input (19-22K loading for NextGEM, 29-32K loading for GEM-X). The majority of libraries were generated from small-scale (8-16 reactions) manual preparations using magnetic racks, while a subset of experiments were processed using automated high-throughput liquid-handling systems. Sequencing was performed on the Illumina NovaSeqX platform.

### Quantification and statistical analysis

#### Sequence alignment and donor assignment

Sequence data was demultiplexed and aligned following the standard Drop-Seq protocol^74^ and was aligned to the GRCh38 reference and GENCODE (RRID:SCR_014966) v43 gene models. The expression matrix was computed using Drop-Seq DigitalExpression program, and used two non-standard flags READ_MQ=0 and FUNCTIONAL_STRATEGY=STARSOLO to replicate STARSolo expression quantification. Ambient/ background RNA was removed from digital gene expression (DGE) matrices using CellBender (v.0.3.2) remove-background^75^ (RRID:SCR_025990). Nuclei selection was performed via DropSift (RRID:SCR_028040; see below), then final DGE matrices were generated for those subsets of cell barcodes. These matrices were then used in combination with MapMyCells (RRID:SCR_024672)^76^ to generate cell type labels. Donor assignment and doublet detection were performed by Dropulation^34^.

#### Nucleus selection

Accurate nucleus selection is a critical step in snRNA-seq analysis. Proper identification of nuclei ensures robust downstream expression quantification, cell type classification, and biological interpretation. However, distinguishing nuclei from empty droplets presents unique challenges, particularly in brain tissue, where high cellular diversity, ambient RNA contamination, and debris can complicate classification. To address these challenges, we developed DropSift, a supervised method that distinguishes nuclei from empty droplets in an experiment-specific manner.

Nucleus-containing droplets were distinguished from empty droplets using DropSift^77^. DropSift is a supervised classifier that labels barcodes as nuclei or empty droplets using a support vector machine (SVM) trained separately for each experiment. The model uses cell-level summary metrics (including total UMI counts and the fraction of intronic UMIs), expression-based features derived from genes differentially expressed between nuclei and empty droplets, and, when available, CellBender-derived estimates of ambient RNA contamination as input features. For each dataset, DropSift first identifies exemplar nuclei and empty droplets using density-based thresholds in the joint distribution of total UMI counts and intronic fraction, with additional guidance from the CellBender contamination estimates. These exemplars are used to train an SVM with a radial basis function kernel to discriminate nuclei from empty droplets. The trained classifier is then applied to all barcodes above a minimal UMI threshold; very low-UMI barcodes (<20 transcripts) which cannot be reliably classified are excluded from downstream analysis.

By integrating multiple signals, DropSift effectively differentiates nuclei from empty droplets while adapting to variability in data structure, including differences in ambient RNA contamination, across experiments and species. Rather than relying on tissue- or organism- specific marker sets, it learns expression patterns that are specific to each dataset, which improves classification accuracy in heterogeneous brain tissue.

DropSift is implemented in R and released as an open-source package. The source code, documentation, and installation instructions are available from the project GitHub repository (https://github.com/broadinstitute/DropSift)

#### Unsupervised clustering

Following nuclei selection to establish a set of non-empty, non-debris libraries, we performed unsupervised clustering to identify cell-type structure in the dataset. All clustering analyses were performed in R using Seurat^78^ (v4.4.3; RRID:SCR_007322), with analysis-specific parameters (e.g., number of highly variable genes, number of principal components, and clustering resolution) specified below.

Raw count matrices were preprocessed using Seurat’s standard single-cell workflow. Gene expression values were log-normalized using NormalizeData. Highly variable genes (HVGs) were selected using FindVariableFeatures, modeling the gene mean-variance relationship, and HVG expression values were linearly scaled using ScaleData such that each gene had zero mean and unit variance across cells. Mitochondrial (MT-) genes were excluded from clustering analyses; no additional manual gene filtering or regression was applied. Principal component analysis was performed using RunPCA on the selected HVGs. Cell-cell neighbor graphs were constructed using FindNeighbors, followed by graph-based clustering using FindClusters (Leiden algorithm). Low-dimensional embeddings were computed using UMAP via RunUMAP. To encourage co-clustering of nuclei across donors, and in some analyses across brain regions, Harmony-based integration was applied using Seurat’s Harmony wrapper (implementation specified per analysis).

Due to the large scale of the dataset (∼5 million nuclei when including all donors, samples, and doublets), analyses were performed using BPCells^79^, an R package that enables memory-efficient storage and streaming of large single-cell expression matrices. BPCells was used in conjunction with Seurat to support normalization, dimensionality reduction, clustering, and visualization at scale.

#### Expression-based doublet removal

Donor multiplexing (experimental) with genotypic demultiplexing (computational) enabled identification of genotypic doublets, defined as nuclei containing genetic material from more than one donor. Experimental batches (“villages”) typically included approximately 20 donors; under even donor representation, the expected proportion of heterotypic genotypic doublets is ∼0.95.

To identify additional doublets based on transcriptional profiles, genotypic doublets were intentionally retained during initial clustering to serve as a reference set. Global clustering was first performed using 5,000 HVGs and 60 principal components, followed by iterative sub-clustering at higher resolution. Initial partitions separated major cell classes, including striatal neurons, cortical glutamatergic neurons, striatal GABAergic interneurons, cortical GABAergic interneurons, astrocytes / OPCs / microglia, oligodendrocytes, vascular cells, ependymal cells, and lymphoid lineage immune cells. These groups were subsequently sub-clustered to generate terminal (“leaf”) clusters corresponding to finer cell-type structure.

For each leaf cluster, the proportion of nuclei labeled as genotypic doublets was computed. Clusters containing >85% genotypic doublets were removed in their entirety. All remaining nuclei identified as genotypic doublets were then removed from retained clusters. No additional expression-based doublet detection algorithms were applied.

This procedure yielded a curated dataset with doublet annotations and an initial set of unsupervised cell-type clusters used in downstream analyses.

#### Label transfer from external taxonomies

To support initial annotation and evaluation of unsupervised clustering results, cell-type labels were transferred from external reference taxonomies. For striatal populations, the consensus basal ganglia taxonomy^80^ was used as the primary reference. Label transfer was performed using the MapMyCells computational framework.

For each nucleus, best-match labels were computed at multiple hierarchical levels (neighborhood, class, subclass, group, and cluster), along with bootstrap-derived confidence probabilities. Rather than relying on nucleus-level assignments, label transfer results were summarized at the cluster level and used to assign initial putative labels to unsupervised clusters.

In some samples, particularly from nucleus accumbens dissections, sub-clusters were identified that corresponded to neuronal populations outside the target regions of analysis (e.g., globus pallidus GABAergic neurons). These populations formed distinct sub-clusters and were recorded as off-target dissection artifacts in sample-level metadata; downstream analyses focused on neuronal populations within the defined target regions (CaH, Pu, NAC, and ic).

Because the dataset also included cortical nuclei (DFC) as a comparator group, some non-basal-ganglia populations were not represented in the basal ganglia taxonomy. Although MapMyCells assigned labels to these populations, these assignments were not used in downstream analyses. For cortical neuronal populations, additional label transfer was performed using structured neuronal taxonomies from the Allen Brain Atlas (RRID:SCR_017001).

Annotated snRNA-seq data from human postmortem DFC^25^ were used to define exemplar populations. scPred^81^ models were trained on these reference datasets and applied to DFC-derived neuronal nuclei. Labels were assigned based on maximum predicted probability, with a minimum probability threshold of 0.8.

#### Sub-clustering neuronal and glial populations

After removal of genotypic and expression-defined doublets (see **Methods**: Outlier identification and filtering), cell-type-specific sub-clustering was performed to achieve higher-resolution annotations.

The initial global clustering was sufficient to resolve certain populations (e.g., ependymal cells, B cells, and T cells), which were not further subdivided. Additional sub-clustering was performed for neuronal and major glial populations as described below. Input nuclei for sub-clustering analyses were filtered to libraries with a log_10_ UMI count equal to or greater than two median absolute deviations (MADs) below the median log_10_ UMI count for the cell type being clustered, calculated for each donor “village” to account for experimental batch effects influencing UMI counts.

Glial populations analyzed included astrocytes, oligodendrocyte precursor cells (OPCs), committed oligodendrocyte precursors (COPs), oligodendrocytes, microglia, and vascular cells. Vascular cells were sparsely sampled across donors and were excluded from downstream cell- type proportion and gene expression analyses.

Microglia were sub-clustered using 3,000 HVGs, 30 principal components, and Harmony integration by donor and brain region. A small sub-cluster of non-microglial myeloid cells annotated as border-associated macrophages (BAMs) by label transfer from the basal ganglia taxonomy was excluded from microglia analyses.

Oligodendrocytes were sub-clustered using 3,000 HVGs, 30 principal components, and Harmony integration by donor and brain region. Labels from the consensus basal ganglia taxonomy (Oligo OPALIN and Oligo PLEKHG1) were used to support annotation.

#### Striatal neuron sub-clustering

Striatal neuronal sub-clustering was performed to confirm representation of expected neuronal classes and to achieve high-resolution annotation. SPNs, eccentric SPNs, and striatal interneurons were jointly sub-clustered using 5,000 HVGs, 30 principal components, and Harmony integration by donor.

Clustering was performed at resolution 0.25. Major striatal interneuron classes defined in the consensus basal ganglia taxonomy (PTHLH–PVALB GABA, TAC3–PLPP4 GABA, SST– CHODL GABA, and cholinergic GABA neurons) formed distinct clusters, supported by marker gene expression and label transfer.

SPNs separated into D1 and D2 populations. Within these populations, cells were further annotated along matrix-striosome and dorsal-ventral axes. For downstream analyses, matrix and striosome SPNs were defined by the intersection of unsupervised cluster membership (restricted to D1 or D2 clusters) and confident label transfer assignment (probability >0.8).

We also identified additional, rarely observed neuronal populations — 0.2% median representation of CAP neurons, with a median absolute deviation of 0.3% (i.e., representation was highly variable by sample). Label transfer using MapMyCells and the HMBA basal ganglia reference assigned these neurons to groups associated with regions outside the intended sampling domain, including the hypothalamus (HTH), external segment of globus pallidus (GPe), and olfactory bulb (OB). We manually annotated these neurons as “out of scope”; however, the complete set of MapMyCells label assignments is provided for each population.

#### Astrocyte sub-clustering

Astrocytes were sub-clustered across brain regions using 3,000 HVGs, 30 principal components, and Harmony (RRID:SCR_022206) integration by donor and brain region. Clustering at resolution 0.25 yielded 11 astrocyte sub-clusters comprising ∼315k nuclei. Two major clusters corresponding to striatal and cortical astrocytes collapsed at lower clustering resolutions and were combined for exploratory analyses (**Figure S20**).

As higher-resolution astrocyte annotations were not available in the consensus basal ganglia taxonomy, marker gene identification was performed using comparison of cluster pseudobulked expression profiles. For each sub-cluster, the top 20-50 genes with the largest fold changes relative to cluster 0 were selected as candidate markers. These gene sets were compared with published markers of astrocyte subtypes and states in the adult human brain (these references were also valuable while exploring OPC clusters observed in our data, described below) ^52,53^ (**Figure S20B**).

A single cluster (cluster 0) accounted for the majority of astrocytes (63%) and expressed canonical homeostatic astrocyte markers, including *SLC1A2*, *KCNJ10*, *FGFR3*, and *GRM3*, consistent with a population of homeostatic “protoplasmic” astrocytes. Although this population was present across all sampled regions, there was a pronounced enrichment in gray matter regions relative to ic (**Figure S20C,D**). Cluster 1 (14%) was characterized by elevated expression of *GFAP*, *VCAN*, *TNC*, *CD44*, and extracellular matrix-associated genes, and was strongly enriched in the ic (**Figure S20C,D**), consistent with white-matter-associated “fibrous” astrocytes.

Among lower-abundance clusters, we observed evidence of regionally enriched astrocyte populations (**Figure S20B-D**). Cluster 6 (2%) showed near-exclusive enrichment in DFC samples and expressed higher levels of *DPP6*, *DPP10*, *SEMA5A*, and *DSCAML1*. This population may correspond with “interlaminar” astrocytes localized to the cortex. Cluster 3 (4%) expressed *CHRDL1*, *DIO2*, and *CALCRL* and was significantly enriched in putamen samples, potentially corresponding to a localized astrocyte population within the striatum, or astrocytes originating from an anatomically adjacent compartment resulting from off-target dissections.

Cluster 5 (3%) was marked by expression of *SLC6A11* together with GABA-signaling-related genes (*GABBR2*, *RGS6*, *ADCY1*) and was enriched in NAC and ic samples. This population may reflect a recently reported *SLC6A11*+ astrocyte population localized to the globus pallidus, captured in more-posterior samples targeting the NAC.

Cluster 9 (1%) was challenging to define. Elevated expression of *AQP4* would be consistent with perivascular astrocytes, but we did not identify a larger blood-brain-barrier associated program.

Two clusters seemed to reflect astrocyte states rather than stable subtypes.

Cluster 2 (2%) expressed inflammatory and cytokine-responsive genes including *C3, CHI3L1, F3, NRCAM, CCL2, SOCS3, OSMR, SERPINA3*, and *CXCL10*. This astrocyte cluster closely resembles the Astro_6 and Ast.5 clusters reported by Gabitto et al. (2024) and Green et al. (2024), respectively ^52,53,82,83^.

Cluster 7 (2%) was distinguished by robust induction of immediate early genes (*FOS*, *JUN*, *NR4A1/2/3*) and stress-response chaperones (*HSPA1A/B*). Perimortem conditions have the potential to impact expression levels of activity/stress programs, so we interpret the presence of this cluster with caution.

Two clusters likely reflect technical artifacts. Cluster 4 (4%) expressed neuronal and synapse-associated transcripts (*RBFOX1*, *SYT1*, *RIMS2*), raising the possibility of low-level neuronal RNA contamination or mixed nuclei, although a contribution from astrocytes with strong synaptic coupling cannot be excluded. Cluster 8 (2%) expressed oligodendrocyte-lineage markers (*MBP*, *PLP1*, *MOBP*, *BCAS1*) and likely represents oligodendrocyte or myelin-debris contamination.

#### OPC sub-clustering

OPCs were sub-clustered using 3,000 HVGs, 30 principal components, and Harmony integration by donor and brain region. Clustering at resolution 0.1 yielded eight sub-clusters, two of which collapsed at lower resolution and were combined, resulting in seven OPC sub-clusters (∼155k nuclei; **Figure S9A**).

To confirm that regional integration did not obscure region-specific structure, region-specific clustering was also performed using 3,000 HVGs for CaH, Pu, and NAC and 2,000 HVGs for ic and DFC, with 30 principal components and Harmony integration by donor. The same seven OPC subtypes were recovered in each region-specific analysis. Marker genes were identified using the same pseudobulk-based approach as for astrocytes^54^ (**Figure S9B**).

The resulting clusters exhibited highly skewed abundances (**Figure S9C,D**). One dominant cluster (cluster 0) accounted for approximately 84% of all OPCs, whereas the remaining clusters each comprised between 1–7% of the population. Despite these differences in abundance, all clusters were present to some degree across brain regions (in contrast with region-restricted sub-clusters like astrocyte cluster 6), with the possible exception of white matter-localized OPCs noted below.

Cluster 0, the most abundant OPC population, expressed canonical OPC-associated genes including *BCAN*, *PCDH15*, and *ITGA8*, and lacked markers of proliferation, stress responses, or inflammatory signaling. The proportion of OPCs in this cluster was lower in the internal capsule relative to striatal gray matter regions; however, this pattern could be driven by the regional distribution of cluster 1 OPCs.

Cluster 1 (7%) was enriched in the internal capsule. We note several genes upregulated in cluster 1 (*CDH19*, *SEMA3E*, *GRIA4*), and several depleted in cluster 1 relative to cluster 0 (*SHISA6*, *GPC5*, *CNTN5*, *TCF7L1*). In spatial data, the former gene set marked OPCs localized to the ic and the latter marked OPCs within striatal gray matter regions. In contrast, genes upregulated in cluster 0 marked OPCs in all striatal regions, consistent with an OPC population localizing throughout gray matter and white matter compartments. The ratio of cluster 1 to cluster 0 OPCs was also found to associate strongly with oligodendrocyte fraction in all regions.

Cluster 2 (3%) represented a smaller but distinct OPC population characterized by strong upregulation of *MIR137HG*, *DPYD*, and *LINC01776*, all located within the 1p21.3 genomic locus. This cluster was present across striatal and cortical brain regions, as well as across donors (**Figure S9C,D**).

Cluster 3 (2%) expressed inflammatory and cytokine-responsive genes including *ARHGEF3, CAMK2D, SORBS1, OSMR, IL6R, GFAP,* and *STAT3*. This OPC cluster closely resembles the OPC_2_2 and OPC.3 clusters reported by Gabitto et al. (2024) and Green et al. (2024), respectively ^52,53,82,83^.

Cluster 4 (1%) was defined by expression of cell-cycle genes (*MKI67*, *TOP2A*, *CENPF*) indicating an actively mitotic OPC population. This cluster likely represents a transient cell-cycle state superimposed on other OPC identities.

Cluster 6 (1%) was marked by expression of hypoxia- and metabolism-related genes including *VEGFA*, *EPAS1*, and *SLC2A1*. Given its low abundance and transcriptional profile, this cluster may reflect a metabolic or hypoxic stress-associated OPC state, potentially influenced by local vascular environments or perimortem factors.

Finally, cluster 5 (1%) expressed mature oligodendrocyte and myelin genes (*MOG*, *MOBP*, *OPALIN*) but was transcriptionally distinct from COPs, suggesting contamination by oligodendrocyte nuclei (OPC-oligodendrocyte doublets) or myelin debris. This cluster was excluded when interpreting OPC heterogeneity.

#### Visualizing Slide-tags marker gene expression

To visualize the spatial distribution of expression of OPC subcluster marker genes, we utilized Slide-tags spatial transcriptomics data generated and analyzed by Kraft et al., 2026. These data is publicly available in https://assets.nemoarchive.org/collection/nemo:dat-aqicdbo. We chose to visualize gene expression in an exemplary sample (s5) that captured all of the striatal regions of interest.

#### Computing genetic principal components

Genetic principal components (PCs) were computed by co-embedding our samples with high- coverage whole-genome sequencing data from the 1000 Genomes Project^84^ (RRID:SCR_006828). We first restricted the 1000 Genomes VCFs to variants detected in our dataset and filtered to retain only common variants (minor allele frequency > 0.05). Sex was inferred using PLINK^85,86^ (RRID:SCR_001757).

Prior to PCA, SNPs were pruned to ensure high-quality, independent markers. Specifically, we removed SNPs with extreme allele frequencies (minor allele frequency < 0.005), high missingness (>2%), excess heterozygosity (Hardy–Weinberg^87,88^ equilibrium p-value < 1×10⁻⁶), and autosomal SNPs showing association with sex. Long-range linkage disequilibrium (LD) regions were also excluded. PCA was performed using EIGENSOFT^89,90^ smartpca (RRID:SCR_004965) on the 1000 Genomes reference panel; our samples were subsequently projected into this PC space (**Figure S1B**).

Genetic ancestry proportions were estimated by fitting linear models for each of the five superpopulations as a function of the top 10 genetic principal components. Models were trained on individuals from the 1000 Genomes project with known ancestry, and predicted values with 95% confidence intervals were generated for all samples^91^.

#### Outlier identification and filtering

Throughout this paper, various levels of outlier filtering were used for analysis (see **Table S1**). Unsupervised clustering was completed using all nuclei (no filtering). Cell type proportions (CTP) analyses, including exploratory analyses of cell type abundances (fractions, ratios) and regressions with metadata, were performed on the core donor set, excluding CTP outlier samples and donors. Cell type-specific analyses, including differential expression and downstream TRADE analysis, as well as RNA-based age modeling and expression quantitative trait locus (eQTL) analyses, were performed with CTP outlier samples, CTP outlier donors, gene expression (GEX) outlier samples, GEX-cell type outlier samples, and GEX-donor outlier samples removed.

#### Gene expression outliers

To identify gene expression (GEX) outliers, we analyzed pseudobulked expression profiles defined at the level of region × village × donor × cell type. Detecting outliers at the cell-type level is critical for downstream cell-type-specific analyses, including differential expression and eQTL mapping. Exclusion of outlier samples can improve statistical power by increasing signal-to-noise ratios, and help minimize spurious signals arising from a small subset of atypical samples. We did not distinguish between biological and technical sources of variation, as samples that deviate strongly from the population distribution may introduce noise regardless of origin.

For each region-village-donor-cell type combination, we generated pseudobulked metacells by aggregating single-cell UMI counts. Within each brain region and cell type, samples with log- transformed library sizes less than the regional mean minus 1.96 standard deviations were removed to exclude poorly ascertained samples. After filtering for library size, remaining metacells had their gene counts normalized to counts per million (CPM) and log-transformed to stabilize variance.

Pairwise Pearson correlations were then computed between all remaining samples within each region-cell-type. For each sample, the median Pearson correlation with all other samples in the same region-cell-type was defined as a conformity score, as described in *Ling et al.*^25^.

Conformity scores were modified z-score-normalized within each region-cell type, and samples with conformity z-scores less than −3.75 were classified as GEX outliers.

Outlier detection was restricted to cell types with sufficient coverage for inclusion in differential expression and eQTL analyses. Glial cell types included oligodendrocytes, astrocytes, oligodendrocyte precursor cells (OPCs), and microglia. Neuronal cell types included SPNs in striatal gray matter regions; because conformity scores were highly correlated across SPN subtypes, SPN D1 matrix neurons were used as a proxy for all SPNs. In DFC, GABAergic VIP interneurons, GABAergic PVALB interneurons, and glutamatergic L2/3 intratelencephalic (L23IT) projection neurons were included.

Donors for whom more than 50% of samples were flagged as GEX outliers (based on either low library size or low conformity score) were designated as donor-level GEX outliers and excluded from selected analyses. Additionally, donors for whom more than 50% of samples for a given cell type were flagged as GEX outliers were designated as donor-cell-type GEX outliers, and only samples of that specific cell type were excluded. Donor-level filtering was applied prior to donor-cell-type-level filtering.

#### Cell type proportion outliers

Cell type proportion outliers were determined using a sequential filtering approach similar to the gene expression approach described above. Cell type proportion vectors were generated by aggregating and normalizing cell type counts at the subclass level (annotation_sub_class_complete) for each distinct combination of donor and village. Donor × village samples with log-transformed nuclei counts less than the region mean minus 1.96 standard deviations were removed to exclude poorly ascertained samples.

Following removal of samples with low nuclei counts, gray matter samples (from CaH, Pu, NAC, and DFC) with unexpected neuron types contributing to more than 20% of the neuronal population were identified and removed in order to exclude potential misdissections. Since the abundance of neurons is much lower in white matter, this step was not performed for internal capsule samples. Unexpected neuron types for all brain regions included the neuronal subclusters identified from outside the target regions described previously. In the striatum, any cortical intratelencephalic neurons and cortical interneurons were deemed unexpected; likewise, all SPN types were deemed out-of-distribution for DFC.

For the remaining samples, cell type proportions were transformed with the arcsine square root transformation to stabilize their variance, and conformity scores were then computed as the modified z-score of the median (within-region) pairwise Pearson correlation for each sample. Samples with a conformity score less than −3.75 were labeled as cell type proportion outliers.

Whole donors were removed from some analyses on the basis of cell type proportions either if all, or if at least 3, of their samples were flagged as outliers.

15% of overall samples were flagged as cell type proportion outliers. This varied by region – 6% of DFC samples were cell type proportion outliers, 10% of CaH samples, and about 20% of NAC, Pu, and ic. Due to the nature of their position in the striatum, NAC, Pu, and ic were most likely to be affected by dissection artifacts from adjacent regions.

#### Cell type proportion analyses

For each donor and brain region, we aggregated counts of all single cells with cell type annotations. These aggregated counts were then used to calculate both the proportions of individual cell types relative to the total number of cells and ratios between cell types of interest. Donors with exclusionary metadata criteria were removed from these analyses (see **Methods:** Experimental model and study participant details**; Table S1,S2**).

#### Comparisons across brain regions

To examine differences in cell type composition across brain regions, we first calculated the proportion or ratio of each cell type for each donor-region combination. For the ratio-based neuron analyses, we removed outliers with z-scores greater than 5, where z-scores were calculated separately within each brain region. We performed pairwise comparisons between regions using the paired Wilcoxon signed-rank test.

To assess the consistency of cell type abundances and ratios across regions within individual donors, we calculated Spearman’s rank correlation coefficient across all pairs of regions for each metric of interest. These comparisons encompassed five categories: (1) cross-region correlations of major cell class proportions (astrocyte, microglia, oligodendrocyte, OPC, projection neuron, and interneuron) across five regions; (2) cross-region correlations of glial:neuron ratios (astrocyte, microglia, oligodendrocyte, and OPC) across four regions; (3) cross-region correlations of SPN subtype ratios (D1:D2, striosome:matrix, eccentric:canonical) across three regions; (4) cross-region correlations of interneuron:SPN ratios across three regions; and (5) within-region correlations between interneuron:SPN ratios of different interneuron subtypes. All resulting p-values (n = 498 tests) were corrected jointly using the Benjamini–Hochberg procedure^92^, rather than within each category separately, to ensure a conservative multiple testing correction. Correlation coefficients and p-values are shared in **Table S4**. Selected region pairs were visualized using scatterplots and heatmaps.

#### Cell type proportion regressions

We sought to evaluate age and sex effects only on cell types that were well-ascertained, with a median fraction > 1% across all samples. Cell-type abundance denominators were selected according to the biological comparison of interest. Broad cell classes were quantified relative to all nuclei, whereas finer neuronal populations were quantified relative to matched neuronal reference populations to reduce sensitivity to variation in gray-/white-matter sampling (see **Table S5** for the specific denominator used in each analysis). We limited analysis of neurons to gray matter regions only (ic excluded). Since not all samples had OPC subcluster labels, we restricted the analysis of subcluster 0 and 1 abundances to the n=397 samples that contained at least 10 nuclei from both cluster 0 and cluster 1.

As is common for postmortem human brain tissue, the total number of nuclei recovered per sample varied substantially. This variability matters because it affects how precisely cell type proportions can be estimated: a proportion based on a few hundred nuclei is much noisier than one based on tens of thousands. Rather than treat every sample’s proportion as equally reliable, we used beta-binomial regression, which models the counts directly and lets samples with more nuclei carry proportionally more weight. Specifically, for each regression, we modeled the log-odds of the cell type(s) of interest (n_nuclei) relative to a reference group (n_other) — either all remaining nuclei or a biologically chosen subset. Because proportions are bounded between 0 and 1, we use a logit link to map the expected proportion onto an unbounded scale, which lets us fit and interpret the model with standard regression tools.

For each cell type of interest, we fit a beta-binomial regression mixed effects model using glmmTMB^93^ (RRID:SCR_025512). Donor and village were modeled as random effects, while the imputed sex, age (in decades) and brain region were incorporated as fixed effects. Analysis of cortical neurons was limited to DFC, so we omitted the fixed effect of brain region as well as the random donor effect, since each donor only had one observation in DFC.

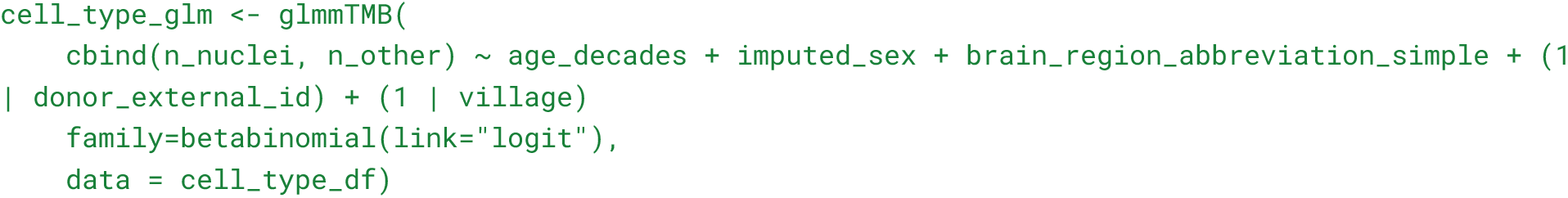

From these models, we extracted point estimates and 95% confidence intervals of the coefficients (β) corresponding to age and sex to quantify their effects on cell-type abundances while controlling for other technical and biological covariates. These effect sizes operate linearly on the log-odds: exponentiating β thus yields the odds ratio, which one can interpret as the fold-change of the ratio of interest (n_nuclei / n_other) with respect to a unit change in the covariate (for age, this is the fold change per decade). For rarer cell types (with proportion <10%) the odds ratio closely approximates the fold change in the proportion (or risk ratio) of that cell type ^94^. Statistical significance was assessed using Wald tests, and p-values were adjusted for multiple comparisons across cell types and covariates of interest (age and sex; n = 54 tests) using the Benjamini–Hochberg procedure, with a false discovery rate (FDR) threshold of 0.01. To reduce the influence of sparse sampling in age- and sex-association analyses of OPC subclusters (most comprising <0.12% of all nuclei), we restricted analyses to samples containing at least one nucleus from the subcluster of interest. All regression results are presented in **Table S5**.

Residuals were computed for each sample as the difference between the observed cell type proportion and the predicted proportion from the model based on fixed effects (age, sex, brain region) only.

To visualize the age effect on cell type abundance in the original proportion scale, we generated model-based predictions from the beta-binomial GLMM across the observed age range, for each brain region, while holding categorical covariates (sex) constant at the mode. Predictions were obtained on the response scale with random effects excluded to obtain the population-level fit, and plotted with 95% confidence intervals.

Population-level predicted cell type abundances were computed using emmeans^95^ (RRID:SCR_018734). Predictions were obtained at selected ages (30 and 80 years) for each brain region, averaging over the levels of sex, and 95% confidence intervals were derived on the response scale. Differences between ages were estimated using contrasts of the back-transformed predictions.

For OPC abundance (relative to all nuclei sampled), predicted proportions decreased significantly with age, from 0.0641 (95% CI: 0.0577–0.0705) at age 30 to 0.0406 (95% CI: 0.0376–0.0436) at age 80. This corresponds to an absolute difference of −0.0235 (95% CI: −0.0303 to −0.0166), an approximate 36.7% reduction in OPC abundance over fifty years.

GABAergic TAC3–PLPP4 interneuron abundance (relative to SPNs) also declined significantly with age, with predicted composition decreasing from 0.0319 (95% CI: 0.0292–0.0345) at age 30 to 0.0259 (95% CI: 0.0244–0.0275) at age 80. This corresponds to an absolute difference of −0.00592 (95% CI: −0.00901 to −0.00284), an approximate 18.6% reduction over fifty years.

#### Age-region interactions in OPC loss

As an additional analysis, in order to determine whether OPC loss is more pronounced in any of the regions sampled, we performed a beta-binomial regression that also incorporated an age-by-region interaction term. There were no statistically significant age-by-region interactions.

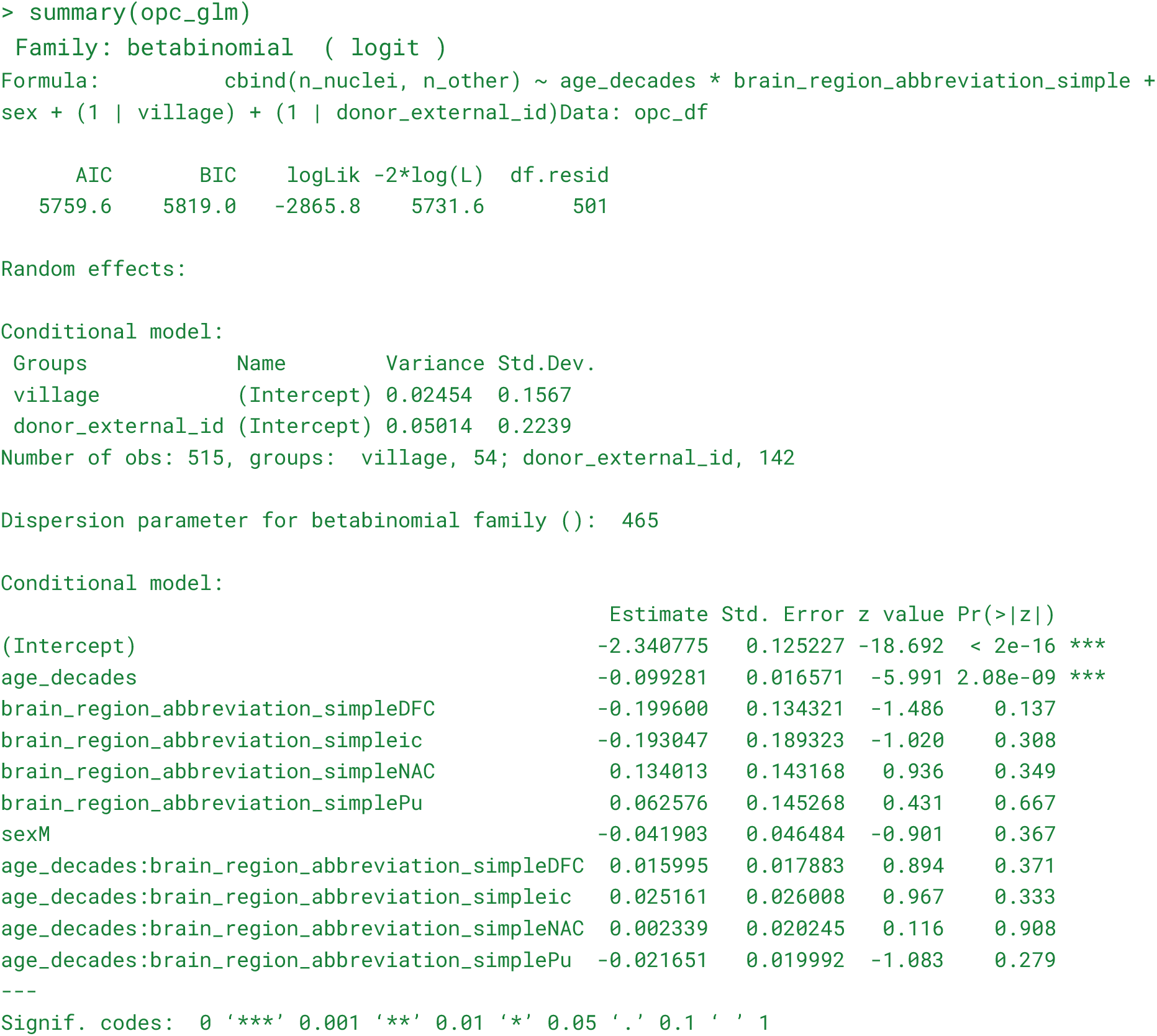

#### Evaluating effects of technical metadata

One important question is whether glial correlations might arise artifactually due to donor-level differences in tissue quality. We identified postmortem interval (PMI, available for all donors) and RNA quality score (RQS, available for 124/142 donors in the core donor set that were not CTP outliers, see **Table S1**) as important donor-level features capturing information about overall tissue quality. Additionally, for each sample, we took the median number of UMIs per projection neuron (normalized to the village median to account for variability in sampling depth between batches) as a metric for sample-level tissue quality. We found that both RQS and pH (only available for 40/142 donors and therefore excluded from the analysis) were significantly correlated with normalized_median_umis_per_neuron, but PMI was not (see **Figure S19**).

For each sample, we regressed the cell type proportion of every glial cell type against PMI, RQS, and normalized median UMIs per projection neuron. We took the residuals from these linear fits to represent the component of the cell type proportion that is not explained by a linear relationship with these donor- or sample- level tissue quality metrics. We found that even after correcting for the effects of tissue quality, the strong correlations of astrocyte, OPC, and microglia (residual) cell type proportions between gray matter regions persisted, and the correlations of (residual) oligodendrocyte fraction were similarly robust within the striatum (See **Figure S5A,B**)

Moreover, the significant negative age effects on OPC, TAC3, and SST interneuron fractions persist when these covariates are included in betabinomial regression frameworks, supporting that these effects are not generated artifactually by age associations with tissue quality.

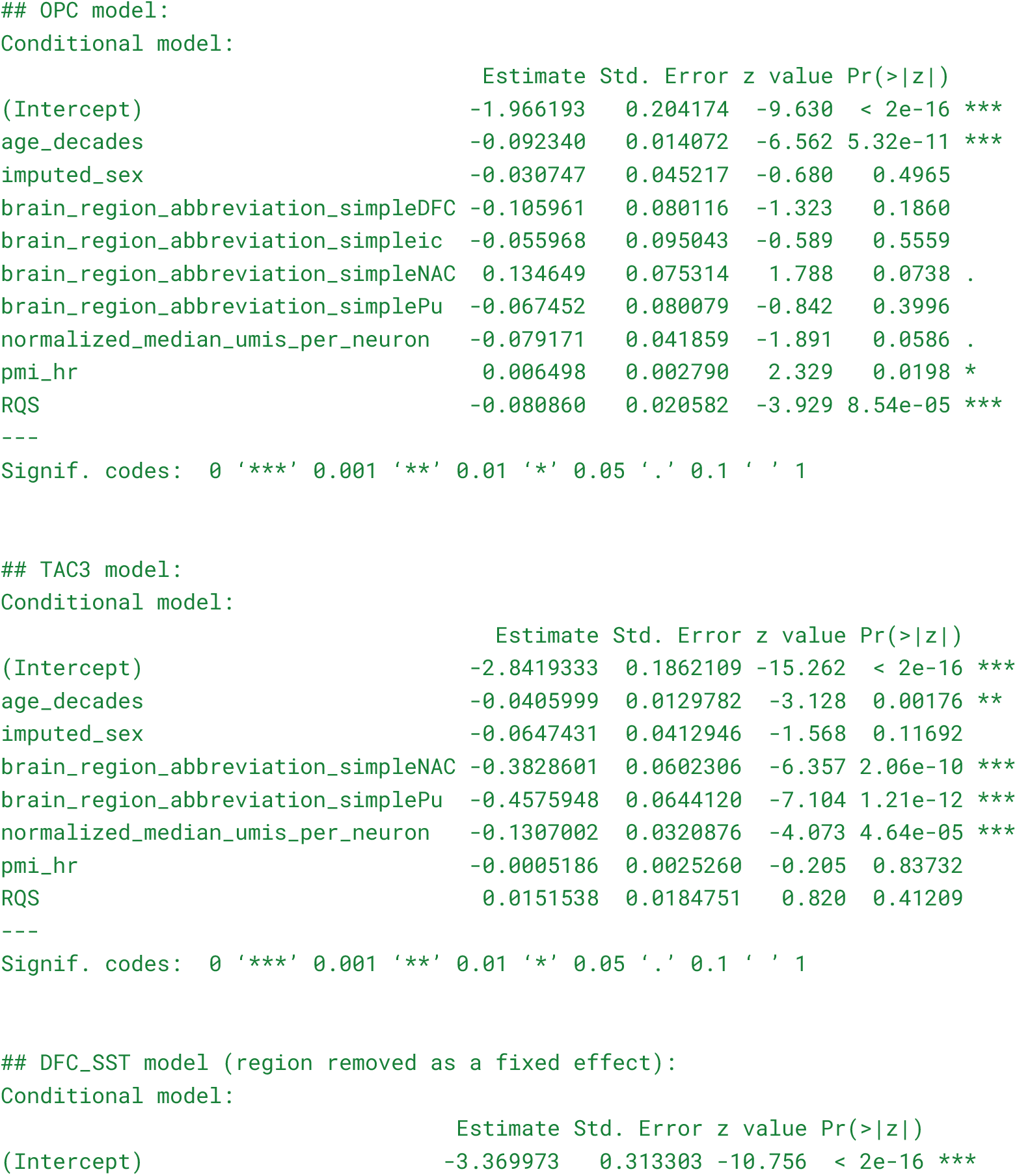
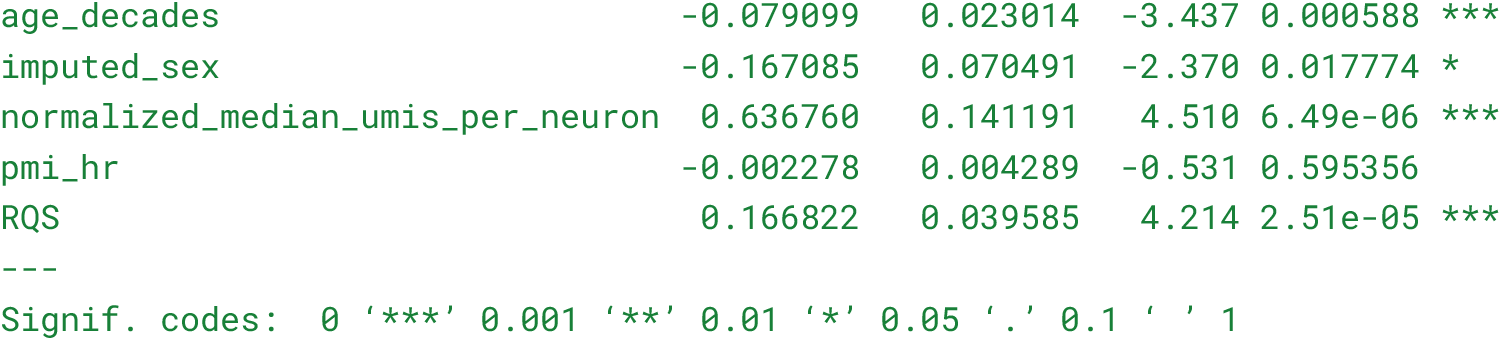

#### Bulk RNA-seq cell-type proportion estimation

To estimate cell type proportions using bulk RNA-seq data we used a tree-based machine learning approach ^96^. The model was trained using simulated “pseudobulk” data ^97^ constructed from the single-nucleus RNA-seq reference data in this study.

#### Cell-type-specific expression profiles

Cell-type-specific expression profiles were generated using our annotated single-nucleus RNA-seq data. We considered a panel of eight main cell types: SPN, PTHLH–PVALB interneuron, TAC3–PLPP4 interneuron, astrocyte, oligodendrocyte, OPC, microglia, and vascular cells.

Using data from caudate, counts were summed to form donor × cell type metacells. These metacells were normalized to sum up to 10⁶, and used as donor × cell type “expression profiles” when generating synthetic pseudobulk training data.

#### Cell-type proportion sampling distribution

To establish a realistic cell-type proportion space, cell type proportions were drawn from a Dirichlet distribution defined by a single concentration parameter (α₀) shared across cell types and by cell-type-specific means estimated from the observed reference donor compositions. No individual donor’s composition was used as a training label.

The concentration parameter α₀ was obtained by inverting the observed across-donor variance of each cell type, taking the median across types, and dividing by a default factor of 3 to widen the simulated range (α₀ = 38.3); this widening is necessary because gradient-boosted trees cannot predict outside the range of labels seen during training, and we verified that the simulated compositions bracketed the observed proportions of every donor for seven of eight cell types.

#### Constructing the training data

For training, we created synthetic pseudobulk mixtures using the donor × cell type expression profiles and cell-type proportions drawn from the Dirichlet distribution described in the two sections above.

For each synthetic pseudobulk training sample, a cell-type composition **p** was drawn from the distribution described above. For each of the eight cell types independently, one donor was selected at random and that donor’s normalized expression profile for that cell type was used. This separation of cell-type proportion sampling from gene-expression profile sampling was done to ensure that no training sample exposed specific donor cell-type proportion / gene expression relationships, while simultaneously creating synthetic pseudobulk training data that represented realistic cell type proportions and gene expression profiles.

Cell type profiles were combined using RNA-content weights w ∝ **p·m**, where **m** is the mean number of UMIs per nucleus for each cell type, estimated from the same reference; accounting for mRNA-content differences when simulating pseudobulk follows Dietrich et al. (2022) ^98^, who use spike-in- or gene-count-derived scaling factors instead. This accounts for the fact that bulk expression represents each cell type in proportion to its transcript contribution rather than its cell fraction, which is the prediction target. Multiplicative lognormal noise (σ = 0.15) was applied per gene. A total of 12,000 training samples were generated (fixed random seed 0), with profiles required to derive from at least 25 nuclei; profiles from 6 of the 48 donors were reserved to build 2,000 validation mixtures used solely for early stopping of training the model, so that no validation profile appears in any training mixture. These six donors’ compositions still contribute to the Dirichlet prior, and they remain in the evaluation set.

#### Constructing the Model

One gradient-boosted regressor (LightGBM; RRID:SCR_021697) was trained per cell type to predict that type’s proportion from expression normalized to counts per 10,000 and log1p- transformed. Hyperparameters were fixed across all cell types and experiments (num_leaves 15, max_depth 4, min_child_samples 150, colsample_bytree 0.4, subsample 0.8 applied every iteration, L2 penalty 20, learning rate 0.02, up to 6,000 trees with early stopping after 150 rounds without improvement in validation RMSE). A 60-configuration random search gave no improvement over this default on real bulk, and differed only by noise. Fixing the configuration also prevents model selection from confounding the comparisons reported below. Increasing the number of simulated training mixtures from 10,000 to 40,000 did not improve performance (mean Spearman correlation 0.395, 0.400, 0.398 and 0.393 at 10,000, 12,000, 20,000 and 40,000 samples), consistent with the combinatorics of cross-donor mixing saturating well below that range; 12,000 was used throughout.

#### Bulk RNA-seq data processing

In bulk RNA-seq data from tissues, intronic reads (i.e. reads aligning to genes’ intronic sequences) come from nascent nuclear transcripts and are therefore closer in composition to the nuclear-RNA reference on which the models had been trained. Models were therefore applied to the intronic component of the bulk RNA-seq libraries, i.e. to reads that aligned to genes’ intronic sequences – about 20% of all reads. Intronic read counts were computed per gene and per donor as gene-body counts minus exonic counts, from quantifications available for all libraries; both types of counts were computed by FeatureCounts (RRID:SCR_012919).

#### Per-gene calibration

Bulk RNA-seq expression levels were rescaled per gene before prediction to match the gene-level means of the training data ^99^. For each donor, a per-gene scale factor was computed as the ratio of mean expression across the remaining donors to the mean of the training pseudobulk, normalized by its median and clipped to [0.05, 20]; the donor’s expression was divided by this factor and renormalized. It uses no cell-type proportions for any donor, and the target donor’s own expression is excluded from its own scale factor.

#### Prediction and evaluation

Per-cell-type abundance estimates were produced independently and are reported after renormalization to sum up to 1 (clipping at zero). Performance was assessed as the Spearman correlation between bulk-RNA-seq-estimated and snRNA-seq-estimated proportions across the 48 donors, computed separately per cell type, with p-values corrected across the eight types by the Benjamini–Hochberg procedure. Calibration was reported separately as the regression slope of prediction on truth.

#### RNAscope Image Analysis

Whole-slide images were analyzed using QuPath v0.7.0 (RRID:SCR_018257) across three probe panels: PDGFRA and SLC1A2 in caudate and PDGFRA in putamen. As each section was hybridized with a single probe, no spectral unmixing was required. DAPI channel acquisition settings were identical across all sections of all panels; TSA 570 exposure was optimized separately for each of the two probes and held constant across all sections stained with that probe. Nuclear detection is therefore directly comparable across the full dataset, while probe-signal comparisons are made within probe groups. Channel display range was adjusted only for visualization during manual annotation; all detection and measurements were computed from raw pixel values and were unaffected by display settings.

Regions of interest were positioned entirely within the caudate nucleus or putamen and identified using gross anatomical landmarks such as the lateral ventricle and internal capsule. Adjacent white matter and the ventricular wall were not included. Intrastriatal fiber bundles falling within the scan region were retained, as comparable structures are present in the tissue dissections used for snRNA-seq. For each section, the entire ROI was annotated, and areas containing tissue folds, tears, detachment, out-of-focus tissue, or obvious fluorescent artifacts were manually excluded from cell detection. Donors in which tissue quality and availability precluded reliable analysis were excluded, leaving 11 donors for the caudate PDGFRA panel, 11 donors for the caudate SLC1A2 panel, and 8 donors for the putamen PDGFRA panel (**Table S12**).

Nuclear detection was performed on the DAPI channel using QuPath’s watershed cell detection, with parameters held constant for every section of every panel: requested pixel size 0.5 µm, background radius 8.0 µm, median filter radius 0 µm, Gaussian sigma 1.5 µm, minimum nuclear area 15 µm^2^, maximum nuclear area 400 µm^2^, intensity threshold 100, watershed post-processing enabled, and cell expansion 5.0 µm. A saved QuPath script was applied across each project so that parameters were identical across sections; no section-specific adjustment was performed. In total, 982,864 nuclei were analyzed across 1,305 mm^2^ for caudate PDGFRA, 1,035,880 nuclei across 1373 mm^2^ for caudate SLC1A2, and 346,734 nuclei across 507 mm^2^ for putamen PDGFRA.

Probe-positive and probe-negative nuclei were classified using Random Trees object classifiers trained on nucleus-compartment measurements only: TSA 570 minimum, mean, maximum, standard deviation, range, and sum; DAPI mean; and nuclear area, circularity, and eccentricity. Restricting measurements to segmented nuclear signals provides a conservative estimate and the closest available analogue to the nuclear transcriptome sampled by snRNA-seq. Training data were generated by manually labeling individual nuclei with the QuPath Points tool, using a minimum of 50 probe-positive and 50 negative nuclei per section. Annotation was performed blinded to donor identity, donor metadata, and snRNA-seq cell type proportions. Positive examples were selected to span the observed range of signal intensity, and negative examples were drawn preferentially from nuclei with sparse or ambiguous puncta rather than from only unlabeled nuclei.

Shared and independent classifiers were trained for each probe panel (**Table S12**). As background and autofluorescence signals varied largely between postmortem specimens, classifier assignment was determined per section by initial signal assessment performed prior to quantification, on image and signal quality alone; assignment was therefore made independently for each probe and region, and a given donor could be classified using a shared classifier in one panel and a separately trained classifier in another. Sections with similar background and signal-to-noise characteristics were analyzed using a common classifier trained from pooled annotations. Sections identified as having markedly elevated or heterogeneous background signal were classified using separately trained classifiers built from annotations of the corresponding section alone ^42,100^, both to prevent misclassification of those sections and to prevent their atypical intensity distributions from impeding shared-classifier performance. All classifier definitions are provided in the deposited QuPath projects.

Cell type proportions were calculated per donor as the number of probe-positive nuclei divided by the total DAPI^+^ nuclei detected within the analyzed region. PDGFRA and SLC1A2 were assayed on near-serial caudate sections within 60 µm from the same tissue block, and proportions for the two markers therefore derive from independent sections and independent denominators. All analysis parameters and classifiers were finalized prior to comparison with snRNA-seq proportions. QuPath project files, classifier definitions, annotation sets and analysis scripts are available.

#### Differential expression

To detect sex-biased and age-dependent gene expression, we summed UMI counts of all assignable single cells for each distinct combination of donor, brain region, and 10x chemistry, generating a pseudobulk (observation-by-gene) matrix at the donor–region–chemistry level for each cell type. Fixed-effect covariates considered for downstream analysis included imputed sex, age, PMI, five genetic principal components (PC1–PC5), 10x chemistry (variable “single_cell_assay”), brain region, and biobank, as well as mean single-cell quality metrics per sample: percent of reads that are intronic (variable “pct_intronic”), the fraction of UMIs removed by CellBender (variable “frac_contamination”), and the z-score of the log10 number of nuclei captured (variable “z_log10_nuclei”). Additionally, donor and village are modeled as random effects.

We evaluated these potential covariates using two complementary approaches. First, we generated multidimensional scaling (MDS) plots in Glimma^101^ (RRID:SCR_017389) for each cell type to visualize relationships among samples and identify variables explaining major axes of variation. This qualitative assessment highlighted covariates contributing visible structure in the data, such as donor identity, brain region, and 10x chemistry. Second, we quantified the contribution of each covariate to gene expression variance using the variancePartition^102^ framework.

Based on these assessments, we selected covariates explaining substantial variation for inclusion in the dream^103^ differential expression models. dream, an extension of limma-voom (RRID:SCR_010943), fits linear mixed models with multiple random effects to account for repeated measurements. Donor and village were modeled as random intercepts, while the selected covariates were incorporated as fixed effects.

For each cell type, we filtered observations to remove donor–region–chemistry samples with low library size that were likely to contribute more noise than signal. Library size filtering was performed by computing the mean and standard deviation of total UMIs per sample and retaining only samples with at least mean − 1.96 SD total UMIs. Genes were then filtered to remove lowly expressed features. Counts were transformed to counts per million (CPM) using edgeR^104^ (RRID:SCR_012802), and we retained genes where at least 10% of observations had CPM ≥ 1.

Differential expression was tested at two levels of granularity. First, to estimate the average effect of sex or age on each cell type across brain regions, we fit a region-averaged model with age as a single fixed-effect term and region included as a categorical covariate. This model was specified in dream as:

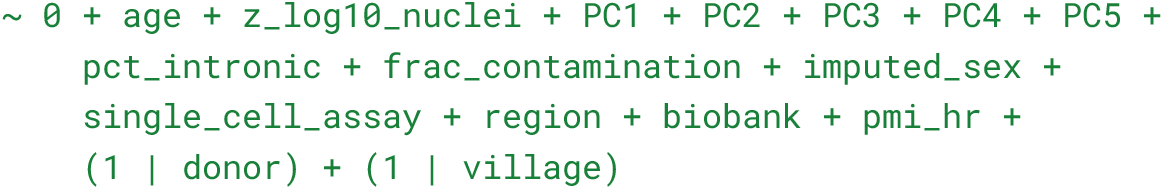

To assess region-specific heterogeneity in age effects, we fit a second model in which age effects were allowed to vary by region using region-specific age terms. Using caudate as the reference region, region-specific age effects were encoded as separate fixed-effect terms (for example, age_regionCaH, age_regionDFC). The corresponding dream formula was:

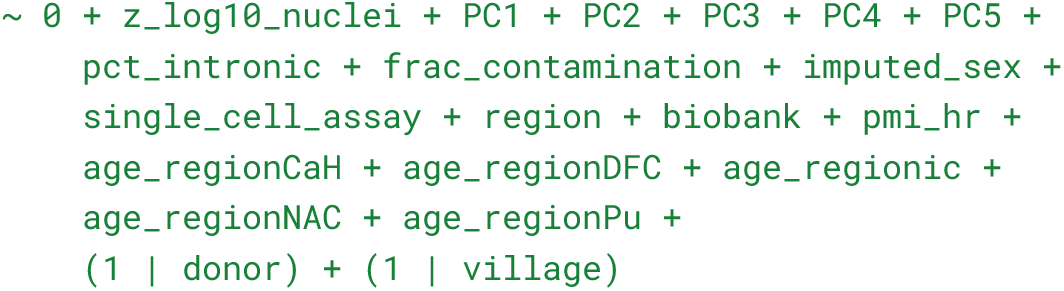

This formulation allowed us to estimate region-specific age effects from a single model fit and to compare the magnitude and direction of age-associated differential expression for each gene across brain regions.

#### Differential expression analyses

Age-associated differential expression was assessed using precomputed results from linear modeling frameworks treating age as a continuous variable, as described in the differential expression section. Results included gene-level effect size estimates, test statistics, and FDR- adjusted significance values. Effect size uncertainties derived from model outputs were used in downstream analyses requiring standard error estimates. Analyses were performed both with brain regions combined and with region-specific models, as noted in individual analyses below.

#### BMI and smoking status analysis

We performed a further exploratory analysis of BMI and smoking status to determine the impacts of these phenotypes on gene expression. We extended the regression analysis described above to include BMI as a continuous variable and smoking status as a binary variable. BMI information was available for 115 donors and smoking status for 96 donors, with 77 donors having complete information for BMI, smoking status, age, and sex. Because differences in sample size affect statistical power to detect differential expression, we restricted comparisons among the four phenotypes to these 77 donors. This allowed us to compare the numbers of differentially expressed genes associated with BMI, smoking, age, and sex without unfairly penalizing phenotypes with smaller sample sizes.

In this subset of 77 donors, few genes showed statistically significant associations with BMI or smoking status (**Table S13**). The limited number of detected associations may reflect insufficient statistical power in our cohort rather than an absence of effects on gene expression.

Because BMI and smoking status were available for only a subset of donors, including either as a covariate in the primary differential expression analysis would reduce power to detect age effects. We therefore separately tested whether accounting for BMI or smoking added information to the regression model and altered estimated age effects. For each phenotype, we restricted the analysis to donors with the corresponding metadata and fit nested regression models that differed only by inclusion of BMI or smoking status. For example, for BMI:

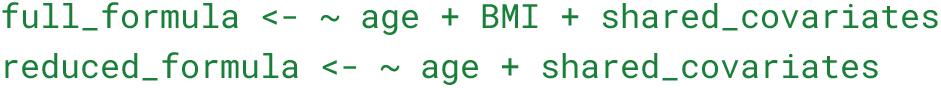

We formally compared the nested models and also compared the estimated age effect between them. Inclusion of BMI or smoking did not substantially change age-effect estimates in any cell type. The estimates were highly concordant across cell types and regions: correlations were ≥ 0.987 after including BMI and ≥ 0.978 after including smoking status.

#### Cell-type aging effect-size correlations

Concordance of age-associated transcriptional effects across cell types and brain regions was evaluated using pairwise comparisons of gene-level effect sizes in R using cor.test. Genes were aligned across conditions, and comparisons were restricted to genes showing evidence of age association in at least one condition (FDR < 0.05). Spearman’s rank correlation was used. Correlation strength and direction were summarized using signed metrics derived from correlation coefficients. Selected comparisons were visualized using scatterplots.

#### Gene k-means clustering across cell types

A matrix of gene-level age-associated effect sizes estimated across all five brain regions was constructed across analyzed cell types. To arrive at a set of informative genes for this analysis we filtered to genes expressed at 10 transcripts per million (for the cell type being analyzed; in either striatum or cortex) with an FDR < 0.01 and an absolute fold change (per decade) > 1.05 in at least one cell type. This left us with 2,478 input genes. These genes were clustered based on shared age-related expression patterns using k-means clustering (stats::kmeans) with Euclidean distance, a fixed random seed (42), 200 random starts, and a maximum of 20 iterations. Effect sizes were z-score-scaled across cell types prior to clustering. The number of clusters was guided by silhouette-based evaluation (evaluating k = 10-30). Clustered gene patterns were visualized as heatmaps.

#### TRADE analysis by cell type

Transcriptome-wide impact of aging was quantified using TRADE^60^, which integrates gene-level effect sizes and associated uncertainties to estimate global transcriptional perturbation. Our primary TRADE analyses (**Figure 4I**) were performed separately for each cell type using effect sizes estimated across all five brain regions. Autosomal genes were included in primary analyses, with sex chromosome genes analyzed separately. The primary outputs used in downstream analyses were the estimated transcriptome-wide impact values per cell type. For region-level TRADE analyses (**Figure S13E**), analyses of the internal capsule were restricted to non-neuronal cell types (astrocytes, OPCs, oligodendrocytes, and microglia). Because the internal capsule is a white-matter region, neuronal populations were sparsely and inconsistently sampled, likely reflecting variable inclusion of adjacent striatal gray matter during dissection.

#### TRADE following downsampling

Cells were downsampled separately within each donor, brain region, and cell type. Within each donor-region combination, we defined the target cell count using lower-abundance cell types relevant to our DE analyses (interneurons and OPCs) thereby avoiding reduction to the very lowest-ascertained populations (such as microglia and vascular cells) while ensuring that interneurons and OPCs were sampled equivalently relative to the more abundant neuronal and non-neuronal classes. Cell types above the target were randomly downsampled, cell types below the target were retained in full, and donor-region combinations lacking one or more required target cell types were excluded.

#### Gene set enrichment analyses

Rank-based gene set enrichment analysis was performed using fgsea (RRID:SCR_020938) in R. Genes were ranked by age-associated t-statistics. Curated gene sets from the MSigDB (RRID:SCR_016863) Human C5 collection (biological process, molecular function, and cellular compartment) were tested using permutation-based enrichment with multiple hypothesis correction. Gene sets used for visualization were selected from leadingEdge genes of statistically significant (padj < 0.01) GO categories related to the selected themes (e.g., GOBP_REGULATION_OF_SYNAPTIC_PLASTICITY, GOBP_DENDRITIC_SPINE_MORPHOGENESIS, GOBP_G_PROTEIN_COUPLED_DOPAMINE_RECEPTOR_SIGNALING_PATHWAY, GOBP_CALCIUM_ION_TRANSPORT; **Table S7**).

### RNA based age prediction

#### Data preprocessing and feature selection

Single-nucleus RNA-sequencing data were aggregated to generate pseudobulk expression profiles defined at the donor × cell type × region level. For donors with multiple libraries or chemistries, raw UMI counts were summed across nuclei to yield a single expression profile per donor within each cell type and region. Only autosomal genes (chromosomes 1–22) were retained for analysis; genes located on sex chromosomes or mitochondrial DNA were excluded prior to further filtering.

Samples with low library size were removed as described in the differential expression analysis. Genes were filtered using the same criteria applied in the differential expression framework, including removal of lowly expressed features based on counts per million (CPM).

#### Model training and prediction

For each modeling iteration, raw count data were transformed to log counts per million (log- CPM) using edgeR. Gene expression features were standardized by z-score transformation, computed separately for each gene using the mean and standard deviation estimated from the training donors only. The model was fit to these standardized expression values, yielding a regression coefficient for each gene and a fitted intercept. For each held-out donor, raw counts were transformed to log-CPM and then to z-scores using the gene-specific means and standard deviations learned from the training data. Donor age was then predicted as the dot product of these z-scored expression values and the fitted regression coefficients, plus the model intercept. Thus, held-out donors contributed neither to model fitting nor to estimation of the expression-scaling parameters, mirroring the procedure that would be used for prediction in an external dataset.

#### Repeated age-stratified cross-validation

A central concern when evaluating an age-prediction model is ensuring that the donors used to evaluate prediction accuracy do not also contribute to fitting the model. We therefore explicitly separated donors into training and held-out test sets. Training donors were used to estimate model weights and select model parameters, whereas the chronological ages of the held-out test donors had no influence on model fitting. Predictions for these donors were generated only after the model had been trained. In initial cross-validation analyses, however, we found that prediction accuracy could vary depending on the particular donors assigned to the training and test sets, especially for cell types represented by relatively few donors. We therefore used a repeated age-stratified cross-validation procedure to reduce sensitivity to any single partition of the data.

For each cell type by region model, donors were divided into five age bins based on age quintiles, and approximately 20% of donors within each bin were randomly withheld to form a test set, preserving the overall age distribution. The remaining donors formed the training set. This procedure was repeated 200 times, with a new stratified split sampled at each iteration. Thus, each donor was repeatedly evaluated as a held-out sample against many different sets of training donors and received approximately 40 out-of-fold predictions on average.

Within each iteration, only the training donors were used to fit the age-prediction model. Chronological age, expressed in decades, was predicted using elastic net regression, with α = 0.5 and a Gaussian error model. The regularization parameter λ was selected using random 10- fold cross-validation within the training set to minimize mean absolute error (MAE). Model coefficients and the intercept were then estimated from the training donors using the selected λ. Only after model fitting was complete were the learned model weights applied to the withheld test donors to generate predicted ages.

For each donor, final predicted age was defined as the mean of all out-of-fold predictions obtained across iterations in which that donor was withheld. Model performance, including Pearson correlation between predicted and chronological age and MAE, was calculated from these aggregated donor-level out-of-fold predictions.

#### Bias correction and residual age

Across models, predicted ages exhibited regression-to-the-mean shrinkage, with younger donors predicted older and older donors predicted younger. The magnitude of this shift tracked the number of age-associated differentially expressed genes available to each model (**Figure S15B,C**). Uncorrected residual age therefore showed a shared age-dependent bias across models (**Figure S15D**). To account for this systematic age-dependent bias, we fit a generalized additive model (GAM) for each cell type by region using aggregated out-of-fold predictions as a smooth function of chronological age. The model used a cubic regression spline (k = 5) which was estimated by restricted maximum likelihood, and employed a scaled t distribution to provide robustness to outliers.

Bias-corrected predicted age was defined by removing the estimated age-dependent deviation from the identity relationship between predicted and chronological age, thereby eliminating regression-to-the-mean effects. Residual age was defined both as the raw difference between predicted and chronological age and as the bias-corrected difference. The correction eliminated the age dependence of the residuals and removed the correlation driven by this shared bias (**Figure S15E**). Both forms were used in downstream analyses, with bias-corrected residual age used to assess coordination of transcriptional variation across cell types and regions.

#### Covariate adjustment of prediction residuals

To test whether correlations among cell-type-specific age residuals reflected variation in sample quality or sequencing depth, we adjusted for postmortem interval (PMI), RNA quality score (RQS), and library size. For each cell type by region, donors with missing covariate values were excluded, library size was log_10_-transformed, and the three covariates were standardized by z- score transformation. We then fit a linear model with mean out-of-fold predicted age as the response and chronological age and the three standardized covariates as predictors. Including chronological age allowed covariate effects to be estimated while accounting for age-associated variation in predicted age.

Covariate-adjusted predicted age and residual age were defined by subtracting the fitted contributions of PMI, RQS, and log library size from the corresponding unadjusted values. Pairwise Pearson correlations among bias-corrected age residuals were calculated before and after covariate adjustment using the same donor subsets. P values for covariate associations were adjusted using the Benjamini–Hochberg procedure.

After adjusting for PMI, RQS and log library size, the pairwise correlations between age- prediction residuals were generally slightly weaker, but remained positive and of similar magnitude (r = 0.27–0.39 after adjustment, compared with 0.26–0.40 before adjustment in the same donor subset; **Table S14**). The coordinated age-prediction residuals across cell types remained after accounting for these covariates.

#### Age prediction with DE feature selection

To test whether incorporating prior information about age-associated genes improved prediction, we repeated the analysis using only genes significantly associated with age (FDR ≤ 0.05) in the corresponding cell type and region–specific differential expression analysis. These genes served as a predefined feature set, and models were fitted using ridge regression (elastic net α = 0). Predictive accuracy was modestly higher than in the primary models using all expressed genes (median model-level mean absolute error, 6.3 versus 6.6 years) (**Figure S16A,B**), and the correlation structure among cell-type-specific age residuals was preserved (**Figure S16C–E**), supporting the robustness of the principal findings to the choice of feature set and regularization. Differences in predictive accuracy by cell-type-region are listed in **Table S15**.

#### DE-selected features in independent cohort

To test whether age-associated gene features identified in our cohort generalized to an independent dataset, we repeated the age-prediction analysis in the Ling et al. cohort. For each cell type, candidate features were restricted to age-associated DE genes identified in our primary cohort, and elastic-net models were trained and evaluated in the Ling et al. data using the same cross-validation framework. The models predicted chronological age with comparable accuracy across cell types (**Figure S16F,G**), and correlations among cell-type-specific age residuals were preserved (**Figure S16H-J**), indicating that age-associated expression features identified in our cohort capture reproducible inter-individual variation in an independent cohort.

#### Cross-cohort transfer of model weights

To test whether an age-prediction model trained in one cohort could be applied directly to another without refitting, we used models trained on our primary cohort donors and restricted to genes that were associated with age (FDR < 0.05) to predict donor ages in the independent Ling et al. dataset. For each cell type, Ling et al. expression values were transformed to log CPM and standardized using the gene-specific means and standard deviations learned from the corresponding training data in our cohort. Predicted age was then calculated using the fitted elastic-net coefficients and intercept from our cohort. As a reference, we independently trained age-prediction models in the Ling et al. data using the same analysis framework and age- associated genes identified within that cohort.

Because the model intercept reflects the mean age of the training cohort, differences in cohort age distributions can introduce a systematic offset when coefficients are transferred. We therefore evaluated both the unmodified transferred model (“raw”) and an intercept-corrected model in which the original intercept was replaced by the mean chronological age of the Ling et al. cohort. Transferred models produced age predictions similar to those from models trained directly in Ling et al., particularly after intercept correction (**Figure S16K-M**).

### eQTL analysis

#### Discovery

To generate eQTLs, we used a workflow combining parts of previous pipelines^25,105^. We summed UMI counts of all assignable single cells included in nucleus selection for each distinct combination of donor, brain region, and cell type to create pseudobulked (donor-by-gene) expression matrices at the region cell-type level. Pre-processing of expression data included summing the gene expression per gene defined by the GENCODE v43 gene models and removing the lower 50% of expressed genes, leaving 18,984 to 19,056 genes per brain region cell type matrix. Each matrix was then normalized per-donor using code derived from the pyQTL^106^ implementation of edgeR CPM, then standardized using the pyQTL implementation of an inverse normal transformation (INT).

The covariates from the differential expression were used for identifying fixed effects. Probabilistic Estimation of Expression Residuals (PEER)^107,108^ was used to generate 10 additional covariates for each brain region × cell type analysis.

Variants included in the analysis were from chromosomes 1-22 and X using the GRCh38 reference. Sites with a minor allele frequency less than 0.05 were excluded from the analysis. For each variant, 90% of the donors were required to have a genotype quality greater than 30, as measured by the GQ quality tag in the VCF. Sites where the HTSJDK^109^ calculated a Hardy Weinberg equilibrium (HWE)^87,88^ p-value below 1e-4 were excluded.

The variant genotypes, normalized gene expression phenotypes, and set of covariates were input into tensorQTL (RRID:SCR_028171)^110^ to generate cis-QTLs using its cis mode with a seed of 777, a default genome-wide false discovery rate (FDR) setting of 0.05, using a setting of one megabase window before and after each transcription start site (TSS) as defined by the GENCODE v43 gene models. For cis-QTL results reaching genome-wide significance, the tensorQTL cis_independent mode was then used to generate conditionally independent cis- QTLs^111^ with the same seed, covariates, and TSS window. A list of all pairwise variant and gene expression summary results, without genome-wide FDR calculation, was generated per matrix by using the same tensorQTL inputs with the cis_nominal mode.

Donor exclusion criteria were evaluated using CTP- and gene-expression–based outlier metrics, consistent with the approach used for differential expression analyses (**Figure S10A**). Data-driven filtering preserved or improved discovery yield without materially altering estimated eQTL effect sizes, whereas metadata-driven exclusions substantially reduced power (**Figure S10B**).

#### K-means clustering

We compiled the union of SNP–gene pairs that were significant (q < 0.01) in at least one cell type–region combination. For each gene, we then selected a single lead cis-eQTL variant, defined as the variant with the largest absolute effect size across cell type–region combinations. Effect directions were reoriented so that the strongest effect for each gene was positive. The effect sizes of these lead variants across 17 region-stratified cell-type profiles defined the cross-context effect profile for each gene. Missing effect sizes, arising from genes not tested in specific cell types, were imputed as zero. Genes were then clustered based on these effect-size profiles using k-means.

We selected K = 13 as the smallest value that resolved distinct, biologically interpretable effect-size patterns without merging them. For visualization, cell types were displayed in a fixed biologically informed order (for example, SPN subtypes adjacent to each other), and clusters were arranged to group related patterns. Within clusters, genes were ordered by hierarchical clustering of effect-size profiles (correlation distance, average linkage) to highlight internal structure.

#### Validation of cell-type-specific eQTL effects

To formally evaluate whether cell-type-specific eQTL patterns identified by k-means clustering reflected differences in genetic effect rather than limited power in comparator cell types, we tested each cluster-associated gene-variant pair using a linear mixed-effects model with a genotype-by-group interaction. The interaction term tested whether the genotype effect in the target cell type group differed from the pooled effect in the other cell types. Genes expressed only in the target group were classified as not testable across cell types, whereas genes expressed in at least one comparator cell type were evaluated for a significant interaction.

For each gene-variant pair, inverse-normal-transformed expression from the eQTL pipeline was modeled as:

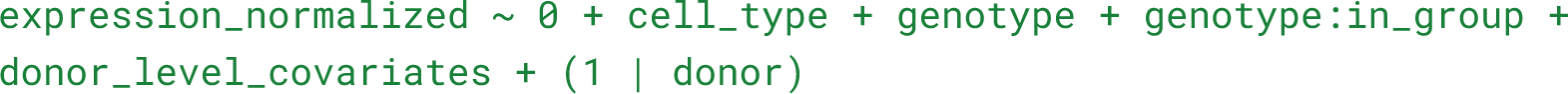

The model included cell-type-specific intercepts and a donor random intercept to account for repeated observations from the same donor; region was additionally included as a fixed effect when observations spanned multiple regions. The genotype:in_group interaction directly tested whether the genotype effect in the target group differed from the pooled effect across comparator cell types. Thus, this analysis tested differences in effect size across cell types rather than inferring specificity from an eQTL being significant in one cell type but not another.

For each k-means cluster, interactions with p-values < 0.05 were considered significant. Gene-variant pairs without an eligible comparator cell type were classified as not testable. This distinguished three outcomes: (1) gene-variant pairs that could not be tested across cell types because the gene was evaluable only in the target group; (2) pairs with a significant difference in genotype effect between the target and comparator groups; and (3) pairs that were evaluable in both groups but showed no significant evidence of a difference in genotype effect.

#### Expression fraction

To quantify the extent to which cell-type-specific eQTLs are driven by cell-type-specific gene expression, we computed median CPM expression across donors for each of the 9,825 eGenes in each of the 17 cell type-region combinations. For each gene-cell type pair, the median was computed from donors with non-NA expression for that gene; gene-cell type pairs with no expressing donors were set to zero. For each of the eight cell-type-specific K-means clusters (clusters 6-13), we identified the cell type with the highest median expression for each eGene and computed the fraction of eGenes whose highest-expressing cell type matched the cell type(s) in which the eQTLs manifested cell-type-specific regulatory effects.

#### Pairwise correlation

To quantify similarity of eQTL architectures across cell types, we computed pairwise Spearman correlations of cis-eQTL effect sizes between all cell-type pairs. For each pair of cell types, we included all SNP–gene pairs significant (q < 0.01) in at least one of the two cell types.

Correlations were assembled into a symmetric matrix and reported as squared correlation coefficients (R²) to facilitate interpretability.

The resulting R² matrix was visualized as a heatmap, with hierarchical clustering applied to both rows and columns to group cell types with similar effect-size profiles.

#### eQTL effect size and genetic constraint

To assess whether evolutionarily constrained genes tend to have smaller eQTL effect sizes, we correlated eQTL effect sizes with LOEUF (loss-of-function observed/expected upper bound fraction) scores from gnomAD^65^ v4.1. Lower LOEUF indicates stronger selective constraint against loss-of-function variants. We restricted the analysis to protein-coding genes with LOEUF scores, using canonical MANE Select transcripts.

For the cluster-level analysis, we used the same lead SNP-gene pairs as the K-means clustering (see above). For each gene, we computed the mean absolute eQTL effect size across cell types in which the eQTL reached significance (nominal p-value below the per-gene empirical threshold from tensorQTL permutations). We then computed Spearman rank correlations between LOEUF and mean effect size within each K-means cluster (K=13), with p-values corrected for multiple testing using the Benjamini–Hochberg procedure. To obtain an overall measure of the LOEUF-effect size association while controlling for cluster membership, we fit a linear regression of mean effect size on LOEUF and cluster (as a categorical covariate) and report the p-value on the LOEUF coefficient.

For the cell-type-level analysis, we used the lead SNP nominated by tensorQTL for each eGene (q < 0.01) independently in each of the 17 cell type-region combinations and computed Spearman correlations between LOEUF and absolute effect size per cell type, with Benjamini– Hochberg correction across cell types.

#### eQTL analyses following downsampling

Cells were downsampled separately within each donor, brain region, and cell type. Within each donor-region combination, we defined the target cell count using lower-abundance cell types relevant to our DE analyses (interneurons and OPCs) thereby avoiding reduction to the very lowest-ascertained populations (such as microglia and vascular cells) while ensuring that interneurons and OPCs were sampled equivalently relative to the more abundant neuronal and non-neuronal classes. Cell types above the target were randomly downsampled, cell types below the target were retained in full, and donor-region combinations lacking one or more required target cell types were excluded.

## Additional Resources

We provide a publicly available web-based portal to query and visualize the data: https://bican.mccarrolllab.org/app/bican.

## Supplemental tables

**Table S2: Donor demographics**. Comprehensive donor-level metadata, including demographic information, clinical and neuropathological diagnoses, tissue source, and cohort inclusion criteria. This table also indicates donor inclusion in downstream analyses.

**Table S4: Cross-region correlations of cell-type proportions and ratios**. Related to **Figure 2**. Spearman’s rank correlations (rho), nominal p-values, and BH-adjusted p-values from cross-region correlations in cell type proportions and cell type ratios. These correlations comprise 498 tests in total.

**Table S5: Beta-binomial regression results for age and sex effects on cell-type abundances.** Related to **Figures 2,3**. Effect sizes, standard errors, test statistics, and p-values for age and sex effects on cell-type abundances estimated using beta-binomial regression. Abundances are defined relative to the indicated denominator (e.g., all nuclei). Age is modeled per decade, and sex is coded as male versus female. Effect sizes (β) are reported on the log- odds scale, with positive values indicating increased relative abundance and negative values indicating decreased relative abundance with the covariate. Odds ratios (exp(β)) are reported as well, and represent the fold-change (per unit change in the covariate) of the cell type of interest with respect to the reference group. Nominal p-values and p-values adjusted for multiple comparisons using the Benjamini–Hochberg procedure are reported. We focused on analysis with BH-adjustment p-values < 0.01.

**Table S7: Age-associated differential expression Gene Set Enrichment Analysis (GSEA) results by cell type.** Related to **Figure 4**. Enriched gene sets, effect directions, and statistical significance across neuronal and glial cell populations.

**Table S8: Donor counts by cell type and brain region.** Related to **Figures 4,6**. Number of unique donors included in differential expression and eQTL analyses for each cell type and brain region.

**Table S13: Differential expression results in donors with complete age, sex, BMI, and smoking status data.** Related to **Figure 4**. Numbers of significantly upregulated, downregulated, and total differentially expressed genes for age, sex, BMI, and smoking status by cell type and brain region, restricted to the 77 donors with complete data for all four phenotypes.

## Supplementary Figures and Tables

**Figure S1.**
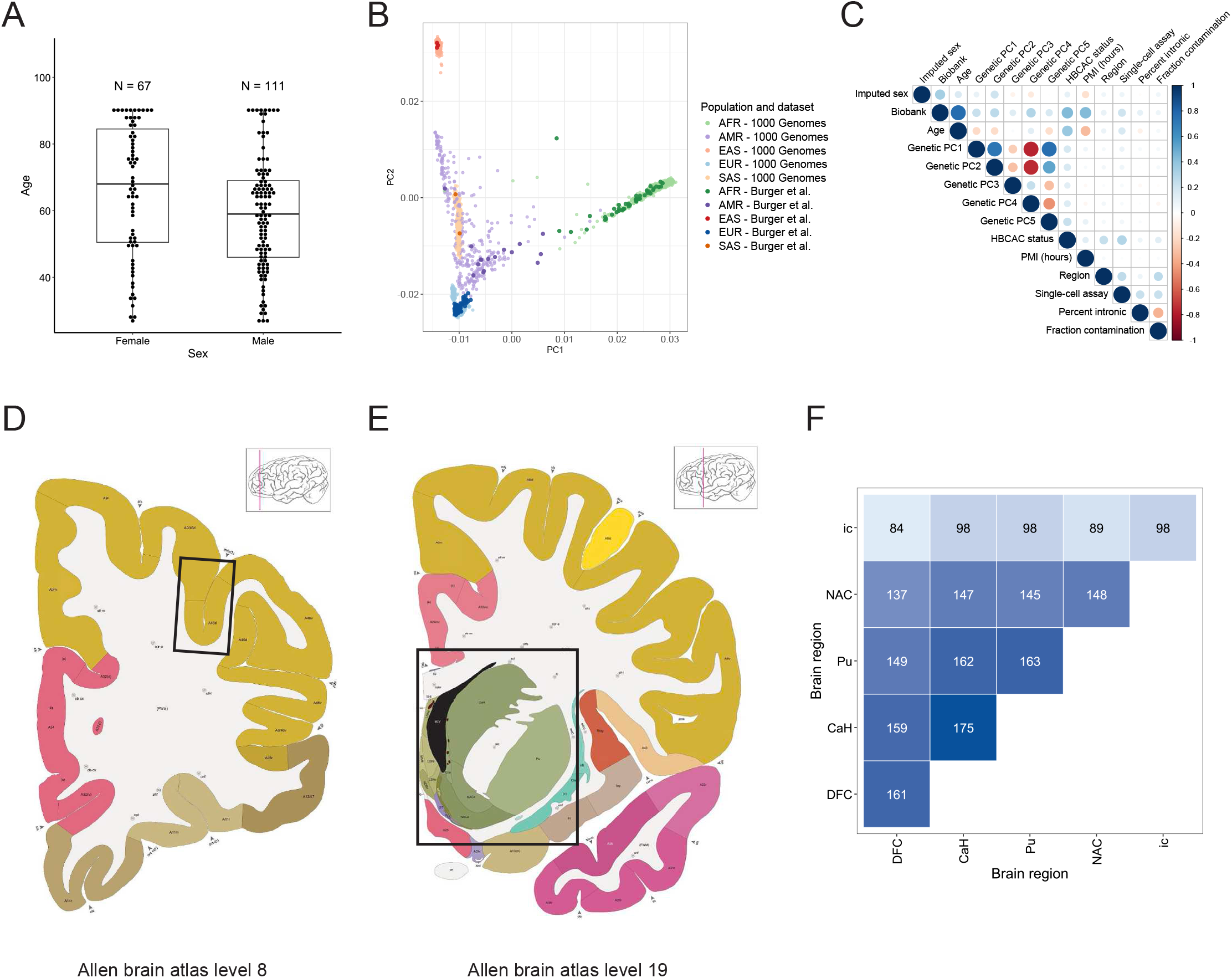
Overview of donor sampling, metadata structure, and tissue dissection reference for regions of interest (ROIs). **A.** Counts and age distribution of female and male donors. **B.** Donor PCA embedding with 1000 Genomes reference. Ancestry PCs and inferred superpopulation assignment for each donor is provided in **Table S2**. **C.** Heatmap summarizing pairwise associations among demographic, genetic, technical, and sequencing-quality variables. Numeric-numeric pairs show Pearson correlations, factor-factor pairs show Cramér’s V, and mixed numeric-factor pairs show the proportion of variation in the numeric variable explained by group differences. Pearson correlations range from −1 to 1, whereas the other measures range from 0 to 1. Circle size and color indicate the magnitude and direction of association. A complete description of variables is provided in **Table S3**. **D-E.** Images from the Allen Institute adult human brain atlas, modified Brodmann annotation^1^. Black box represents the requested ROIs from NBB. Atlas level for (**D**) dorsolateral prefrontal cortex (DFC) samples and (**E**) striatum complex (caudate, putamen, nucleus accumbens, and internal capsule) samples. **F.** Donor sampling across brain regions. For each pair of brain regions, the heatmap shows the number of donors for which both regions were sampled. Values along the diagonal are the total donor counts for each region, as shown in Figure 1B.

**Figure S2.**
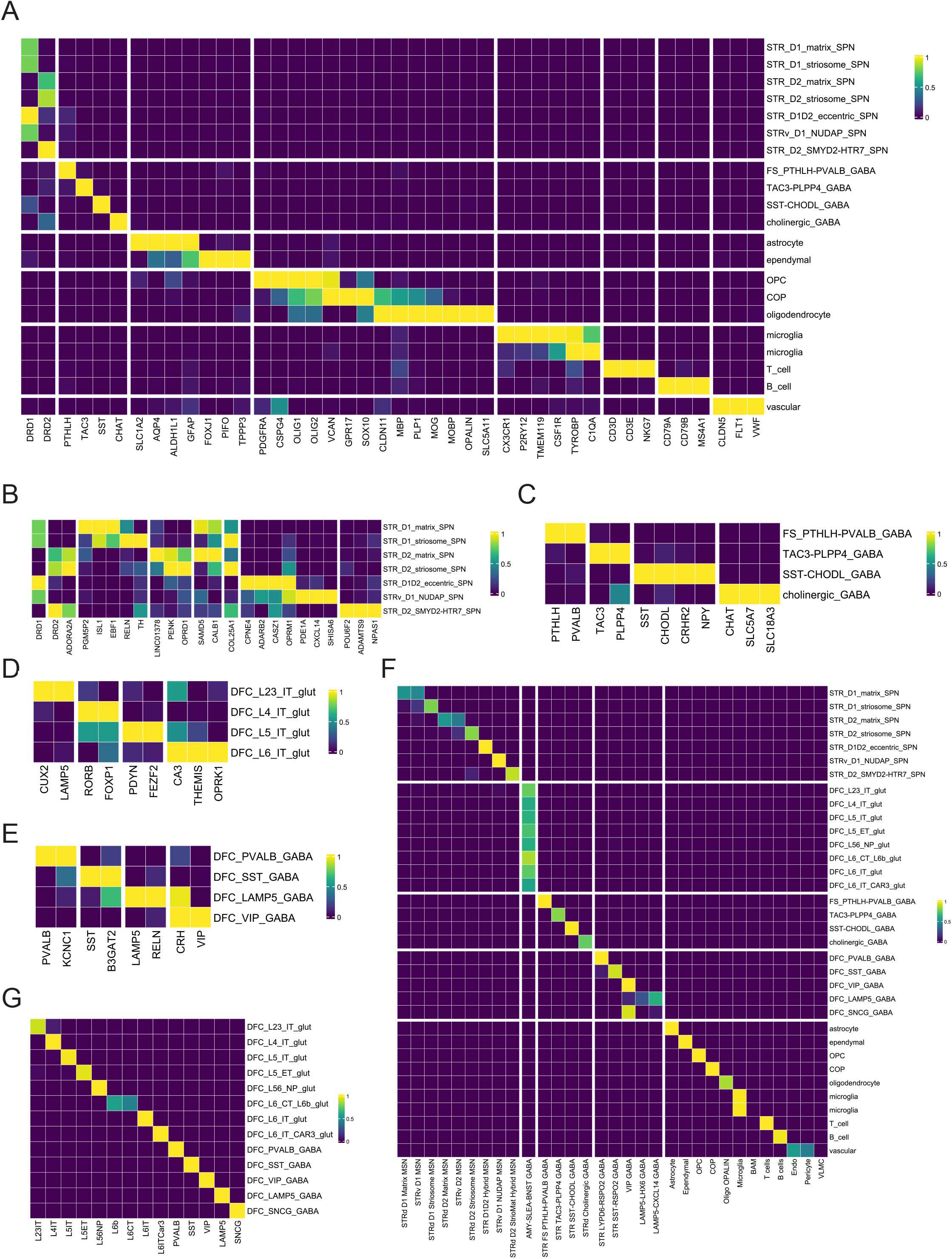
Overview of cell type annotation, marker genes, and taxonomy alignment. **A.** Cell-type specificity of gene expression measurements. Heatmap shows expression measurements for some of the genes (columns) whose expression levels distinguish major cell classes (rows) (neurons of the striatum and non-neuronal population from the striatum and DFC). Colors represent normalized expression by column (across major cell classes) such that the highest expression is 1 (bright yellow) and lowest is 0 (dark purple). **B-E.** Expression levels of selected marker genes for (**B**) SPNs, (**C**) striatal interneurons, (**D**) cortical intratelencephalic projection neurons, and (**E**) cortical interneurons. Colors represent expression levels that have been normalized by column (e.g., in **B**, across the SPN types shown) such that highest expression is 1 (bright yellow) and lowest is 0 (dark purple). **F.** Confusion matrix comparing cell identities as inferred from two analytical approaches: MapMyCells label transfer from the consensus basal ganglia taxonomy (columns), and manual unsupervised clustering annotations (rows). Values (colors) represent fractions (of the total) in each row. **G.** Confusion matrix comparing cell identities as inferred from scPred label transfer from the Allen Brain human cortical neuron taxonomy (columns), and manual unsupervised clustering annotations (rows). Values (colors) represent fractions of the total in each row.

**Figure S3.**
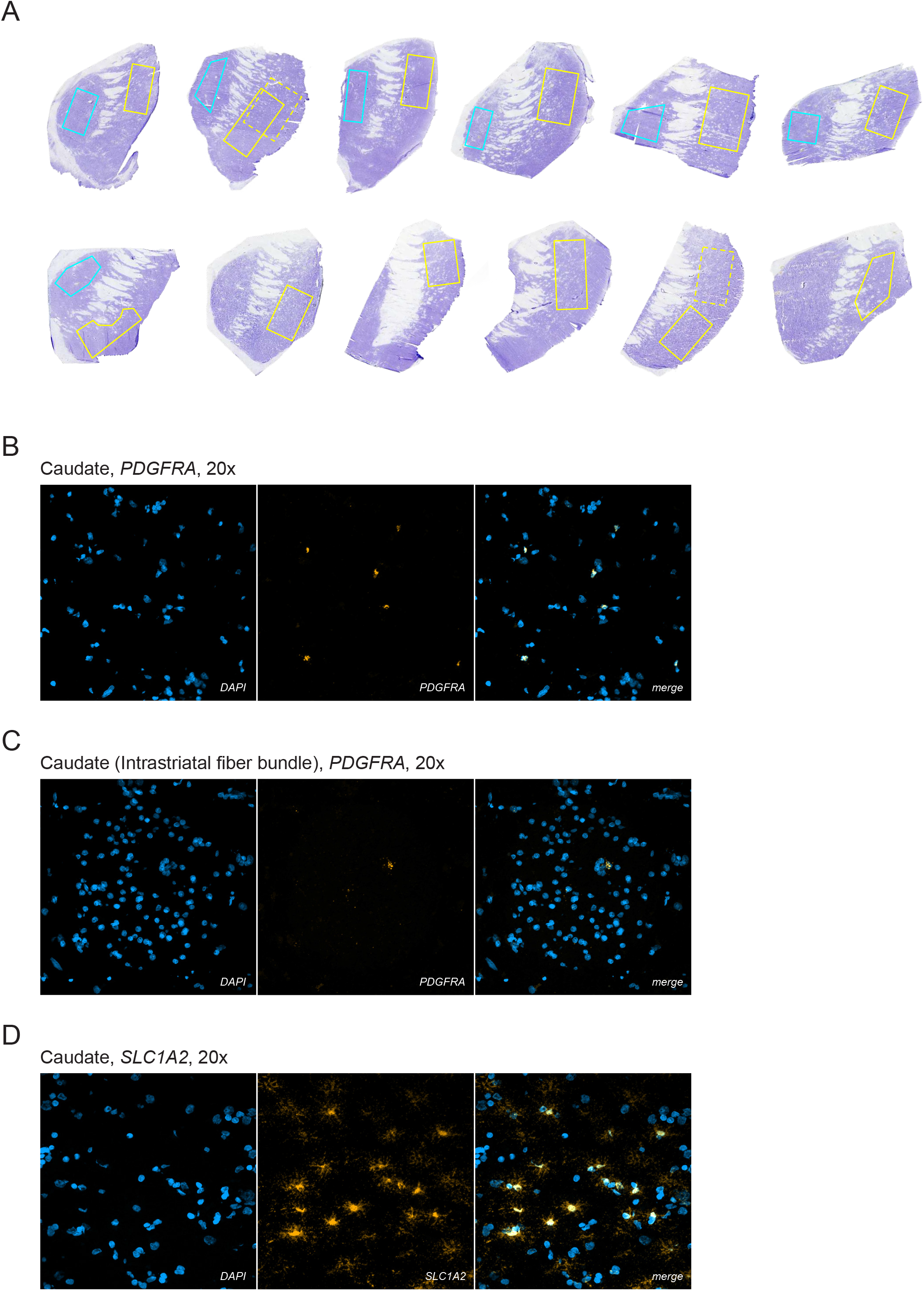
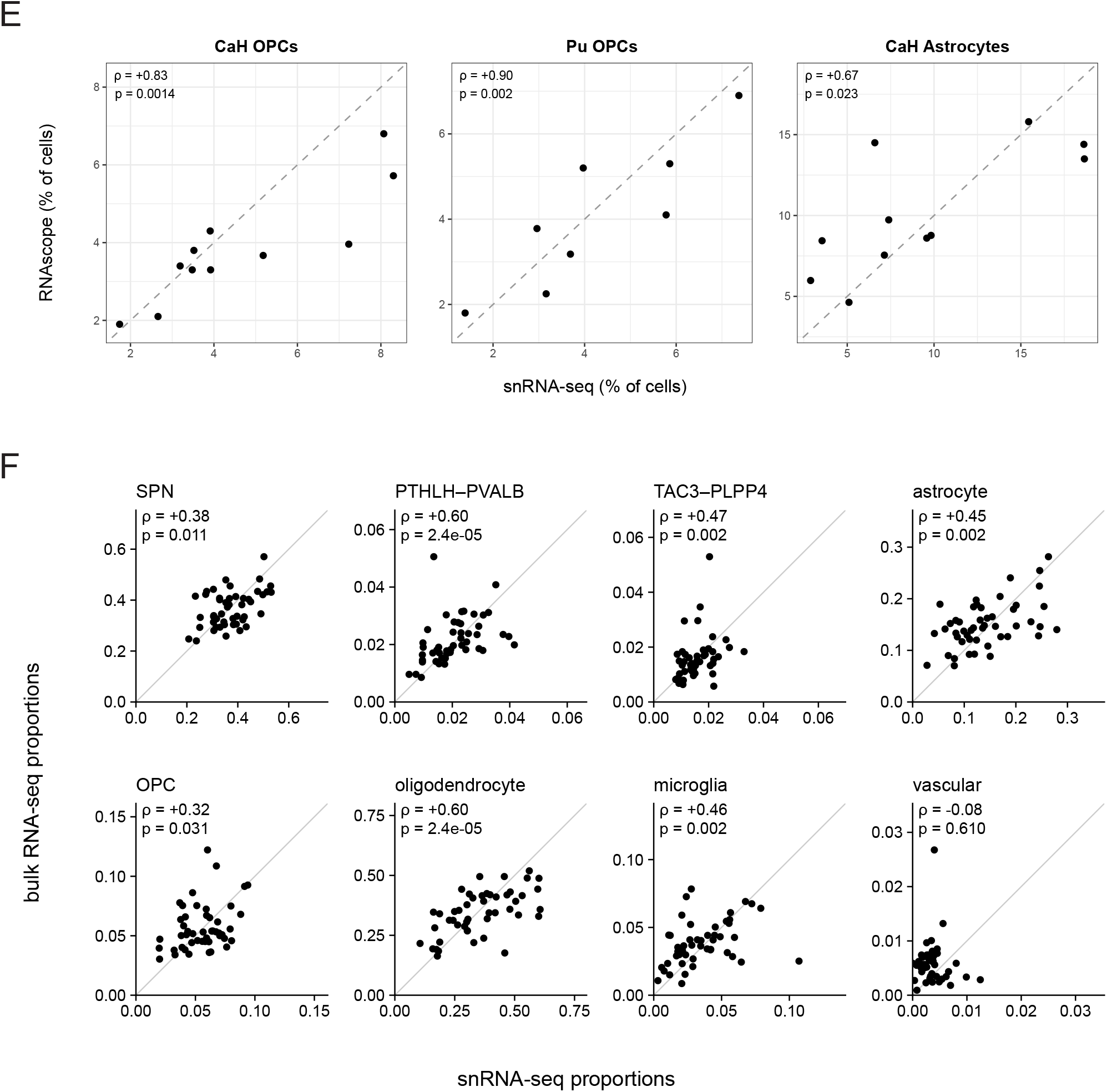
Cell type abundance analyses using RNAscope and bulk RNA-seq. **A.** Nissl-stained human striatum sections. Adjacent 12 µm sections of RNAscope slides were collected from each donor and stained with cresyl violet acetate for anatomical reference. Imaging area approximation was labeled with yellow (caudate) and blue (putamen) annotations. For donors in which *PDGFRA* and *SLC1A2* imaging was performed in different areas, solid lines indicate *PDGFRA* ROIs and dotted lines indicate *SLC1A2* ROIs. **B.** Representative RNAscope image of *PDGFRA* OPC labeling in caudate. Blue, DAPI. Yellow, *PDGFRA* signal. Image taken under 20x magnification. **C.** Representative RNAscope image of *PDGFRA* OPC labeling in an intrastriatal fiber bundle in caudate. Blue, DAPI. Yellow, *PDGFRA* signal. Image taken under 20x magnification. **D.** Representative RNAscope image of *SLC1A2* astrocyte labeling in caudate. Blue, DAPI. Yellow, *SLC1A2* signal. Image taken under 20x magnification. **E.** RNAscope observed cell-type proportions (y-axis) versus snRNA-seq reported cell-type proportions. Left, *PDGFRA*-labeled OPC proportions in caudate (n = 11) and putamen (n = 8). Right, *SLC1A2*-labeled astrocyte proportions in caudate (n = 11). Per-donor OPC and astrocyte proportions ascertained by snRNA-seq and RNAscope were significantly rank-correlated in all three conditions: caudate OPCs (Spearman ρ = +0.83, p = 0.0014), putamen OPCs (ρ = +0.90, p = 0.002), and caudate astrocytes (ρ = +0.67, p = 0.023). **F.** Cell-type proportions predicted from bulk RNA-seq (y-axis) versus snRNA-seq cell-type proportions (x-axis) in CaH, across the same 48 donors (free scales). Each point represents a donor; the gray line indicates identity. Predictions were generated using a tree based machine learning deconvolution model (see **Methods**). Spearman correlation coefficients and corrected p-values are shown for each cell type (Benjamini-Hochberg (BH) procedure, n = 8 tests). Seven of eight cell types (all other than vascular cells) are significantly correlated.

**Figure S4.**
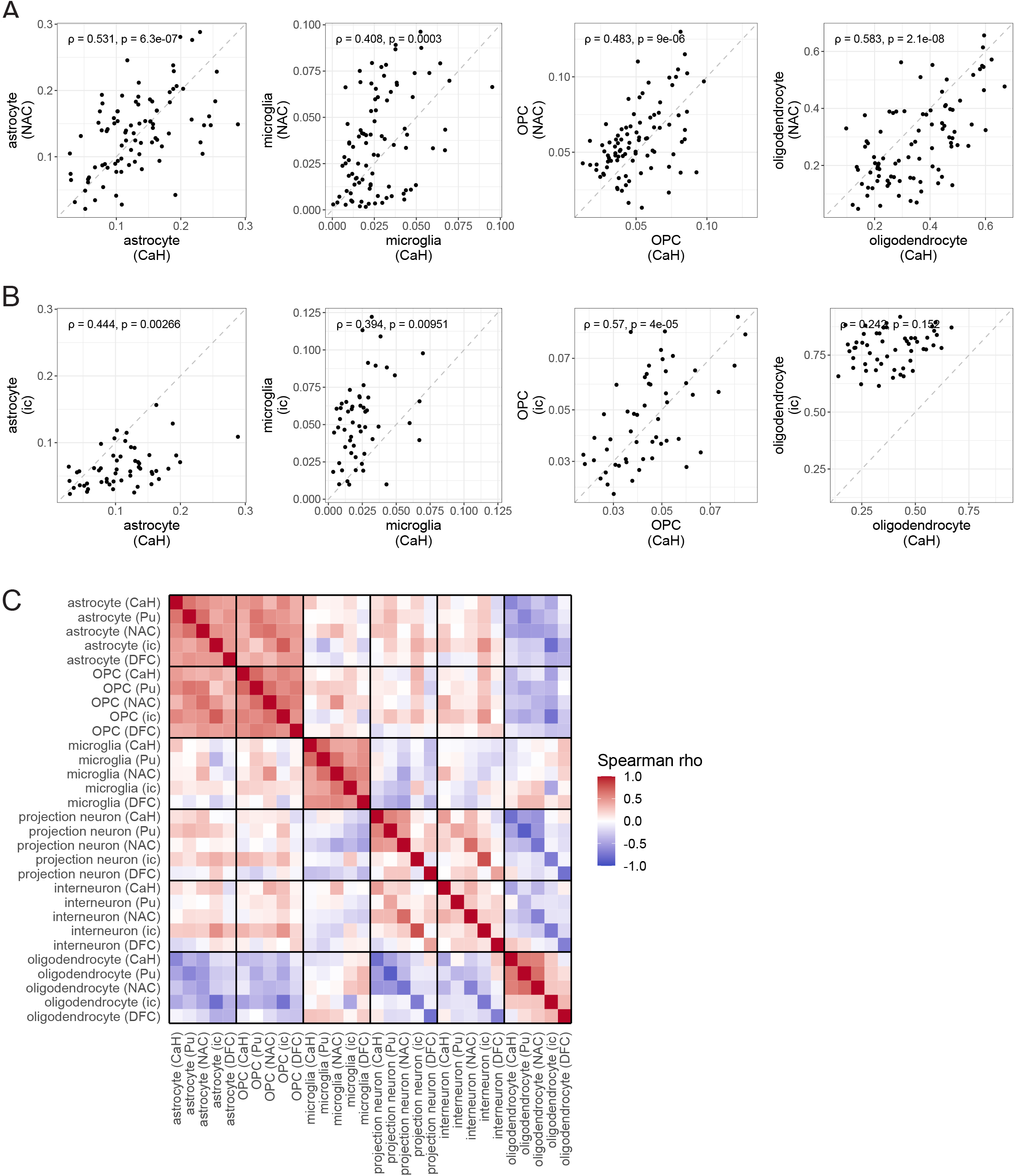
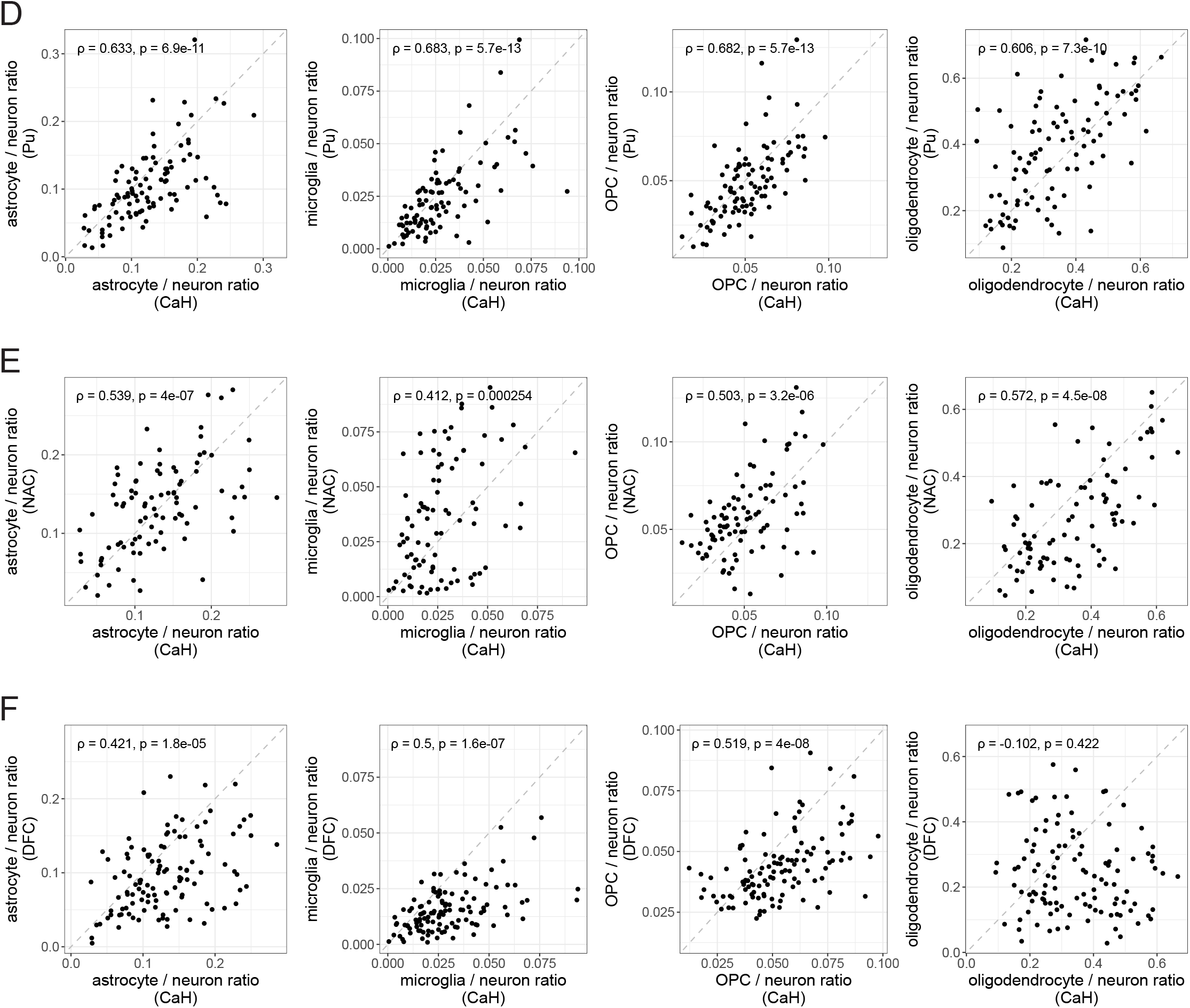
Cross-brain-region correlations in cell type abundances. Related to Figure 2**. A.** Representations of glial cell types (astrocytes, microglia, OPCs, oligodendrocytes) in the caudate (x-axis) and nucleus accumbens (y-axis) in the same 90 donors (points). Abundances are quantified as fractions of all nuclei ascertained in a donor. Spearman correlation coefficient and corrected p-values are shown for each cell type (Benjamini-Hochberg (BH) procedure, n = 498 tests). **B.** Same as **A**, for the caudate (x-axis) and internal capsule (y-axis) across 54 donors. **C.** Pairwise correlations of cell type abundance measurements – quantified as fractions of all nuclei ascertained in a sample – in all five brain regions surveyed. Spearman correlation coefficients are shown; positive correlations in red, negative in blue. Note the blocks of positive correlation (red) for specific cell types, regardless of the brain regions in which they are measured. **D.** Representations of glial cell types in the caudate (x-axis) and putamen (y-axis) across the same 98 donors. Here, abundances were quantified relative to the total number of neurons in the same sample, so that the glial cells do not influence one another’s abundance estimates. Spearman correlation coefficient and corrected p-values are shown for each cell type (Benjamini-Hochberg (BH) procedure, n = 498 tests). **E.** Same analysis as in **D**, focusing on correlation between caudate (x-axis) and nucleus accumbens (y-axis) across 90 donors. **F.** Same analysis as in **D**, comparing caudate (x-axis) and DFC (y-axis) across 114 donors. Astrocyte, microglia, and OPC abundances are significantly correlated, indicating that these patterns extend beyond the striatum.

**Figure S5.**
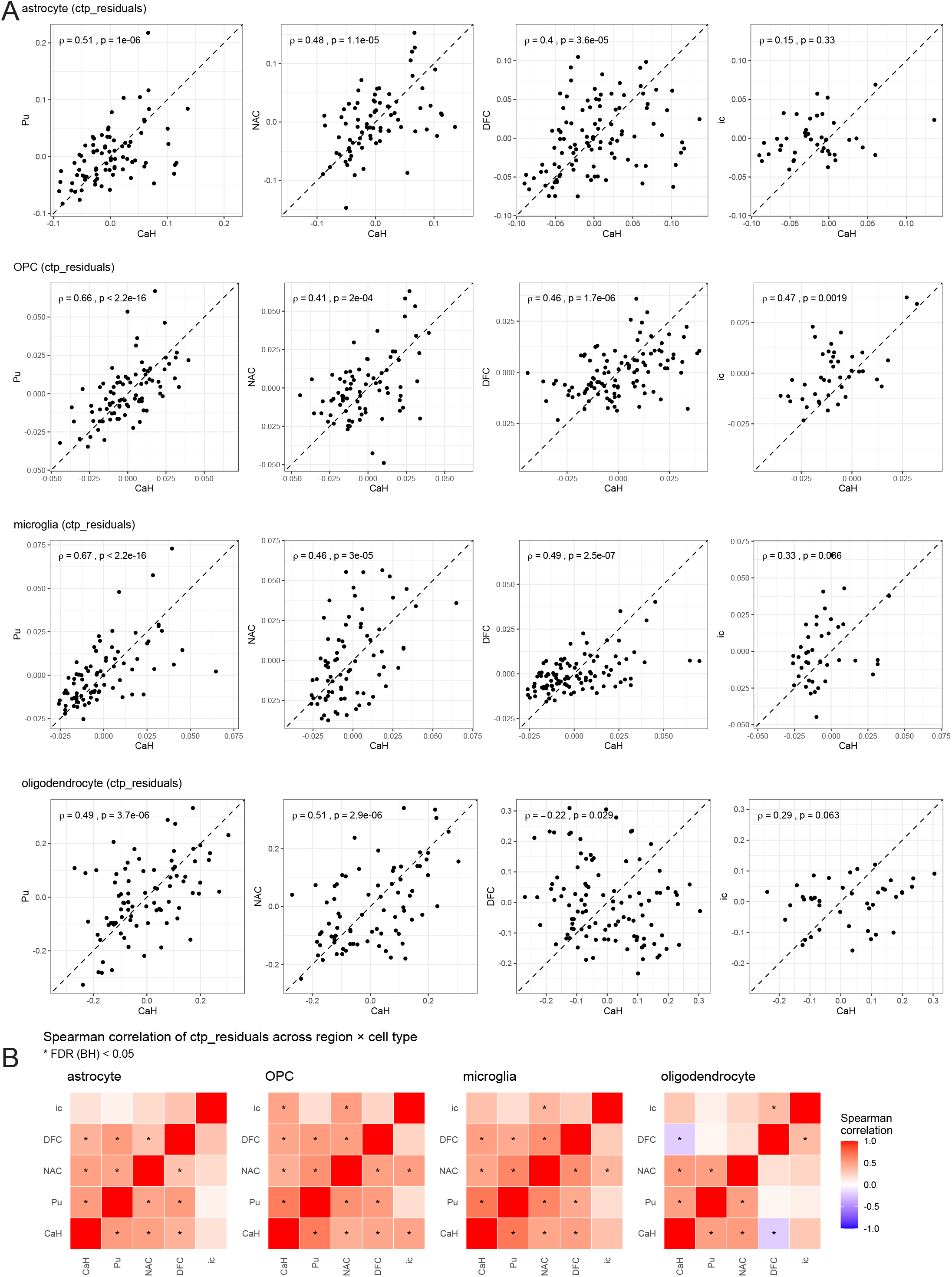
Cross-region correlations after technical metadata correction. Related to Figure 2**. A.** For each glial cell type within each brain region, cell-type proportion was regressed against PMI, RQS, and normalized library size (see **Methods**). Residuals were calculated to represent the component of variation not explained by a linear relationship with these features. Residual astrocyte, OPC and microglial abundances remained positively and significantly correlated across gray matter regions. **B.** Cross-region correlation heatmaps of CTP residuals for each glial cell type reveal glial correlation structure persists after technical metadata correction. Asterisks indicate statistically significant correlations at an FDR of 0.05 using BH correction (100 tests).

**Figure S6.**
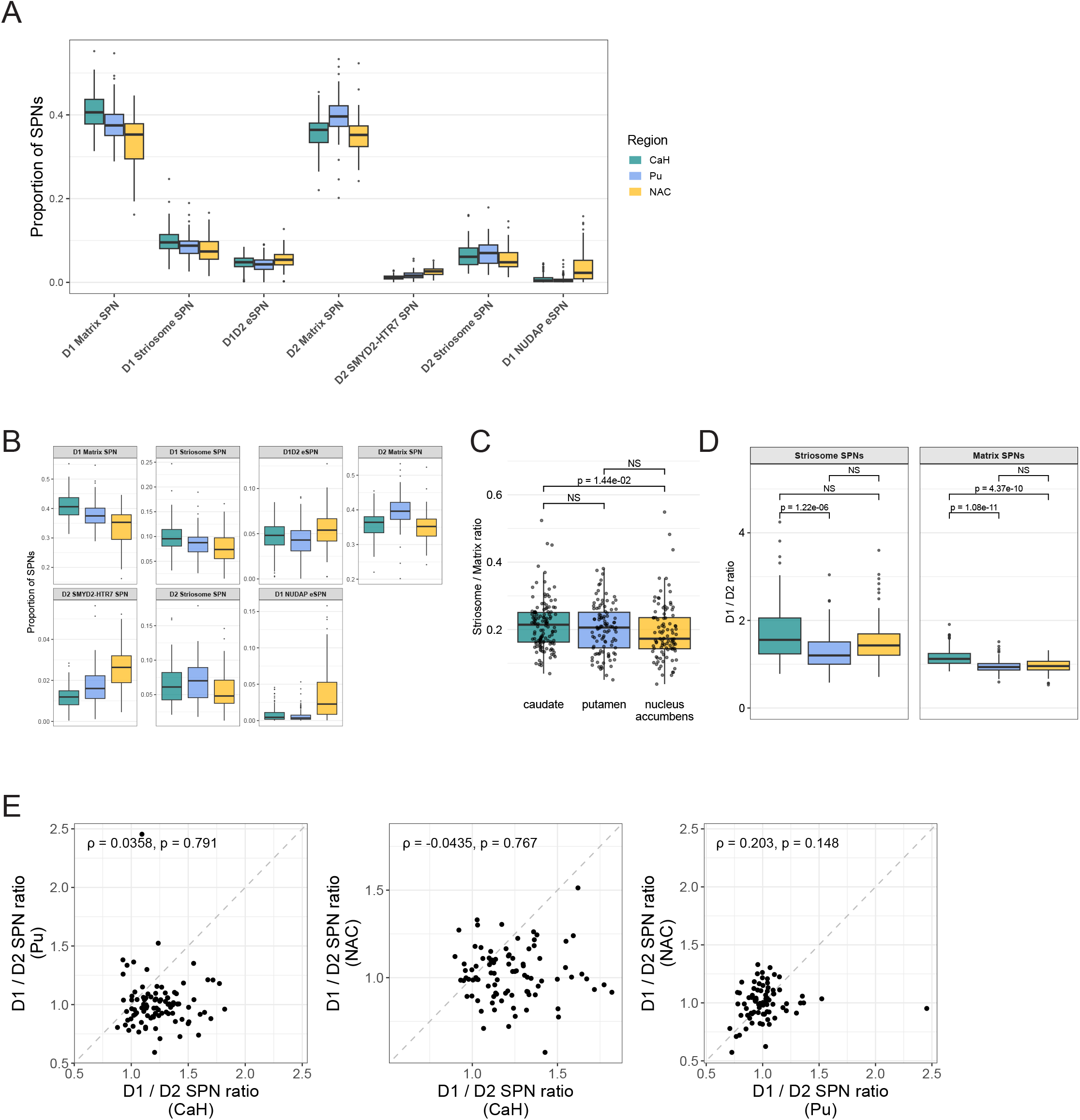
Regional composition of striatal neuron subtypes. Related to Figure 2**. A.** Proportions of SPN subtypes in CaH, Pu, and NAC. D1 and D2 matrix SPNs make up the majority of SPNs in the striatum. Boxes indicate the interquartile range across donors and whiskers extend to the most extreme points within 1.5x the interquartile range. **B.** Distributions of SPN proportions in the striatum (free scales). **C.** The striosome/matrix SPN ratio does not vary dramatically between regions, although there may be a slight striosome depletion in NAC, as evaluated by paired Wilcoxon signed-rank test. **D.** Comparison of D1/D2 SPN ratios across CaH, Pu, and NAC, stratified by striosome/ matrix compartments and evaluated by paired Wilcoxon signed-rank test. **E.** Comparisons (and lack of significant correlations) between D1/D2 SPN ratios comparing different pairs of striatal regions. Spearman correlation coefficient and corrected p-values are shown (Benjamini-Hochberg (BH) procedure, n = 498 tests). Left: caudate (x-axis) versus putamen (y-axis), n = 98 donors. Middle: caudate (x-axis) versus nucleus accumbens (y-axis), n = 90 donors. Right: putamen (x-axis) versus nucleus accumbens (y-axis), n = 78 donors.

**Figure S7.**
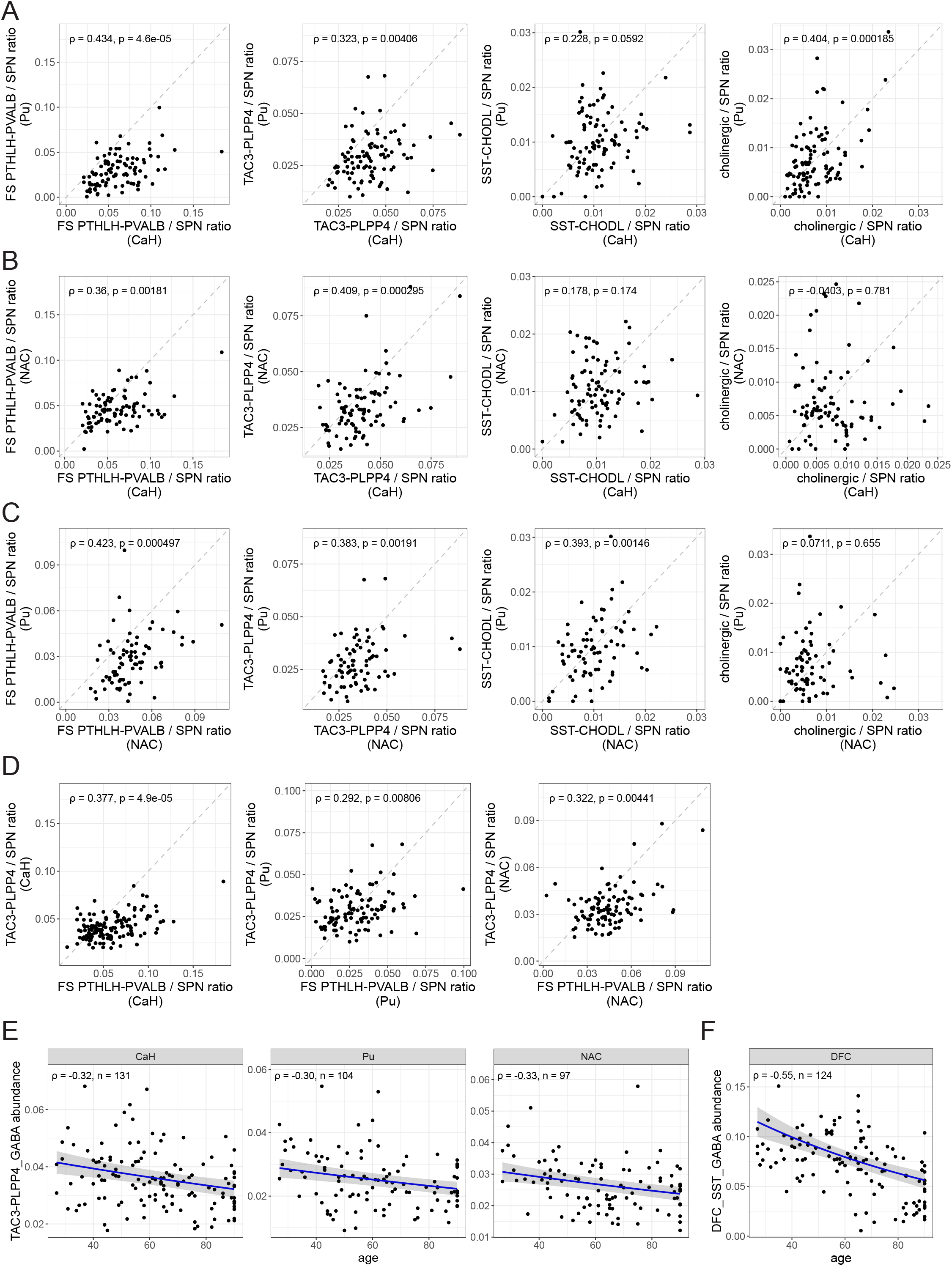
Relationships among the abundances of various types of striatal interneurons. Related to Figure 2. In all plots, abundances have been quantified relative to the number of SPNs (in the same tissue sample) to reduce dependence of these measurements on one another and to minimize the effects of variance in glial populations. **A.** Abundances of various types of interneurons in the caudate (x-axis) versus putamen (y-axis) across 98 donors (points). Spearman correlation coefficient and corrected p-values are shown (Benjamini-Hochberg (BH) procedure, n = 498 tests). PTHLH-PVALB, TAC3-PLPP4, and cholinergic interneuron subtype abundances were significantly correlated between caudate and putamen. **B.** Same analysis as in **A**, comparing caudate (x-axis) and nucleus accumbens (y-axis) across 90 donors. PTHLH-PVALB and TAC3-PLPP4 interneuron subtype abundances are significantly correlated. **C.** Same analysis as in **A**, comparing nucleus accumbens (x-axis) and putamen (y-axis) across 78 donors. PTHLH-PVALB, TAC3-PLPP4, and SST-CHODL interneuron subtype abundances were significantly correlated. **D.** Within-sample comparison of PTHLH- PVALB and TAC3-PLPP4 abundances. Left: caudate (n = 131 donors). Middle: putamen (n = 104 donors). Right: nucleus accumbens (n = 97 donors). Spearman correlation coefficient and corrected p-values are shown (BH-procedure, n = 498 tests) **E.** Attrition of TAC3-PLPP4 interneurons with advancing age, in all striatal gray matter regions (OR = 0.959 per decade, corresponding to a loss of approximately 4% per decade; BH-adjusted p-value = 0.003; n = 54 tests; **Table S5**). Abundance is expressed relative to SPNs (TAC3 / TAC3 + SPN). Each point represents a donor, and blue lines indicate beta-binomial fits with 95% confidence intervals (gray ribbons). **F.** Attrition of cortical SST interneurons with advancing age in DFC (OR = 0.884,corresponding to a loss of approximately 11% per decade; BH-adjusted p-value = 5.36e-11; n = 54 tests; **Table S5**). Abundance is expressed relative to glutamatergic projection neurons (SST / SST + glutamatergic neurons). Each point represents a donor, and blue lines indicate beta-binomial fits with 95% confidence intervals (gray ribbons).

**Figure S8.**
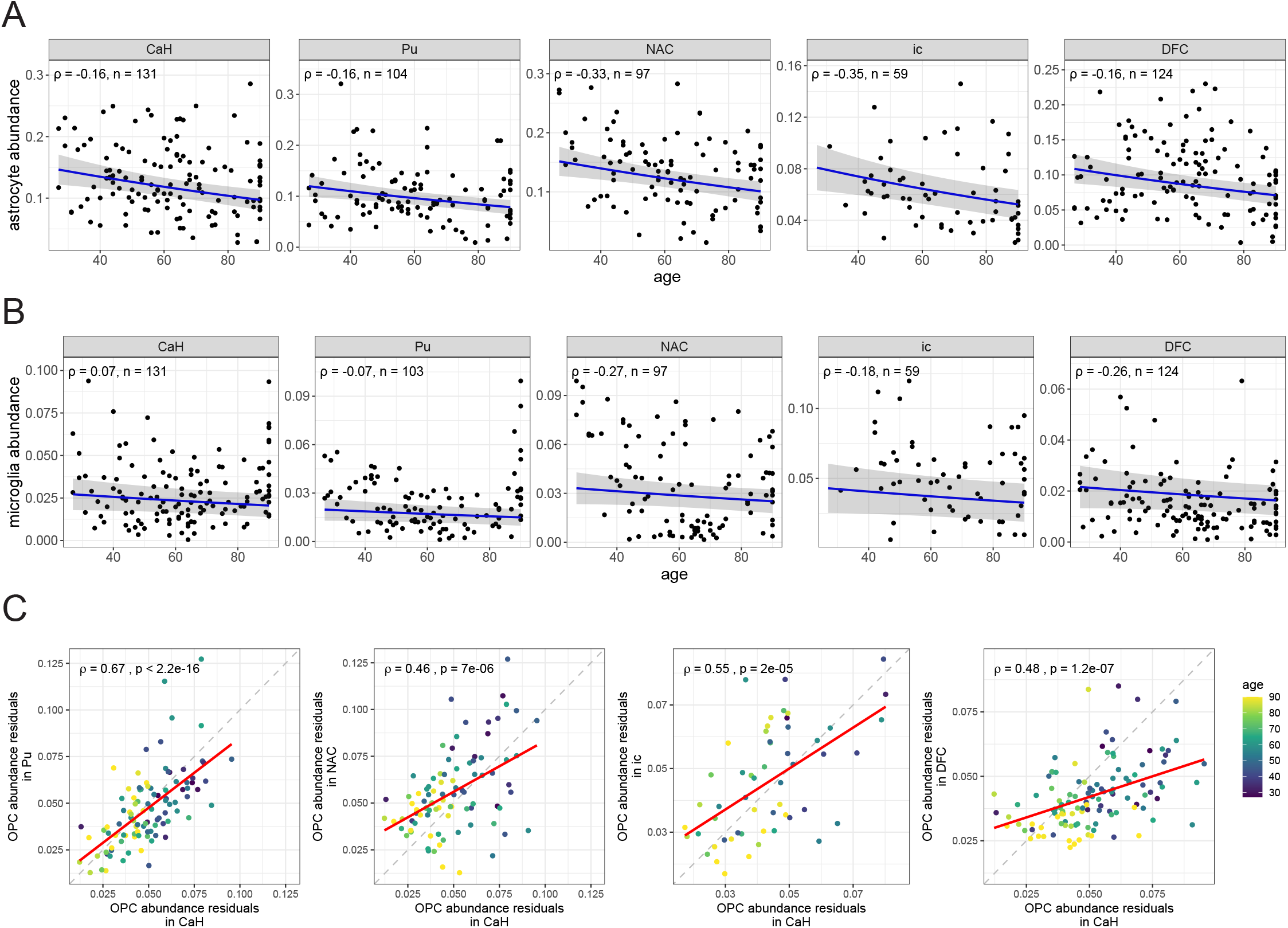
Additional analyses of glial cell abundances in relationship to age. Related to **Figure 3. A.** Abundance of astrocytes, as a fraction of all nuclei sampled. Each point represents a donor, and blue lines indicate beta-binomial fits with 95% confidence intervals (gray ribbons). While nominally-significant negative Spearman correlations were observed in specific regions, the result of global modeling via beta-binomial regression failed to meet our analysis wide BH-adjusted p-value threshold of < 0.01 (OR = 0.929; 95% CI = 0.893 - 0.969; BH-adjusted p-value = 2.70e-2; n = 54 tests; **Table S5**). **B.** Same as **A**, but for microglia abundance. Beta-binomial regression found no significant association between microglial abundance and age (OR = 0.977; 95% CI = 0.927 - 1.031; BH-adjusted p-value = 1; n = 54 tests; **Table S5**). **C.** Comparison of OPC fraction residuals (observed minus age-predicted values from covariate-adjusted beta-binomial models) for the caudate compared to the other four brain regions analyzed. Each point represents a donor, colored by age; red lines indicate linear regression fits. The positive correlations in these residuals across all pairs of brain regions indicate that additional donor-level factors (beyond chronological age) influence OPC abundance in the striatum complex and DFC in a similar manner.

**Figure S9.**
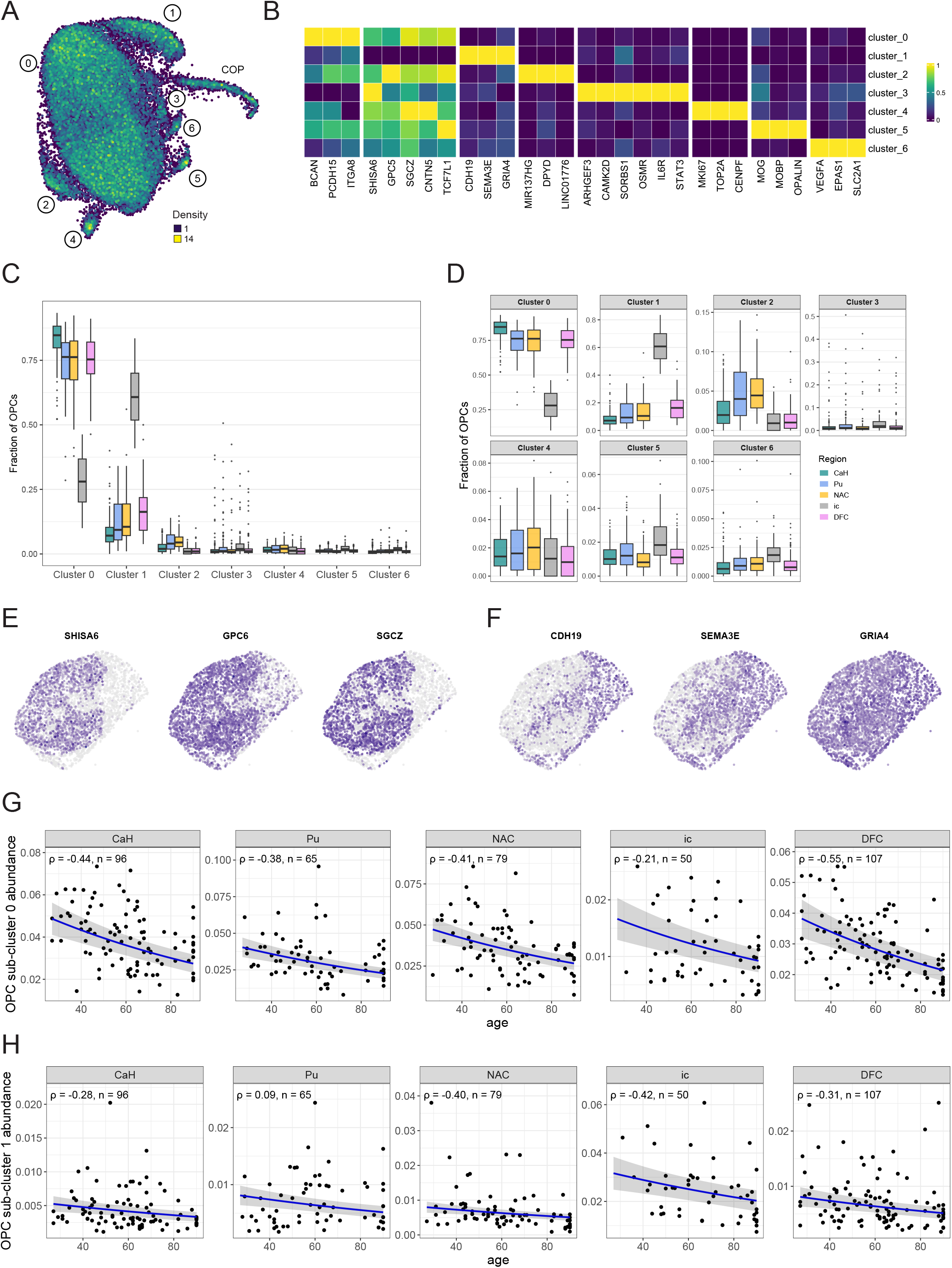
OPC diversity and loss with age. Related to. **Figure 3. A.** Unsupervised clustering assignments and UMAP representation of OPCs sampled from the caudate. Colors on the UMAP plot represent the density of nuclei in that region of the plot. **B.** Expression levels of specific marker genes (columns) for OPC sub-clusters (rows; note, committed oligodendrocyte progenitors (COPs), excluded). Colors represent normalized expression by column (across major cell classes) such that the highest expression is 1 (bright yellow) and lowest is 0 (dark purple). **C.** Distribution of OPC sub-cluster compositions across regions. Gray matter regions exhibit much lower numbers of cluster 1 OPCs and higher numbers of cluster 0 OPCs relative to the internal capsule. **D.** Distribution of OPC sub-cluster compositions across regions (free scales). **E-F.** Slide-tags spatial transcriptomics analysis of the striatum from a representative donor. Expression of genes whose expression defines OPC cluster 0 (e.g. *SHISA6*, *GPC6*, and *SGCZ*) highlights gray matter compartments. Color of the points represents log-normalized transcript counts for each gene (light gray lower expression, dark purple higher expression). Expression of genes whose expression defines OPC cluster 1 (*CDH19*, *SEMA3E*) highlights white matter compartments. *GRIA4* is expressed in all OPC sub-clusters, but most strongly in cluster 1, as seen in **B**. **G.** Decline in abundance of OPC cluster 0 with advancing age, across the brain regions sampled. Abundance is expressed as a fraction of all nuclei sampled. Each point represents a donor, and blue lines indicate beta-binomial fits with 95% confidence intervals (gray ribbons). Spearman correlation coefficients and number of donors shown. Modeling with a beta-binomial regression confirmed a significant decline with age (OR = 0.906, 95% CI = 0.882 - 0.931, BH-adjusted p-value = 1.36e-10; n = 54 tests; **Table S5**). **H.** Same as **G**, but for OPC cluster 1. Modeling with a beta-binomial regression confirmed a significant decline with age (OR = 0.927, 95% CI = 0.891 - 0.964, BH-adjusted p-value = 8.91e-3; n = 54 tests; **Table S5**).

**Figure S10.**
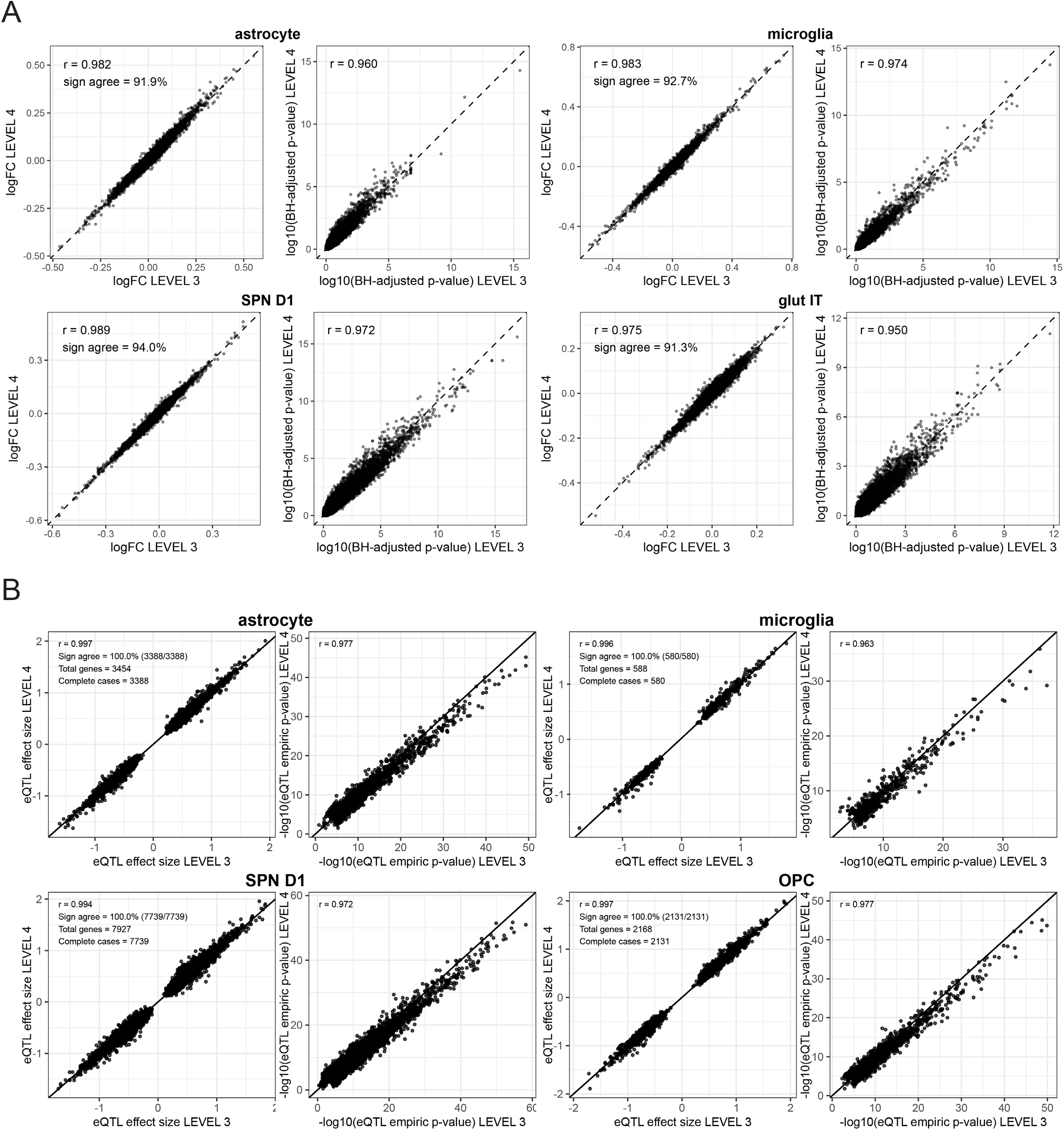
Effects of donor exclusion criteria on recognition and inference of age-associated changes. Related to **STAR Methods**. **A.** Comparison, in multiple cell types, of gene-level log fold-change estimates and statistical significance (-log10 FDR- adjusted p-values) for age-associated genes as donor exclusion criteria are made more stringent. Stringency Level 3 represents snRNA-seq-data-driven exclusions; Level 4 additionally excludes donors based on metadata criteria from clinical records. Effect-size estimates remain highly concordant across filtering levels, and statistical significance is largely preserved or improved under snRNA-seq-data-driven filtering. In contrast, adding metadata-driven exclusions reduces statistical power to detect effects at analysis-wide significance, while leaving effect-size estimates largely unchanged. **B.** Comparison of log fold-change estimates and statistical significance (-log10 empirical p-values) for expression quantitative trait loci (eQTL) discovery. As in **A**, Level 3 represents snRNA-seq-data-driven exclusions; Level 4 additionally excludes donors based on metadata criteria (findings in medical histories). Effect-size estimates remained highly concordant across filtering levels, and statistical significance is largely preserved or improved under snRNA-seq-data-driven filtering. In contrast, adding metadata-driven exclusions reduced statistical significance while effect-size estimates were largely unchanged. Annotated are the correlation of effect sizes, fraction of the time the direction of the eQTL effect sizes agree (sign test), total number of genes compared (union of eGenes), and total number of genes discovered in both data sets (intersect of eGenes).

**Figure S11.**
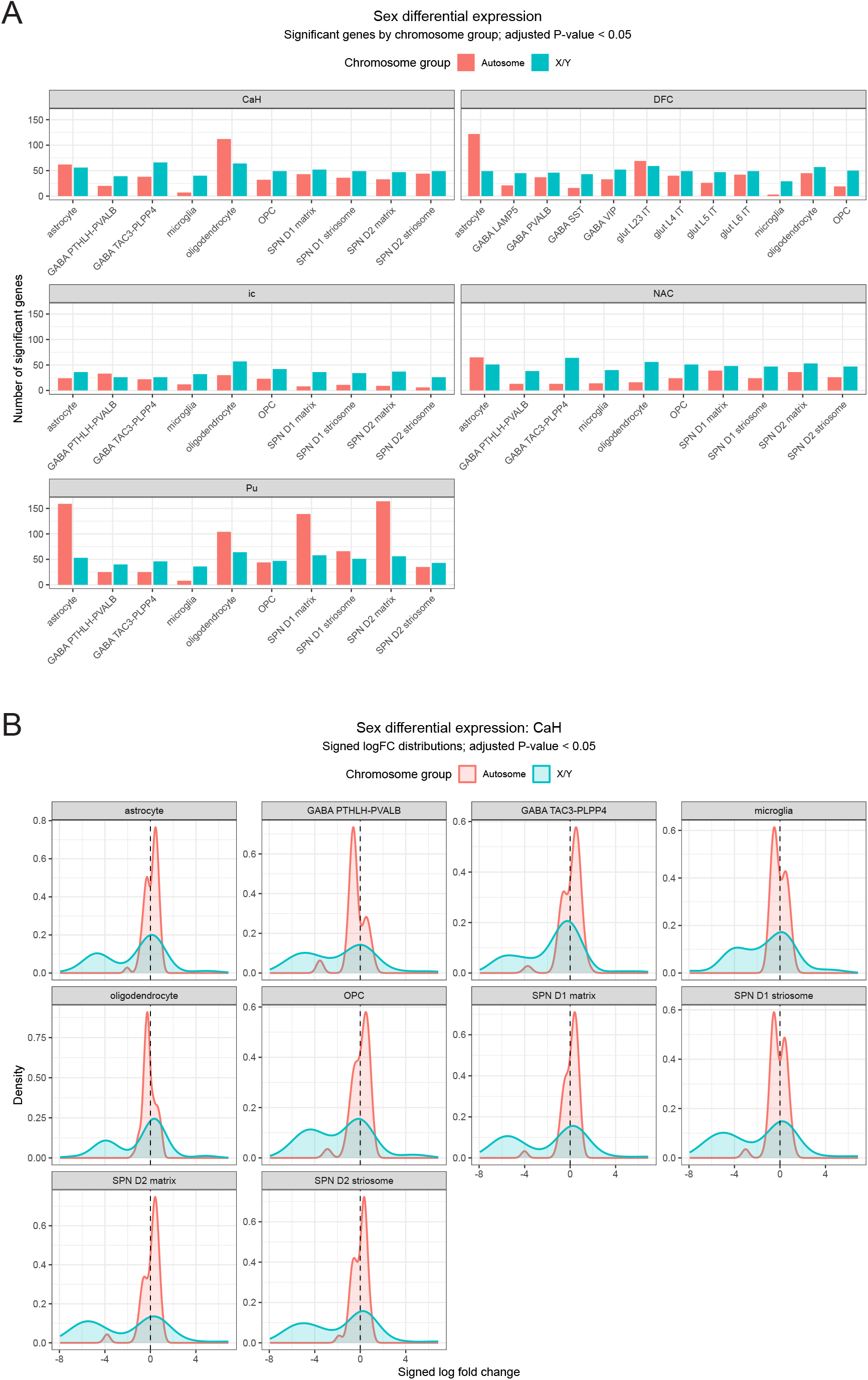
Sex-associated gene expression. Related to Figure 4. **A.** Number of significant genes by chromosome group (autosome or X/Y) in cell types and brain regions sampled. **B.** Effect size distributions for X/Y and autosomal genes in CaH.

**Figure S12.**
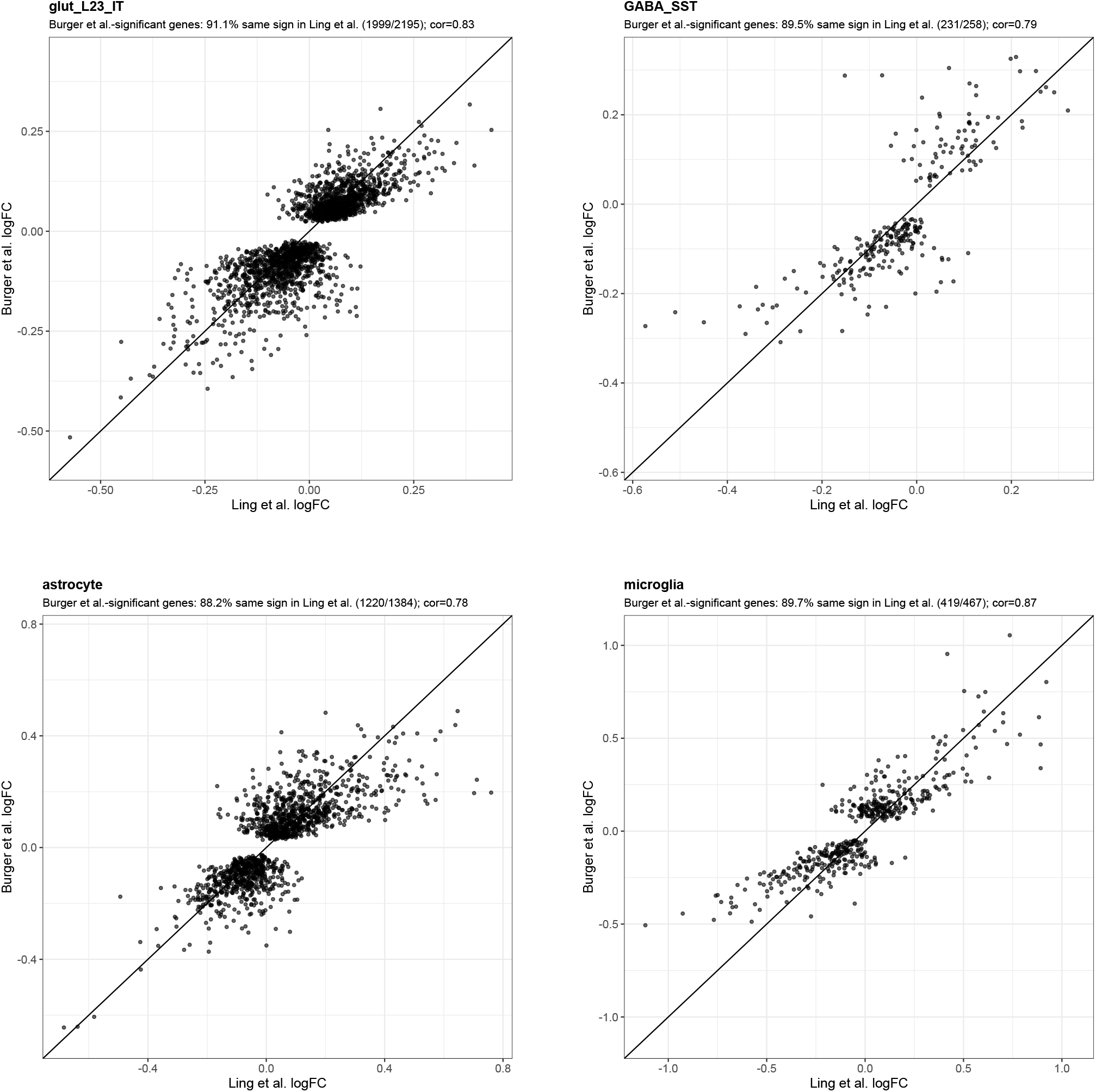
External validation of age-associated differential expression results. Related to Figure 4. Scatter plots comparing age-associated gene-expression effect sizes (log fold-change per decade) between Ling *et al.*^2^ (2024; x-axes) and this study (y-axes), by cell type. Sign-test and effect-size correlation results for additional cell types are provided in **Table S6**.

**Figure S13.**
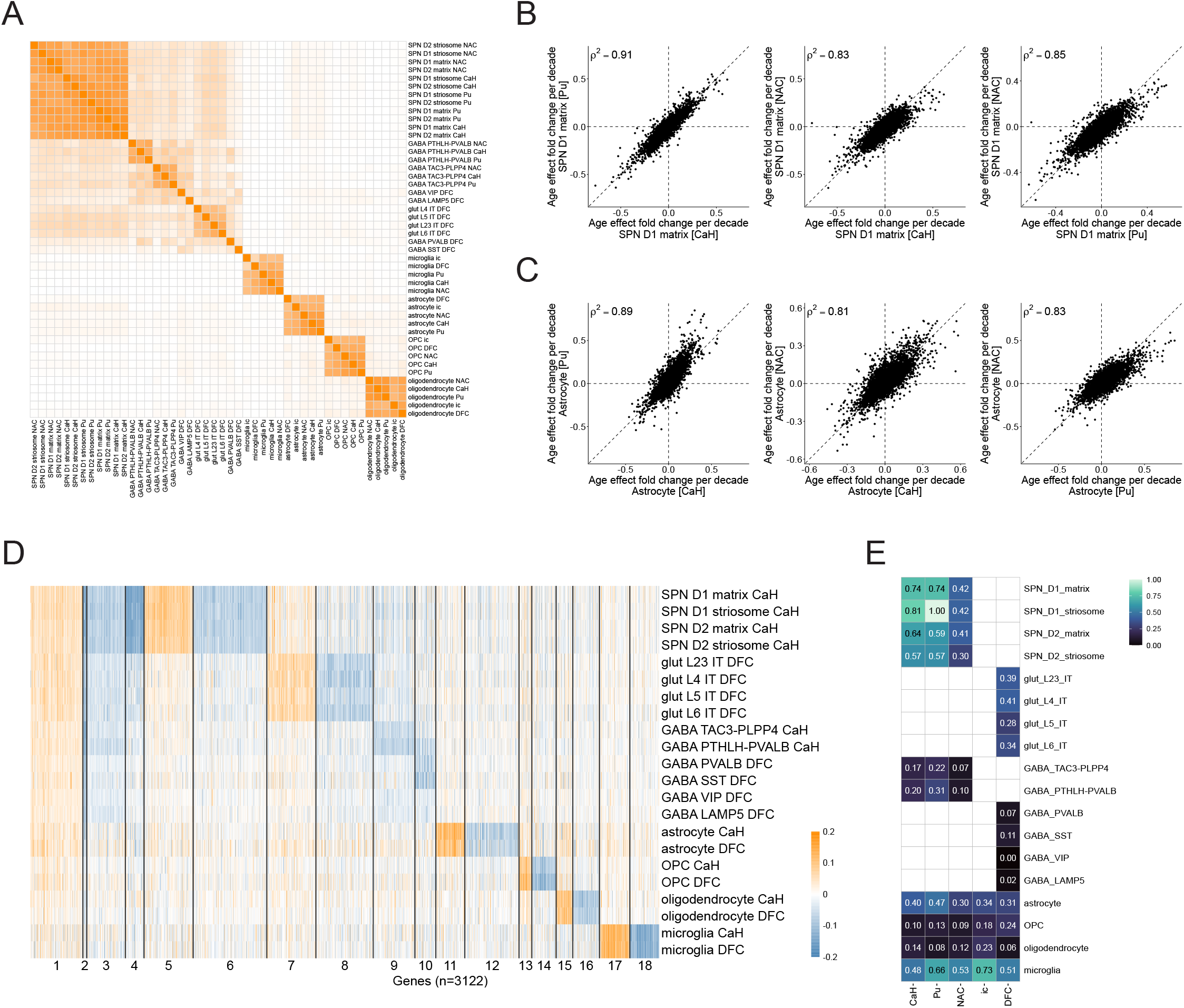
Extended analyses of aging effects. Related to Figure 4. Same analyses as in Figure 4, but extended to additional cell populations. **A.** Heatmap shows correlation of age-associated gene-expression changes for each pair of cell types and brain regions. Colors show Spearman’s ρ_2_ for correlations of gene- level log2-fold-change per decade of age. **B.** Comparison of D1 SPNs (matrix subtype) age-associated gene-expression changes between CaH and Pu, CaH and NAC, and Pu and NAC. **C.** Comparison of astrocyte age-associated gene- expression changes between CaH and Pu, CaH and NAC, and Pu and NAC. **D.** Clustering of genes by their patterns of age-associated expression changes in the various cell types and brain regions. Columns show genes (n = 2478) grouped by k-means clustering (same clustering plotted in Figure 4H) of their age-associated expression changes (log2-fold- change per decade of age) across the various cell types. **E.** Heatmap of TRADE estimated age-associated transcriptome- wide impact (TWI) by cell type (rows) and brain regions (columns). Values were linearly rescaled to fall between 0 and 1 (represented as colors and numeric values). Cell types not represented in a given region are shown in white.

**Figure S14.**
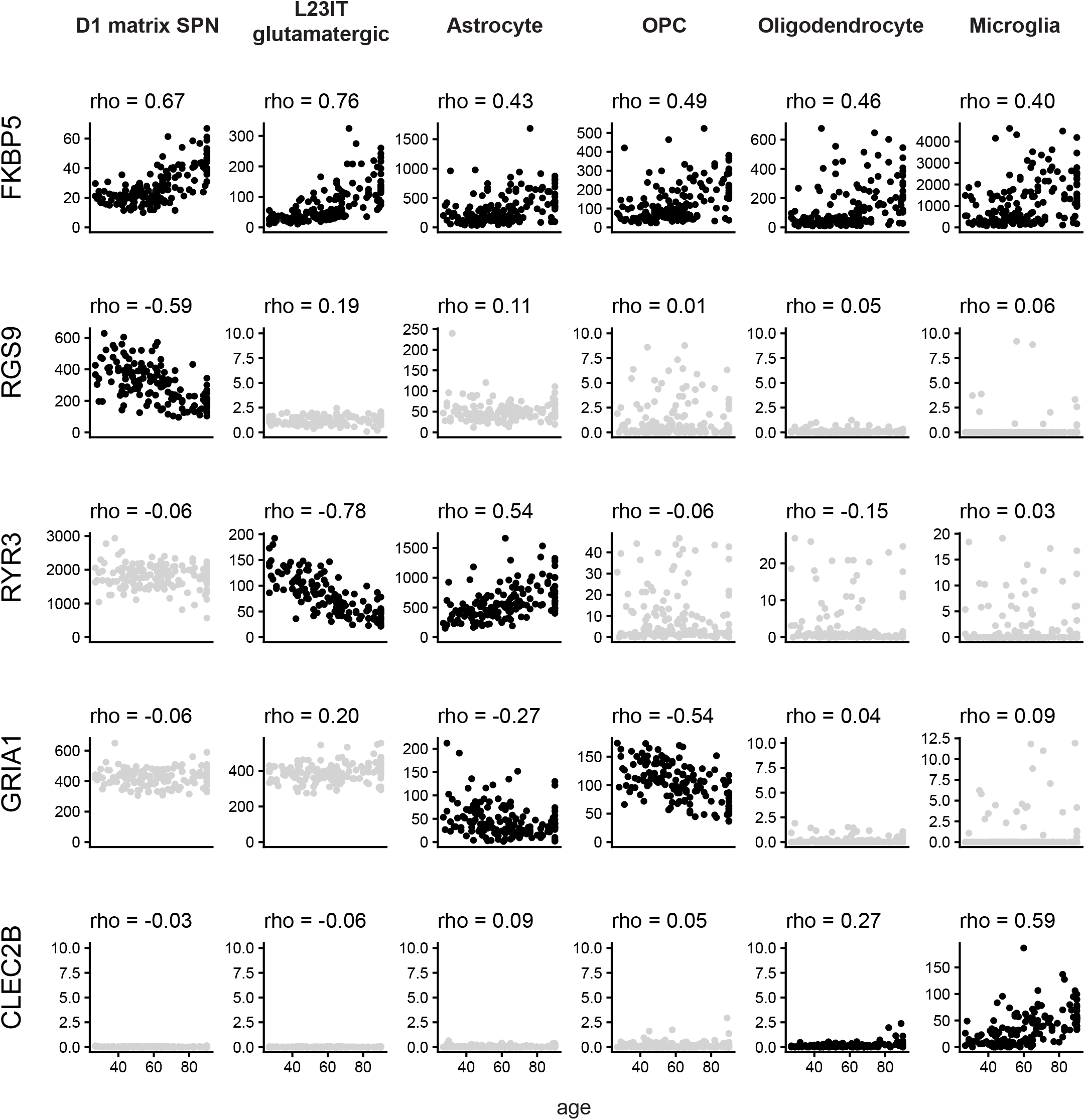
Age-associated effects on expression of specific genes. Related to Figure 4. Scatter plots show expression of specific genes in relationship to donor age (x-axis of each plot); expression levels have been normalized (transcripts per million; y-axis of each plot). Spearman’s ρ, a non-parametric statistic assessing correlation between donor age and expression level (for the selected gene, cell type) is shown above each plot. For non-significant correlations (nominal p-value > 0.05), points are shown in gray.

**Figure S15.**
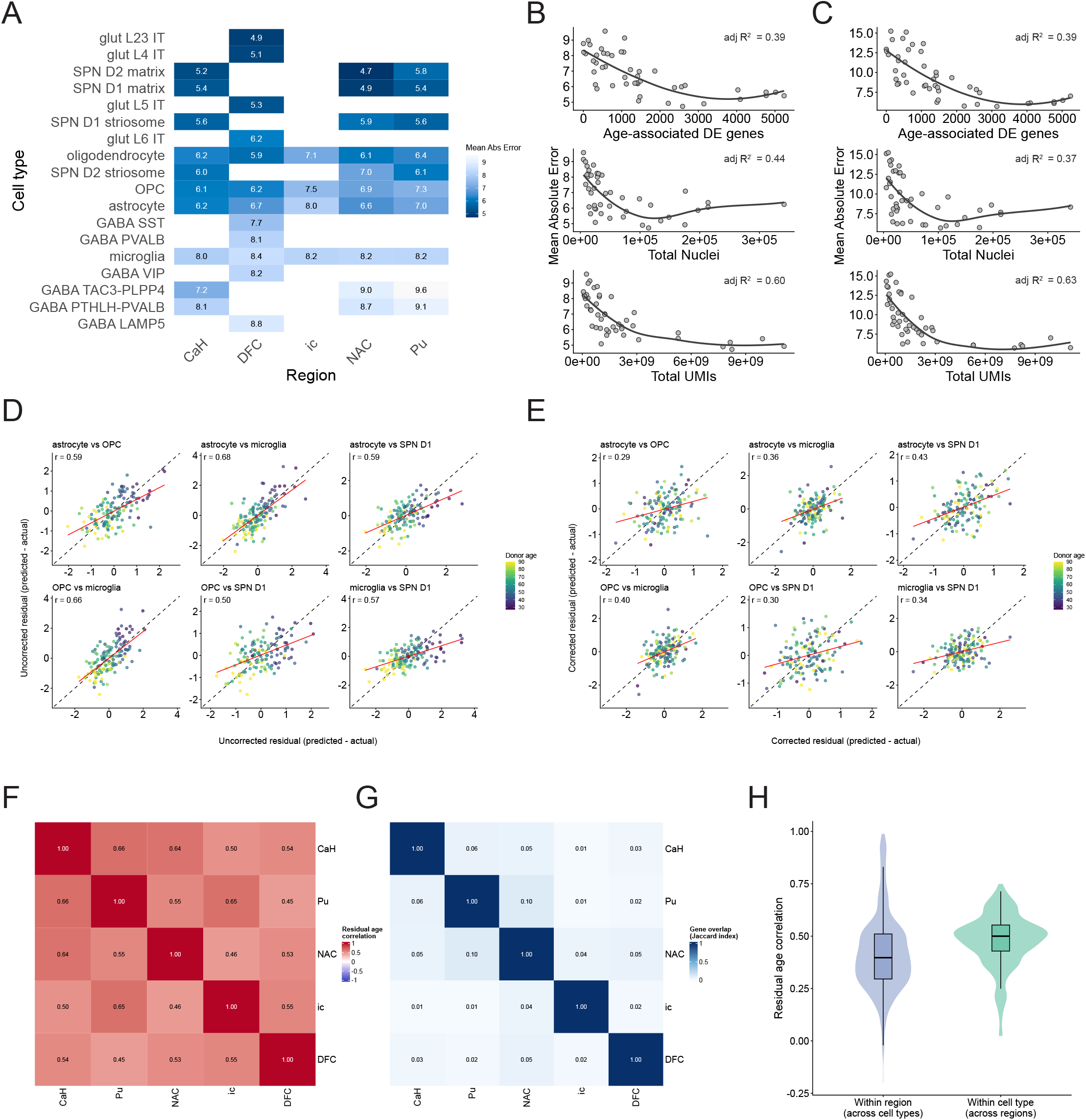
Age-prediction accuracy, bias correction, and residual correlations. Related to Figure 5**. A.** Heatmap showing mean absolute error for each cell type-region age-prediction model. Cell-type/region combinations for which age-prediction models were built are shown in shade of blue, with the shade reporting the model error. **B.** Mean absolute error plotted against the number of age-associated differentially expressed genes, total nuclei, and total UMIs per model. Adjusted R^2^ values are calculated separately for each predictor. A multivariable model including all three predictors explains variation in mean absolute error across models (adjusted R^2^ ≈ 0.736), indicating that prediction accuracy reflects both power to detect age-associated genes and cell-type-specific differences in the magnitude of age-related transcriptional effects. **C.** The analysis in **B** was repeated using only the youngest 20% of donors. Mean absolute error is higher than in the full cohort, reflecting regression-to-the-mean effects, but shows similar dependence on the predictors. **D.** Pairwise comparisons of uncorrected age residuals (predicted minus chronological age) between representative caudate cell types. Residuals were found to be positively correlated across cell types. Regression-to-the-mean effects can introduce age-dependent structure in these residuals, thereby inflating correlation across cell types. **E.** The same pairwise comparisons as **D** after GAM-based correction of predicted versus chronological age. Correlations are reduced relative to the uncorrected residuals, indicating removal of shared age-dependent bias. A non-zero correlation signal remains, consistent with shared inter-individual variation in biological aging beyond regression-to-the-mean effects. **F.** Positive correlation of cell-type-specific RNA-expression “clocks”, beyond the shared effects of chronological age. The heatmap shows pairwise correlations (for each pair of caudate cell types) of the GAM-corrected age residuals. **G.** Minimal overlap of the genes used to predict age in most cell types. The heatmap shows Jaccard indices quantifying overlap among genes used in each cell-type-specific model. **H.** Pairwise correlations of corrected residual age were computed across cell types within each region and across regions within each cell type. On average, correlations were higher across regions for a given cell type than across cell types within a region.

**Figure S16.**
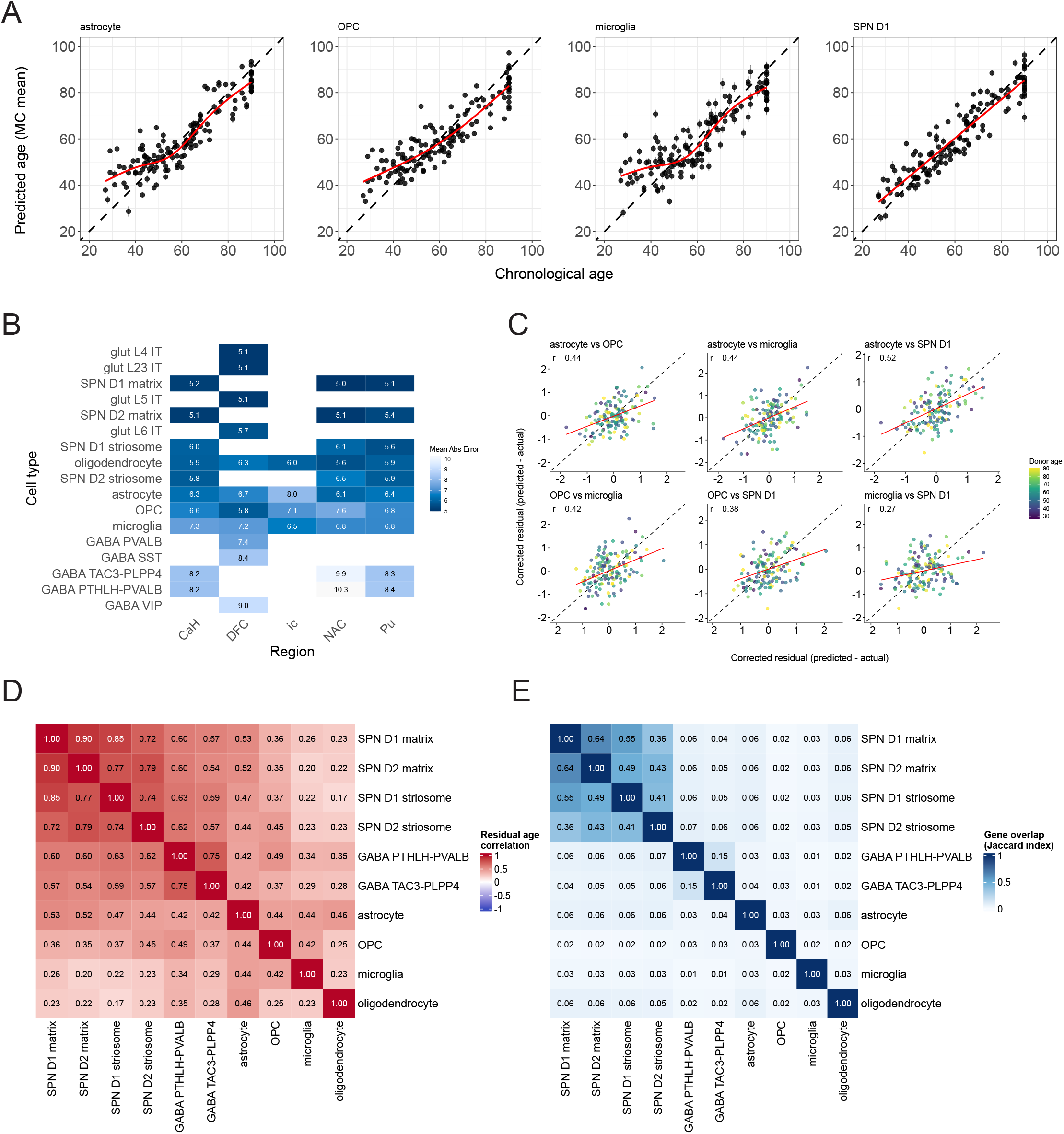
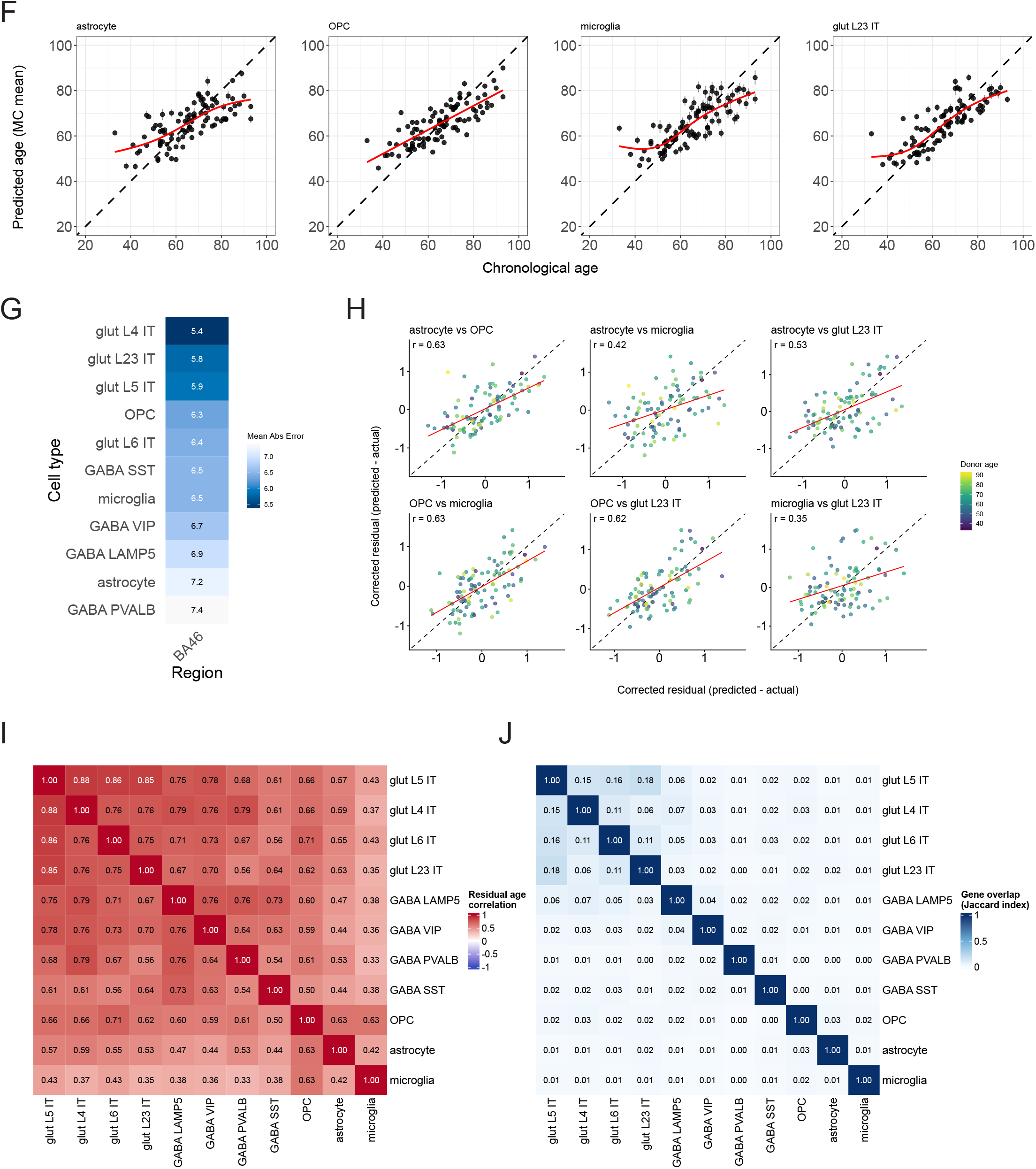
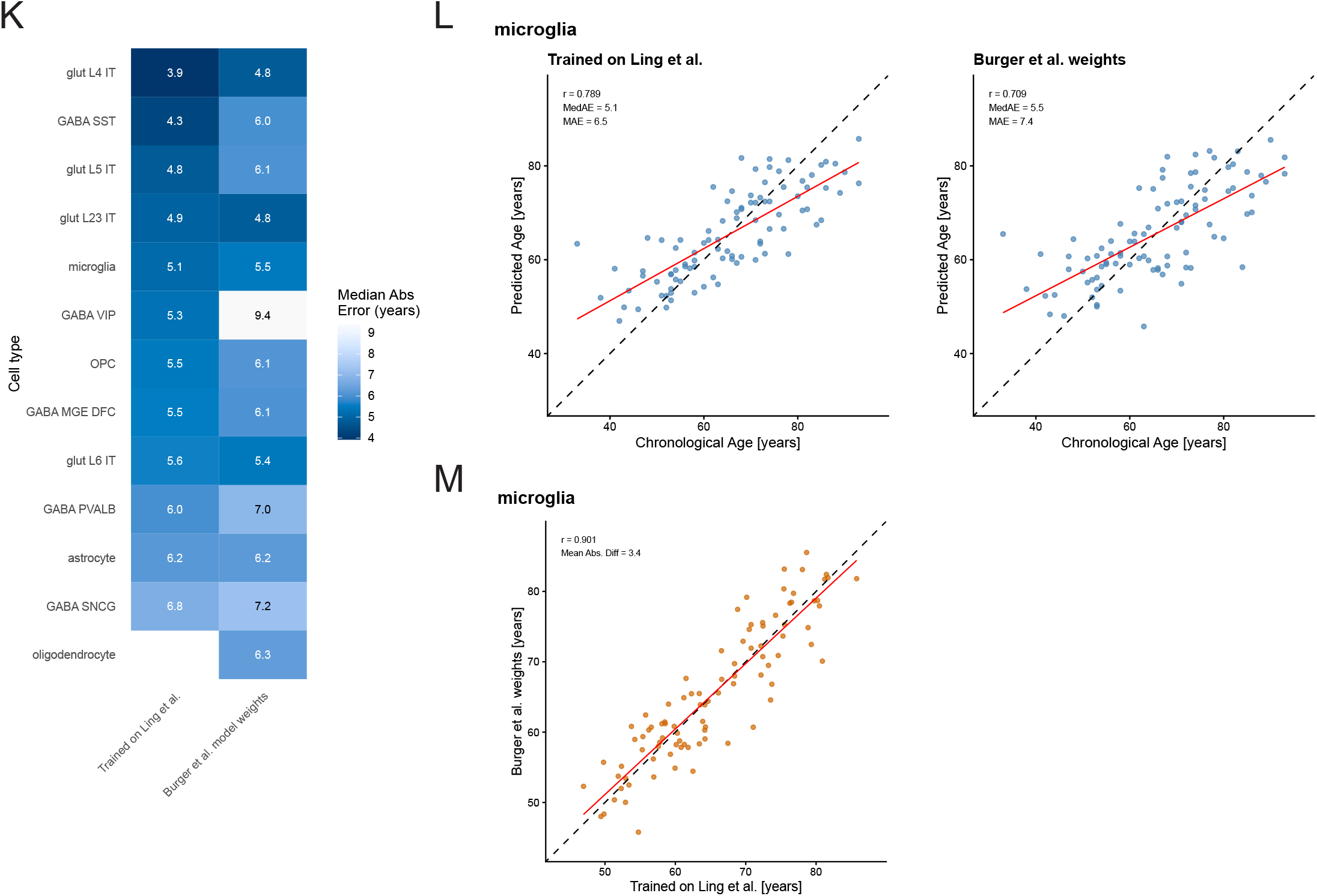
Age prediction using age-associated genes. Related to Figure 5**. A-E.** Repeating age-prediction analyses using only genes previously associated with age (Figure 4; see **Methods**). **A.** RNA-expression-based age prediction in four representative caudate cell types. The model was trained using a subset of genes previously associated with age in the corresponding region and cell type. This reproduces the main figure analysis (Figure 5B). **B.** Heatmap showing mean absolute error for each cell type-region age-prediction model. This reproduces the supplemental figure analysis (**Figure S15A**). **C.** Pairwise comparisons of GAM-corrected age residuals (predicted minus chronological age) between representative caudate cell types. Residuals remained positively correlated across cell types, reproducing the analysis in **Figure S15E**. **D.** Positive correlation of cell-type-specific RNA-expression age-prediction, beyond the shared effects of chronological age. This reproduces the main figure analysis (Figure 5C). **E.** Minimal overlap of the genes used to predict age in most cell types. The heatmap shows Jaccard indices quantifying overlap among age-associated genes used in each cell-type-specific model. This reproduces the main figure analysis (Figure 5D). **F-J.** Age-prediction analyses were repeated in the independent Ling *et al.* dataset, using age-associated DE genes identified in our primary cohort as the candidate feature set. **F.** Predicted versus chronological age for four representative BA46 cell types. **G.** Mean absolute error for each BA46 cell-type model. **H.** Pairwise comparisons of GAM-corrected age-prediction residuals (predicted minus chronological age) among representative cell types. **I.** Correlation of corrected residuals across cell types. **J.** Jaccard overlap of genes selected by the elastic-net models across cell types. **K-M.** Age prediction analysis using Burger *et al.* model weights to predict Ling *et al.* donor age. **K.** Median absolute prediction error in years for models trained directly in Ling *et al.* and for models transferred from the primary cohort after correction of the model intercept to the mean age of the Ling *et al.* cohort. **L.** Predicted versus chronological age for microglia as a representative example, comparing the independently trained Ling *et al.* model with the intercept-corrected transferred models. **M.** Agreement of donor-level microglia age predictions between the independently trained Ling *et al.* model and the model transferred from the primary cohort.

**Figure S17.**
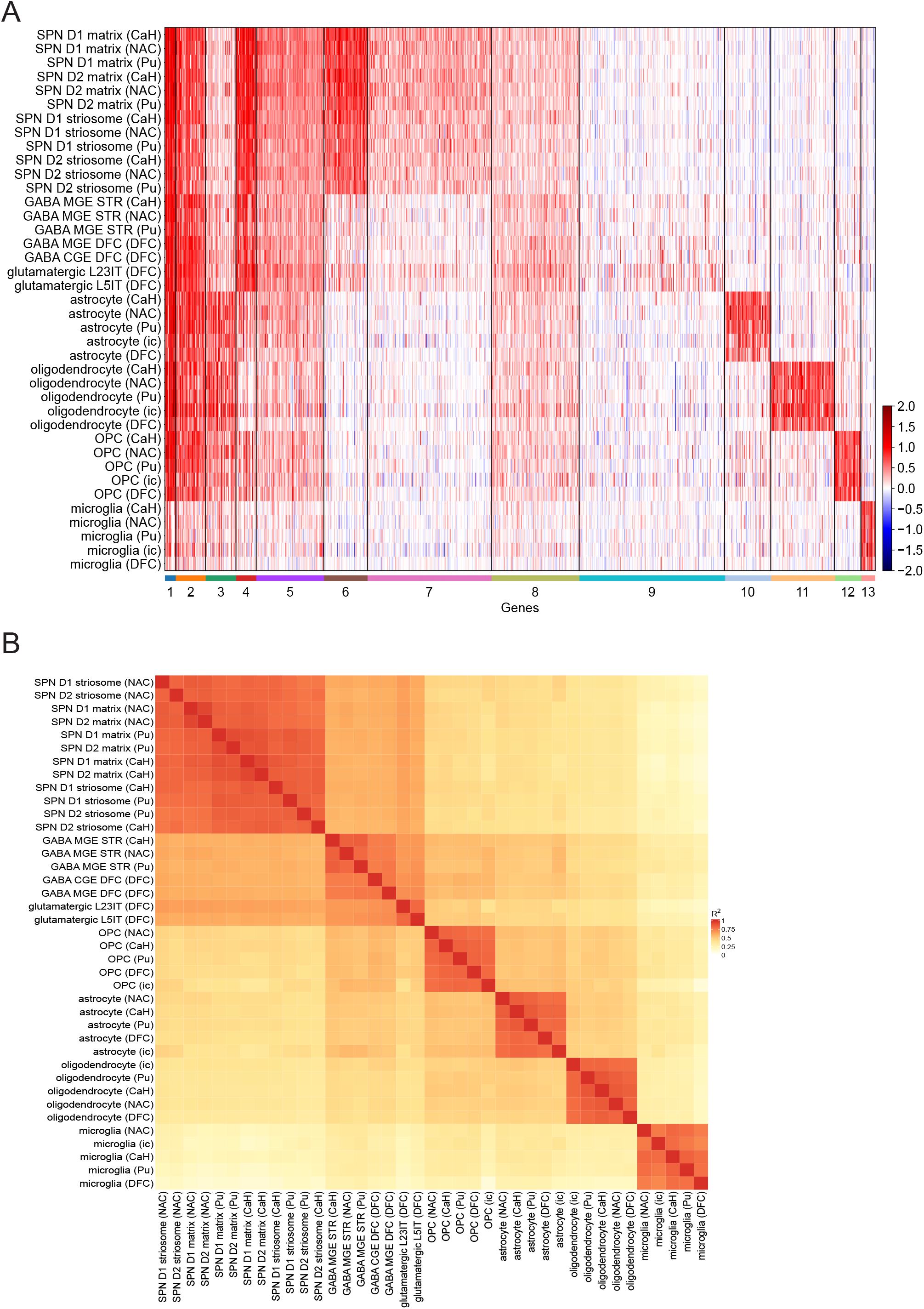
Cell-type specificity of expression QTLs. Extension of Figure 6 **to additional brain regions. A.** k-means clustering (k = 13) of the same 9,825 expression QTLs as shown in Figure 6B (reaching significance in either CaH, DFC, or both regions), using effect sizes estimated across five brain regions (CaH, Pu, NAC, ic, DFC). Rows represent separate region-specific analyses of eQTLs in each cell type, and columns represent these eQTLs (gene- SNP pairs) grouped by cluster. Shades of red and blue indicate standardized effect sizes (inverse-normalized log2- transformed expression), with red indicating the predominant direction of effect and blue indicating opposite-direction effects. **B.** Pairwise correlations (among cell types sampled in each brain region) of genome-wide sets of eQTL effects (corresponding to the rows in **A**). For each cell-type pair, Spearman’s ρ^2^ was calculated using eQTLs significant in at least one of the two cell types. Rows and columns were hierarchically clustered.

**Figure S18.**
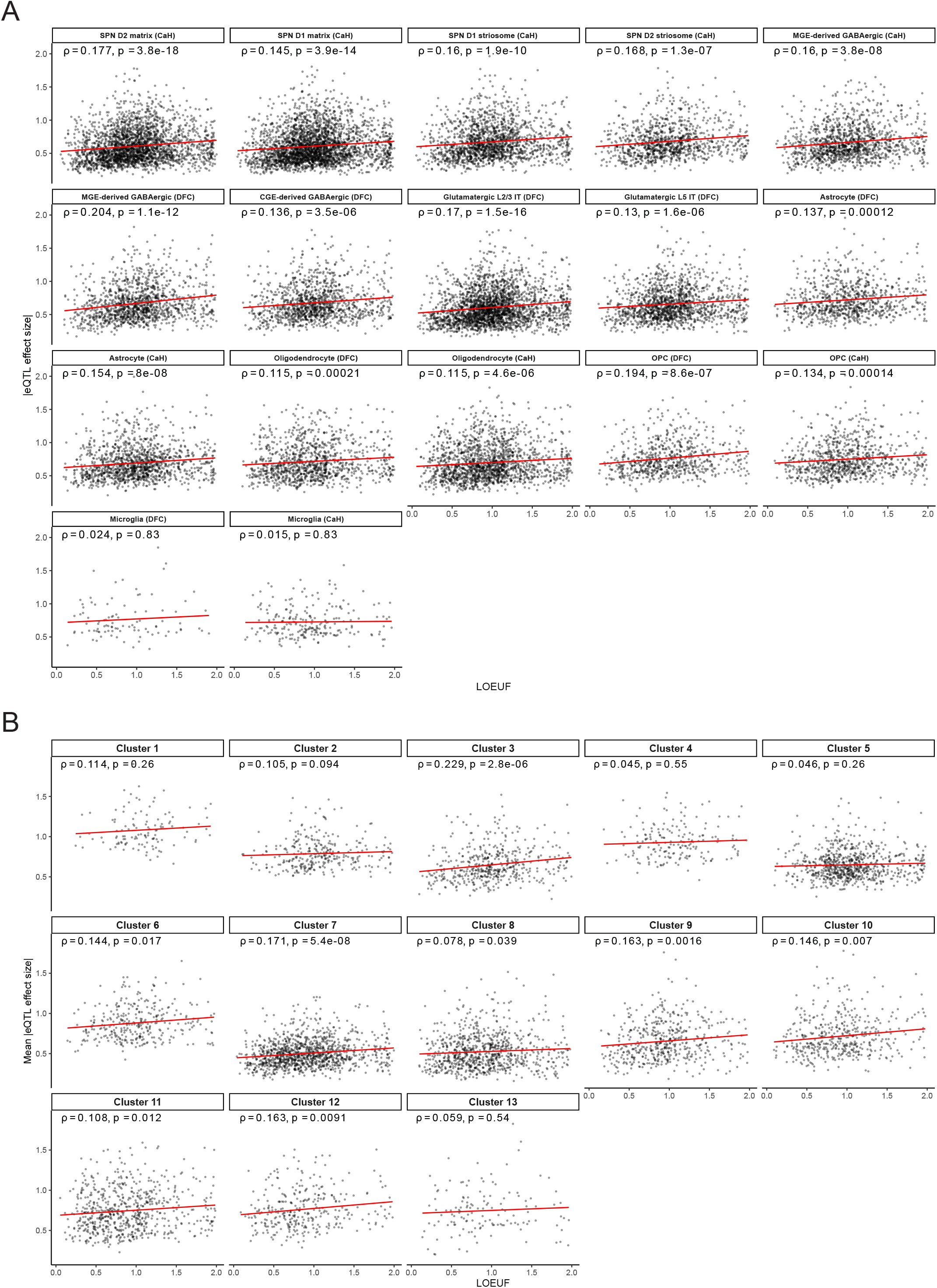
Relationship of eQTL effect sizes and gene function constraint. Related to Figure 6. eQTL effect sizes are measured as the change in gene expression per allele inherited; functional constraint is quantified by LOEUF scores (Loss-of-function Observed/Expected Upper bound Fraction). LOEUF scores of genes have been measured in earlier work^3^ and are based on the frequency (relative to chance expectation) with which genes are found to have loss-of-function mutations in human populations. Genes with low LOEUF scores have very few such mutations and are thus inferred to be under strong functional constraint (such as haploinsufficiency). **A.** Scatter plots showing the relationship of absolute eQTL effect sizes (y-axis) to LOEUF scores (x-axis) for each of 17 combinations of cell type and brain region for which eQTLs were recognized in independent analyses. **B.** Mean absolute eQTL effect sizes across statistically significant cell types for each k-means eQTL cluster (k = 13). Analyses included all protein-coding eGenes (q < 0.01) with LOEUF scores from gnomAD v4.1. Higher LOEUF indicates less constraint (greater tolerance to loss-of- function). Red lines show linear regression fits; Spearman ρ and FDR-adjusted p-values are shown per cluster. Effect sizes were measured from tensorQTL slopes (relationship of inverse-normal-transformed expression level to genotype).

**Figure S19.**
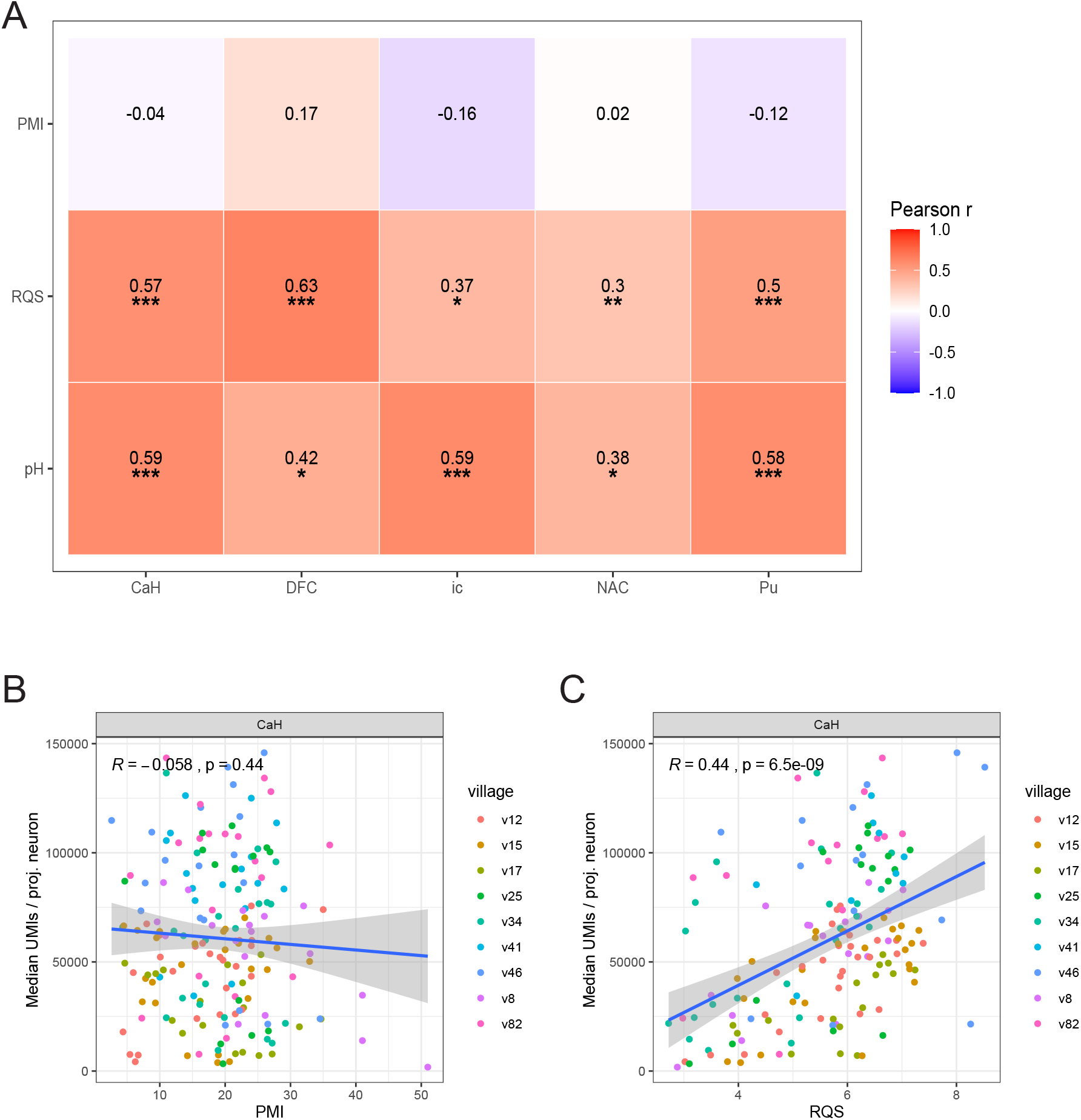
Relationships between PMI/RQS and snRNA-seq median library size. Related to **STAR Methods. A.** Correlation heatmap of tissue-quality related covariates (PMI, RQS, and pH) with village-normalized median UMIs per projection neuron in each brain region. (*) denotes BH-adjusted significance at a 0.05 threshold, (**) denotes significance below threshold / 5, and (***) denotes significance below threshold / 50. **B.** Scatter plot of PMI and median UMIs per projection neuron in CaH. **C.** Scatter plot of RQS and median UMIs per projection neuron in CaH.

**Figure S20.**
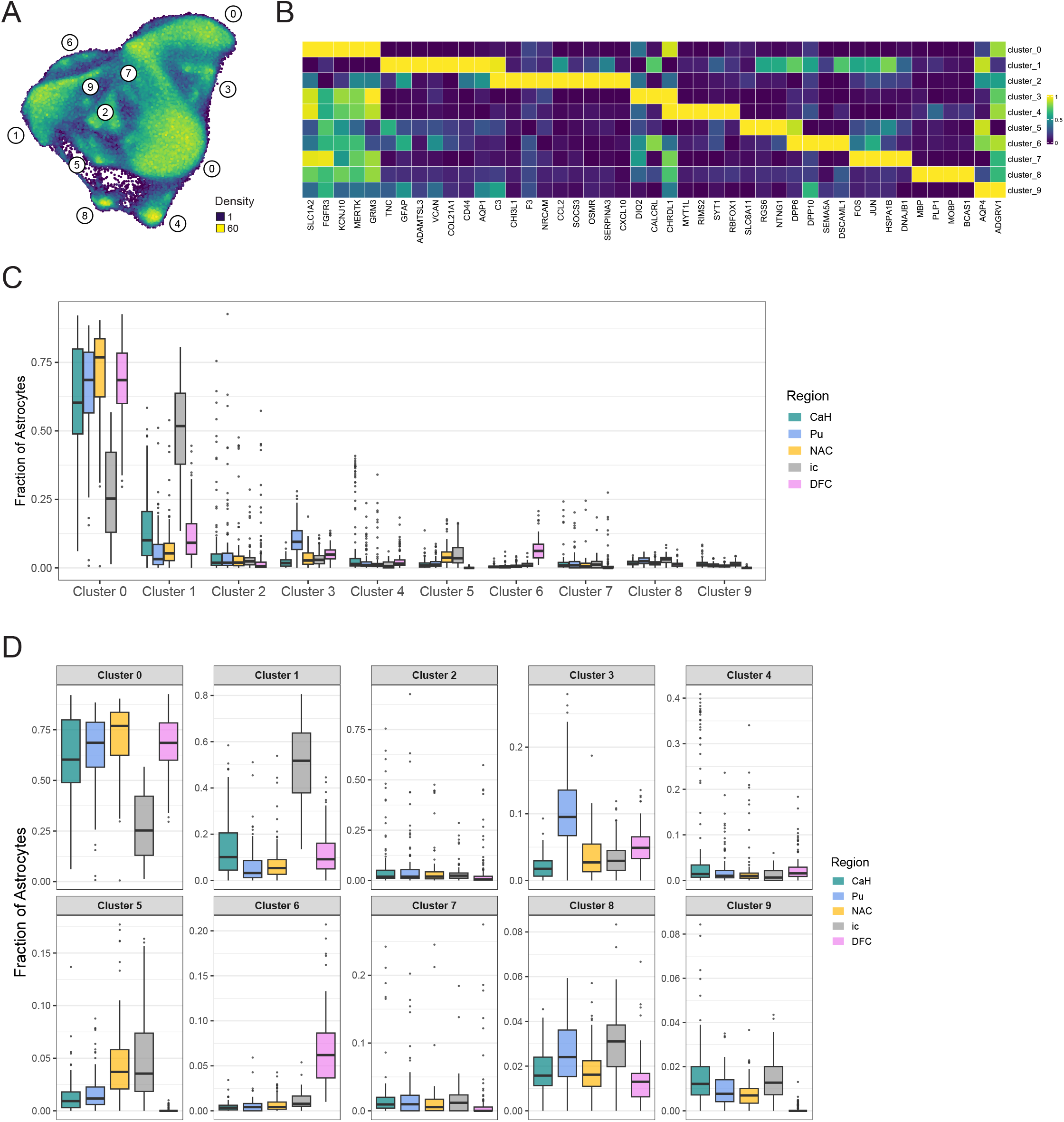
Astrocyte sub-cluster identification and abundance. Related to **STAR Methods. A.** Unsupervised clustering assignments and UMAP representation of astrocytes across striatum and cortex samples. UMAP coloring represents spatial density of nuclei. **B.** Selected marker genes (columns) for astrocyte sub-clusters (rows). Colors represent normalized expression by column (across major cell classes) such that the highest expression is 1 (bright yellow) and lowest is 0 (dark purple). **C.** Distribution of astrocyte sub-cluster compositions across regions. Gray matter regions exhibit cluster 1 astrocyte depletion and cluster 0 astrocyte enrichment relative to the internal capsule. **D.** Distribution of astrocyte sub-cluster compositions across regions (free scales).

**Table S1.** Donor stratification and analysis inclusion by diagnostic “exclusion” category, age, and sex. Related to **STAR Methods.** Donor counts by clinical/neuropathological diagnostic “exclusion” categories, age groups (< 50 years, > 50 years), and sex. Diagnostic “exclusion” categories include overdose/substance dependence, psychiatric conditions, neurodegeneration, and cases with missing clinical brain diagnosis (CBD) or neuropathology (NP). The table also indicates the inclusion of each donor group in downstream analyses.

| **Metadata exclusion category** | **Male** |  | **Female** |  | **Total** | **Analyses** |
|---|---|---|---|---|---|---|
|  | **< 50 y.o.** | **> 50 y.o.** | **< 50 y.o.** | **> 50 y.o.** |  |  |
| N/A | 28 | 62 | 13 | 44 | 147 | CTP, DE, AP, eQTL |
| overdose / dependence | 7 | 4 | 2 | 2 | 15 | DE, AP, eQTL |
| psychiatric | 0 | 2 | 0 | 2 | 4 | DE, AP, eQTL |
| neurodegeneration | 2 | 6 | 1 | 2 | 11 | DE, AP, eQTL |
| missing clinical brain diagnosis / neuropathlogy | 0 | 0 | 0 | 1 | 1 | DE, AP, eQTL |
| **Analyses** | **Donor set** | **n** | **n (after snRNA-seq-data-driven QC)** |  |  |  |
| cell-type proportions (Figure 2,3) | core | 147 | 142 |  |  |  |
| differential expression (Figure 4) | extended | 178 | 161 |  |  |  |
| age prediction (Figure 5) | extended | 178 | 161 |  |  |  |
| eQTLs (Figure 6) | extended | 178 | 161 |  |  |  |

**Table S3.**
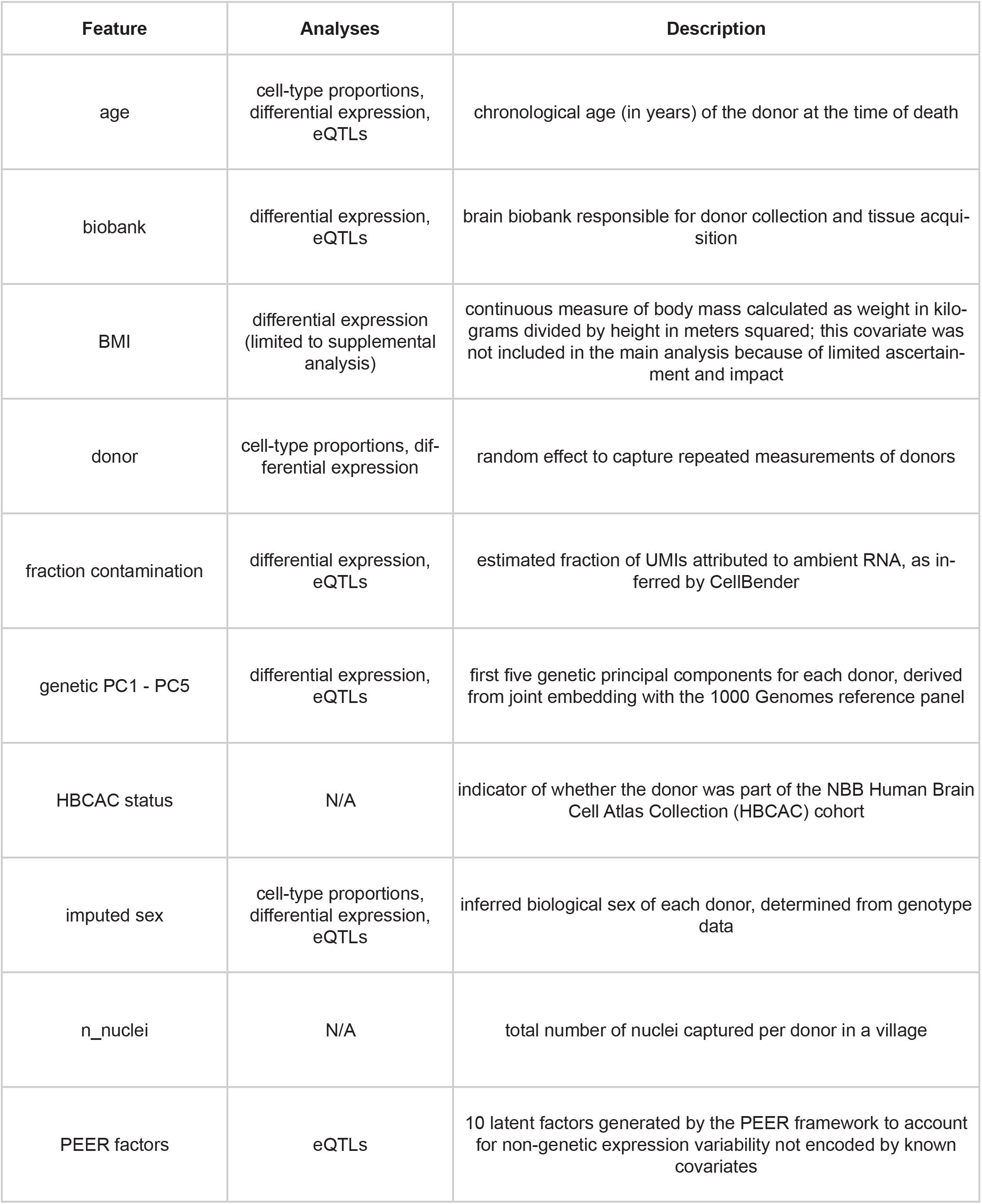
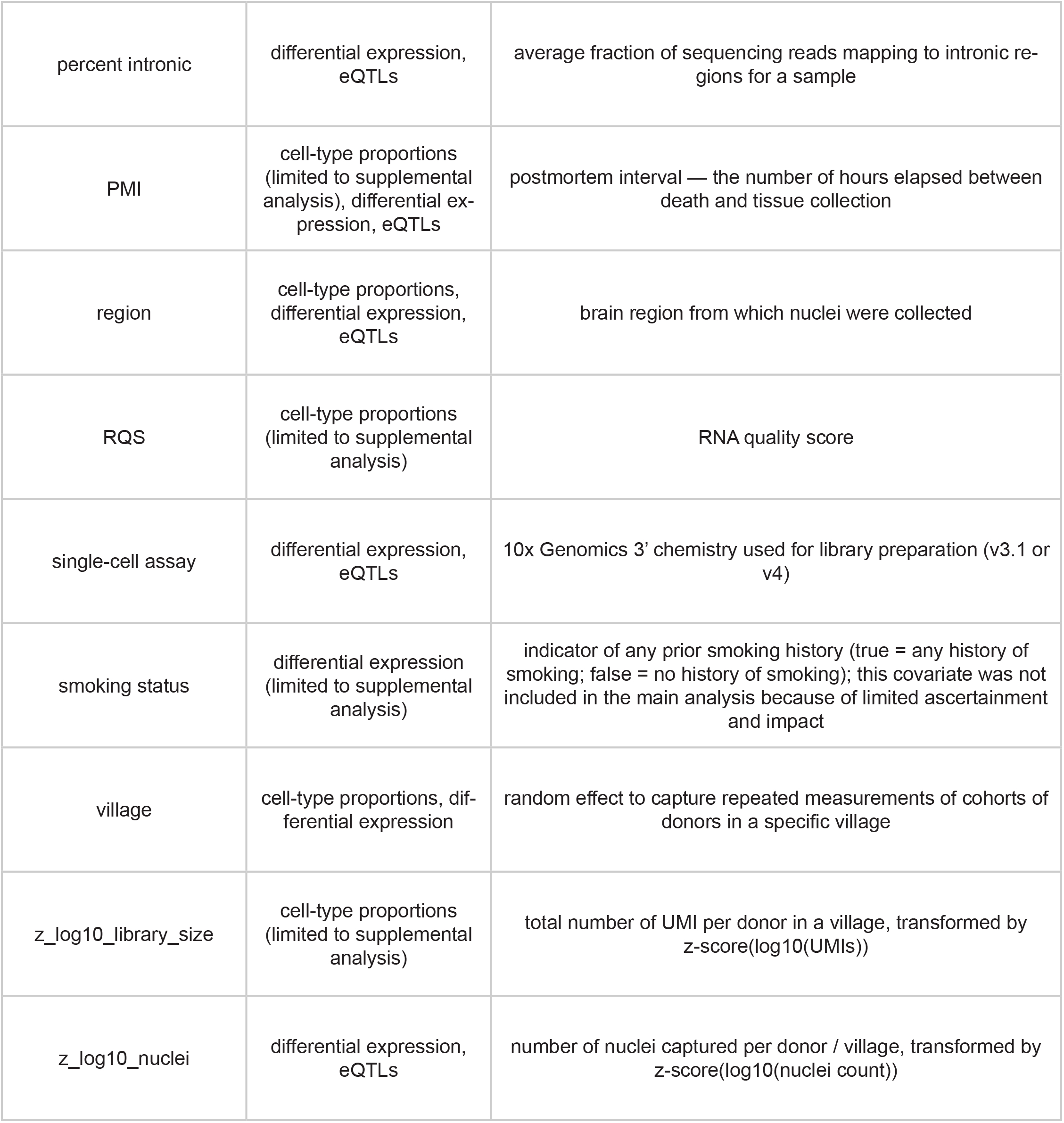
Sample covariates. Description of demographic, genetic, technical, and sequencing-quality variables.

| Feature | Analyses | Description |
|---|---|---|
| age | cell-type proportions, differential expression, eQTLs | chronological age (in years) of the donor at the time of death |
| biobank | differential expression, eQTLs | brain biobank responsible for donor collection and tissue acqui-sition |
| BMI | differential expression (limited to supplemental analysis) | continuous measure of body mass calculated as weight in kilo-grams divided by height in meters squared; this covariate was not included in the main analysis because of limited ascertain-ment and impact |
| donor | cell-type proportions, dif-ferential expression | random effect to capture repeated measurements of donors |
| fraction contamination | differential expression, eQTLs | estimated fraction of UMIs attributed to ambient RNA, as in-ferred by CellBender |
| genetic PC1 - PC5 | differential expression, eQTLs | first five genetic principal components for each donor, derived from joint embedding with the 1000 Genomes reference panel |
| HBCAC status | N/A | indicator of whether the donor was part of the NBB Human Brain Cell Atlas Collection (HBCAC) cohort |
| imputed sex | cell-type proportions, differential expression, eQTLs | inferred biological sex of each donor, determined from genotype data |
| n_nuclei | N/A | total number of nuclei captured per donor in a village |
| PEER factors | eQTLs | 10 latent factors generated by the PEER framework to account for non-genetic expression variability not encoded by known covariates |
| percent intronic | differential expression, eQTLs | average fraction of sequencing reads mapping to intronic re-gions for a sample |
| PMI | cell-type proportions (limited to supplemental analysis), differential ex-pression, eQTLs | postmortem interval — the number of hours elapsed between death and tissue collection |
| region | cell-type proportions, differential expression, eQTLs | brain region from which nuclei were collected |
| RQS | cell-type proportions (limited to supplemental analysis) | RNA quality score |
| single-cell assay | differential expression, eQTLs | 10x Genomics 3’ chemistry used for library preparation (v3.1 or v4) |
| smoking status | differential expression (limited to supplemental analysis) | indicator of any prior smoking history (true = any history of smoking; false = no history of smoking); this covariate was not included in the main analysis because of limited ascertainment and impact |
| village | cell-type proportions, dif-ferential expression | random effect to capture repeated measurements of cohorts of donors in a specific village |
| z_log10_library_size | cell-type proportions (limited to supplemental analysis) | total number of UMI per donor in a village, transformed by z-score(log10(UMIs)) |
| z_log10_nuclei | differential expression, eQTLs | number of nuclei captured per donor / village, transformed by z-score(log10(nuclei count)) |

**Table S6.** External validation of differential expression and eQTL effects. Related to **Figure 4**, **Figure 6**. Compari- son of age- and sex-associated differential expression effects and eQTL effects identified in this study with corresponding effects in Ling *et al.*, including sign concordance and, for differential expression, effect-size correlations by cell type.

| Analysis | Gene set | Cell type | n compared | % sign agreement | Effect-size correlation |
|---|---|---|---|---|---|
| differential expression, sex | XY | glut_L23_IT | 47 | 93.6 | 0.98 |
| differential expression, sex | XY | glut_L4_IT | 42 | 100.0 | 0.99 |
| differential expression, sex | XY | glut_L5_IT | 40 | 100.0 | 0.99 |
| differential expression, sex | XY | glut_L6_IT | 41 | 100.0 | 0.99 |
| differential expression, sex | XY | GABA_LAMP5 | 38 | 100.0 | 0.99 |
| differential expression, sex | XY | GABA_PVALB | 41 | 97.6 | 0.99 |
| differential expression, sex | XY | GABA_SST | 36 | 100.0 | 0.98 |
| differential expression, sex | XY | GABA_VIP | 45 | 97.8 | 0.98 |
| differential expression, sex | XY | astrocyte | 41 | 97.6 | 0.97 |
| differential expression, sex | XY | OPC | 41 | 100.0 | 0.97 |
| differential expression, sex | XY | oligodendrocyte | 55 | 94.5 | 0.97 |
| differential expression, sex | XY | microglia | 29 | 100.0 | 0.96 |
| differential expression, sex | autosomes | glut_L23_IT | 67 | 89.6 | 0.92 |
| differential expression, sex | autosomes | glut_L4_IT | 36 | 88.9 | 0.76 |
| differential expression, sex | autosomes | glut_L5_IT | 24 | 87.5 | 0.88 |
| differential expression, sex | autosomes | glut_L6_IT | 40 | 85.0 | 0.90 |
| differential expression, sex | autosomes | GABA_LAMP5 | 20 | 100.0 | 0.93 |
| differential expression, sex | autosomes | GABA_PVALB | 35 | 88.6 | 0.90 |
| differential expression, sex | autosomes | GABA_VIP | 30 | 90.0 | 0.92 |
| differential expression, sex | autosomes | astrocyte | 121 | 86.0 | 0.82 |
| differential expression, sex | autosomes | oligodendrocyte | 44 | 59.1 | 0.49 |
| differential expression, age | all | glut_L23_IT | 2195 | 91.1 | 0.83 |
| differential expression, age | all | glut_L4_IT | 1429 | 89.9 | 0.80 |
| differential expression, age | all | glut_L5_IT | 1465 | 88.9 | 0.80 |
| differential expression, age | all | glut_L6_IT | 1112 | 93.3 | 0.84 |
| differential expression, age | all | GABA_PVALB | 59 | 96.6 | 0.84 |
| differential expression, age | all | GABA_SST | 258 | 89.5 | 0.79 |
| differential expression, age | all | astrocyte | 1384 | 88.2 | 0.78 |
| differential expression, age | all | OPC | 1328 | 87.5 | 0.76 |
| differential expression, age | all | oligodendrocyte | 352 | 79.5 | 0.61 |
| differential expression, age | all | microglia | 467 | 89.7 | 0.87 |
| eQTLs | all | glut_L23_IT | 4751 | 91.9 | N/A |
| eQTLs | all | glut_L4_IT | 1733 | 94.1 | N/A |
| eQTLs | all | glut_L5_IT | 2884 | 92.9 | N/A |
| eQTLs | all | glut_L6_IT | 1724 | 93.3 | N/A |
| eQTLs | all | GABA_LAMP5 | 1386 | 94.9 | N/A |
| eQTLs | all | GABA_PVALB | 2207 | 93.4 | N/A |
| eQTLs | all | GABA_SNCG | 335 | 97.9 | N/A |
| eQTLs | all | GABA_SST | 914 | 94.6 | N/A |
| eQTLs | all | GABA_VIP | 1509 | 95.2 | N/A |
| eQTLs | all | astrocyte | 1722 | 95.1 | N/A |
| eQTLs | all | OPC | 1343 | 95.6 | N/A |
| eQTLs | all | oligodendrocyte | 2035 | 92.7 | N/A |
| eQTLs | all | microglia | 202 | 97.0 | N/A |

**Table S9.** Union of eGenes across DFC and CaH by cell type. Related to Figure 6. Counts of unique eGenes identified per cell type, defined as the union of eGenes detected in dorsolateral prefrontal cortex (DFC) and caudate head (CaH).

| **Cell type** | **Union of eGenes detected in DFC and CaH** |
|---|---|
| SPN_D1_matrix | 4520 |
| SPN_D2_matrix | 4088 |
| glut_L23_IT | 3973 |
| oligodendrocyte | 2841 |
| SPN_D1_striosome | 2665 |
| astrocyte | 2412 |
| glut_L5_IT | 2296 |
| GABA_MGE_DFC | 2016 |
| GABA_MGE_STR | 1983 |
| GABA_CGE_DFC | 1966 |
| SPN_D2_striosome | 1659 |
| OPC | 1569 |
| microglia | 382 |

**Table S10.**
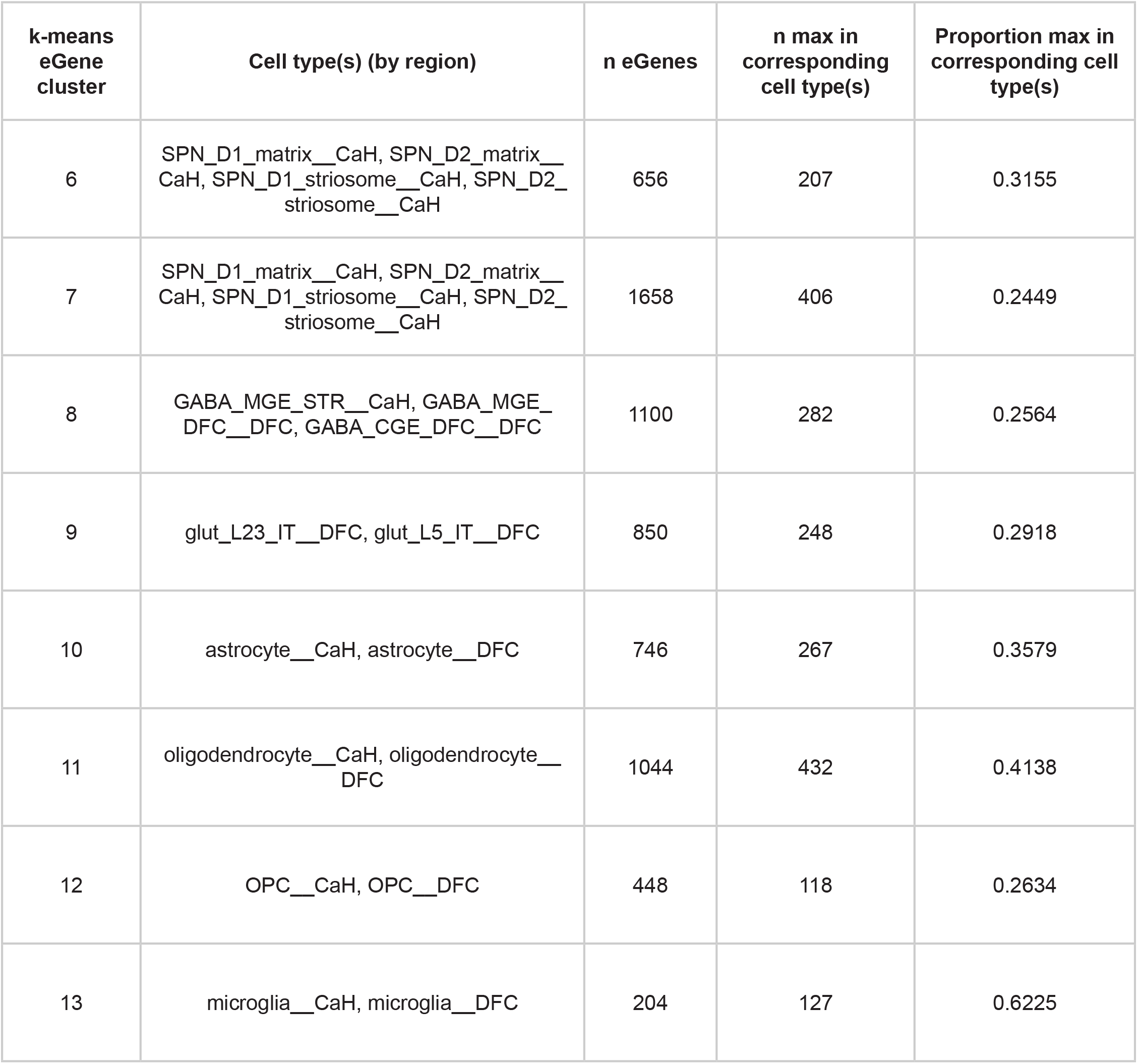
Fraction of cell-type-specific eGenes with maximum expression in corresponding cell type(s). Related to **Figure 6**. For each eGene cluster (k-means; see **Figure 6**), the expected cell type(s) (by region), the number of eGenes with nonzero median expression across cell types, the number of eGenes whose highest median expression occurs in one of the corresponding cell types, and the proportion of eGenes whose maximum median expression falls in one of the expected cell type(s) (see **Methods**).

**Table S11.** Genotype-by-cell-type interaction validation of cell-type-specific eQTL patterns. Related to **Figure 6**. For each cell-type-specific k-means cluster, counts of cluster-associated eGenes that were expressed only in the target cell-type group, testable across cell types, or supported by a significant genotype-by-group interaction. Results are shown for both the full and downsampled datasets; percentages indicate the fraction of eGenes expressed only in the target group and the fraction of cross-cell-type testable eGenes supported by a significant interaction.

| Dataset type | Cell type / cluster | Cluster-associated eGenes | Expression-private eGenes | Cross-cell-type testable eGenes | Significant interaction eGenes | Expression-private % | Interaction-supported % |
|---|---|---|---|---|---|---|---|
| full dataset | SPN (strong effect cluster) | 656 | 69 | 587 | 541 | 10.5 | 92.2 |
| full dataset | SPN (moderate effect cluster) | 1658 | 128 | 1530 | 1103 | 7.7 | 72.1 |
| full dataset | Glutamatergic L23 + Glutama-tergic L5 | 850 | 104 | 746 | 575 | 12.2 | 77.1 |
| full dataset | astrocyte | 743 | 67 | 676 | 580 | 9 | 85.8 |
| full dataset | OPC | 446 | 27 | 419 | 366 | 6.1 | 87.4 |
| full dataset | oligodendrocyte | 1042 | 142 | 900 | 817 | 13.6 | 90.8 |
| full dataset | microglia | 204 | 55 | 149 | 143 | 27 | 96 |
| downsampled dataset | SPN (strong effect cluster) | 419 | 36 | 383 | 361 | 8.6 | 94.3 |
| downsampled dataset | SPN (moderate effect cluster) | 1315 | 111 | 1204 | 943 | 8.4 | 78.3 |
| downsampled dataset | Glutamatergic L23 + Glutama-tergic L5 | 630 | 63 | 567 | 453 | 10 | 79.9 |
| downsampled dataset | astrocyte | 502 | 41 | 461 | 400 | 8.2 | 86.8 |
| downsampled dataset | OPC | 431 | 24 | 407 | 367 | 5.6 | 90.2 |
| downsampled dataset | oligodendrocyte | 464 | 47 | 417 | 364 | 10.1 | 89.4 |
| downsampled dataset | microglia | 186 | 51 | 135 | 131 | 27.4 | 97 |

**Table S12.** Donor-level RNAscope and snRNA-seq cell-type proportion measurements. Donor age and sex, brain region, RNAscope probe, RNAscope classifier assignments, and cell-type proportions measured by RNAscope and snRNA-seq are reported for each donor included in the RNAscope analyses. *PDGFRA* was used to identify OPCs in caudate and putamen, and *SLC1A2* was used to identify astrocytes in caudate. RNAscope classifier assignments indicate whether sections were analyzed using the shared (“universal”) classifier or a separately trained, section-specific (“individual”) classifier (see **Methods**).

| **Donor** | **Age** | **Sex** | **CaH OPCs** |  |  |
|  |  |  | **snRNA-seq proportion** | **RNAscope proportion** | **RNAscope classifier** |
|---|---|---|---|---|---|
| MD9047 | 35 | F | 7.23 | 3.96 | individual |
| UMBEB23073 | 36 | M | 8.3 | 5.72 | individual |
| MD9162 | 45 | M | 8.07 | 6.8 | individual |
| PT13967 | 48 | M | 3.48 | 3.3 | universal |
| MD6592 | 53 | F | 5.18 | 3.67 | individual |
| UMBEB23033 | 64 | M | 3.92 | 3.3 | universal |
| UMBEB23076 | 66 | M | 2.17 | N/A | N/A |
| MD9261 | 69 | M | 3.52 | 3.8 | universal |
| MS986638 | 83 | F | 3.91 | 4.3 | universal |
| MS587476 | 89 | M | 3.19 | 3.4 | universal |
| MS53636 | 89 | M | 2.66 | 2.1 | universal |
| MLS04249 | 90 | F | 1.74 | 1.9 | universal |
| **Donor** | **Age** | **Sex** | **Pu OPCs** |  |  |
|  |  |  | **snRNA-seq proportion** | **RNAscope proportion** | **RNAscope classifier** |
| MD9047 | 35 | F | 7.12 | N/A | N/A |
| UMBEB23073 | 36 | M | 7.44 | N/A | N/A |
| MD9162 | 45 | M | 7.37 | 6.9 | individual |
| PT13967 | 48 | M | 3.97 | 5.2 | individual |
| MD6592 | 53 | F | 5.86 | 5.3 | individual |
| UMBEB23033 | 64 | M | 2.35 | N/A | N/A |
| UMBEB23076 | 66 | M | 3.58 | N/A | N/A |
| MD9261 | 69 | M | 2.96 | 3.78 | individual |
| MS986638 | 83 | F | 5.78 | 4.1 | individual |
| MS587476 | 89 | M | 3.69 | 3.18 | individual |
| MS53636 | 89 | M | 1.39 | 1.8 | individual |
| MLS04249 | 90 | F | 3.16 | 2.25 | individual |
| **Donor** | **Age** | **Sex** | **CaH astrocytes** |  |  |
|  |  |  | **snRNA-seq proportion** | **RNAscope proportion** | **RNAscope classifier** |
| MD9047 | 35 | F | 18.63 | 14.4 | individual |
| UMBEB23073 | 36 | M | 9.83 | 8.77 | universal |
| MD9162 | 45 | M | 18.66 | 13.5 | individual |
| PT13967 | 48 | M | 7.39 | 9.73 | universal |
| MD6592 | 53 | F | 12.93 | N/A | N/A |
| UMBEB23033 | 64 | M | 6.59 | 14.5 | universal |
| UMBEB23076 | 66 | M | 5.1 | 4.63 | universal |
| MD9261 | 69 | M | 3.55 | 8.44 | individual |
| MS986638 | 83 | F | 9.58 | 8.6 | universal |
| MS587476 | 89 | M | 7.14 | 7.55 | universal |
| MS53636 | 89 | M | 2.89 | 5.98 | universal |
| MLS04249 | 90 | F | 15.45 | 15.8 | individual |

**Table S14.** Age-residual correlations before and after covariate adjustment. Related to **Figure 5**. Pairwise Pearson correlations among bias-corrected age-prediction residuals across cell types before and after adjustment for postmortem interval (PMI), RNA quality score (RQS), and log_10_ library size. The number of donors and change in correlation (adjusted minus unadjusted) are reported for each cell-type pair, with both correlations calculated using the same donors.

| **Pair** | **n donors** | **Correlation unadjusted** | **Correlation adjusted** | **Correlation change** |
|---|---|---|---|---|
| astrocyte vs microglia | 118 | 0.351 | 0.315 | -0.035 |
| astrocyte vs OPC | 119 | 0.257 | 0.266 | 0.009 |
| astrocyte vs SPN D1 | 114 | 0.398 | 0.349 | -0.048 |
| microglia vs OPC | 121 | 0.398 | 0.387 | -0.011 |
| microglia vs SPN D1 | 122 | 0.368 | 0.321 | -0.047 |
| OPC vs SPN D1 | 117 | 0.333 | 0.32 | -0.013 |

**Table S15.** Age-prediction accuracy with all genes and DE-selected genes. Related to Figure 5. Mean absolute error (MAE, in years) for age-prediction models using all expressed genes or restricted to genes significantly associated with age (FDR ≤ 0.05), reported by cell type and brain region. The MAE difference is calculated as the all-gene model MAE minus the DE-selected model MAE; positive values indicate improved accuracy with DE feature selection.

| Cell type | Region | MAE DE priors | MAE all genes | MAE difference |
|---|---|---|---|---|
| microglia | ic | 6.525 | 8.152 | 1.627 |
| microglia | NAC | 6.82 | 8.241 | 1.422 |
| microglia | Pu | 6.823 | 8.223 | 1.4 |
| GABA_TAC3-PLPP4 | Pu | 8.314 | 9.573 | 1.259 |
| microglia | DFC | 7.215 | 8.435 | 1.22 |
| oligodendrocyte | ic | 5.962 | 7.094 | 1.133 |
| GABA_PTHLH-PVALB | Pu | 8.413 | 9.132 | 0.719 |
| GABA_PVALB | DFC | 7.423 | 8.123 | 0.699 |
| microglia | CaH | 7.343 | 8.002 | 0.659 |
| astrocyte | Pu | 6.388 | 7.005 | 0.617 |
| astrocyte | NAC | 6.07 | 6.596 | 0.525 |
| glut_L6_IT | DFC | 5.664 | 6.175 | 0.511 |
| SPN_D2_striosome | NAC | 6.512 | 6.974 | 0.462 |
| oligodendrocyte | NAC | 5.604 | 6.059 | 0.455 |
| OPC | Pu | 6.832 | 7.285 | 0.452 |
| oligodendrocyte | Pu | 5.907 | 6.359 | 0.452 |
| OPC | ic | 7.093 | 7.527 | 0.434 |
| OPC | DFC | 5.822 | 6.221 | 0.399 |
| SPN_D2_matrix | Pu | 5.406 | 5.79 | 0.385 |
| oligodendrocyte | CaH | 5.909 | 6.243 | 0.334 |
| SPN_D1_matrix | Pu | 5.099 | 5.428 | 0.33 |
| glut_L5_IT | DFC | 5.111 | 5.348 | 0.237 |
| SPN_D2_striosome | Pu | 5.85 | 6.081 | 0.231 |
| SPN_D1_matrix | CaH | 5.219 | 5.413 | 0.194 |
| SPN_D2_striosome | CaH | 5.843 | 5.989 | 0.146 |
| SPN_D1_striosome | Pu | 5.587 | 5.636 | 0.049 |
| SPN_D2_matrix | CaH | 5.146 | 5.182 | 0.036 |
| glut_L4_IT | DFC | 5.057 | 5.082 | 0.025 |
| astrocyte | ic | 8.019 | 8.028 | 0.009 |
| astrocyte | DFC | 6.708 | 6.669 | -0.039 |
| astrocyte | CaH | 6.251 | 6.194 | -0.056 |
| SPN_D1_matrix | NAC | 4.984 | 4.897 | -0.087 |
| GABA_PTHLH-PVALB | CaH | 8.223 | 8.116 | -0.107 |
| SPN_D1_striosome | NAC | 6.065 | 5.939 | -0.126 |
| glut_L23_IT | DFC | 5.096 | 4.926 | -0.17 |
| SPN_D1_striosome | CaH | 5.953 | 5.644 | -0.308 |
| SPN_D2_matrix | NAC | 5.091 | 4.726 | -0.365 |
| oligodendrocyte | DFC | 6.335 | 5.86 | -0.475 |
| OPC | CaH | 6.649 | 6.128 | -0.521 |
| OPC | NAC | 7.59 | 6.947 | -0.643 |
| GABA_SST | DFC | 8.357 | 7.654 | -0.703 |
| GABA_VIP | DFC | 9.044 | 8.218 | -0.826 |
| GABA_TAC3-PLPP4 | NAC | 9.901 | 8.982 | -0.919 |
| GABA_TAC3-PLPP4 | CaH | 8.249 | 7.239 | -1.009 |
| GABA_PTHLH-PVALB | NAC | 10.304 | 8.7 | -1.604 |
| GABA_LAMP5 | DFC | N/A | 8.774 | N/A |

